# Basal p53 maintains a distinct transcriptional program from irradiated p53 in tissue, including tumor suppressors

**DOI:** 10.1101/2024.07.11.601367

**Authors:** Royce W. Zhou, Lois Silverman, James J. Manfredi, Ramon E. Parsons

## Abstract

The significance of p53’s primary and secondary tumor suppressor programs cannot be overstated. A context- and stress-dependent transcription factor, p53 accumulates to mount its most well-characterized programs in response to a variety of stressors, most notably DNA damage. As cells and tissues never exist in a complete absence of stress, a small amount of p53 exists in cells under physiologic stress, detectable by chromatin immunoprecipitation and sequencing, termed basal p53. Recently, we and others have shown that basal p53 is sufficient to regulate tumor suppressor function. Furthermore, others have suggested the possibility that p53 accumulation in response to experimental stress may be dispensable for its tumor suppression. We previously showed basal p53 occupancy and regulation of known tumor suppressor genes, including *PTEN* and *PHLDA3,* in non-transformed breast cells, but this study was limited by experimental stress inherent to cell culture. Given the lack of global characterization of the basal p53 landscape under non-malignant physiologic stress *in vivo,* we utilized a multi-omics approach to define the murine basal p53 epigenome and its transcriptional program in various normal murine tissues. In this study, we observed basal p53 binding to cis regions of multiple tumor suppressor genes in different tissues, of which some showed p53-dependent regulation of their expression, including *Phlda3, Bbc3, Xaf1,* and itself. Furthermore, the vast majority of basal p53 target genes were not induced upon irradiation, suggesting basal p53 operates a transcriptional program that is largely distinct from its DNA damage response. Similarly, the basal p53 target gene repertoire is unique to each tissue type.

## Introduction

p53 undergoes constant MDM2-mediated ubiquitination and proteasomal degradation under basal conditions, defined as the absence of experimental or malignancy-induced stress[1–3]. Whether low levels of basal p53 protein plays a role in tumor suppression remains incompletely understood. A recent laboratory showed an experimental mouse model of p53 unable to stabilize its protein levels is nevertheless sufficient for tumor suppression[4]. Likewise, well-described target genes regulating cell cycle arrest, apoptosis, and senescence are similarly dispensable for p53 tumor suppression, underscoring the potency of its other tumor suppressor programs[5–7]. We previously observed non-malignant breast epithelial MCF10A cells harboring WT p53 to bind and activate expression of numerous tumor suppressor genes under basal culture conditions, which is not without its own stressors especially oxidative stress[8, 9].

In this study, we expand upon our previous study and characterize the basal p53 landscape and transcriptional programs in the setting of healthy mouse tissue. We find that basal p53 maintains a small transcriptional repertoire that is surprisingly distinct from irradiated p53. Whereas irradiated target genes exhibited significant overlap between tissues, the basal p53 targets were largely unique to each assayed tissue. We also find p53 regulation of DNA integrity genes such as *Xrcc2, Fignl1,*and *Cdt1* in basal liver, and *Polh* in basal spleen. We also find that basal p53 maintained the expression of several anti-viral genes, including *Mx1, Mx2,* as well as *Slfn2* a gene required for T cell mediated immunity in basal colon.

While irradiated targets are not activated in the basal setting, they are nevertheless bound by p53. This suggests basal p53 may be priming these genes for activation upon subsequent stress.

## Results

### The basal p53 landscape *in vivo*

To investigate the basal p53 transcriptional program in healthy normal tissue under physiologic conditions, we optimized a double cross-link ChIP-seq protocol towards the detection of basal p53 in various mouse tissues[10, 11]. We initially performed p53 ChIP-seq, using the CM5 antibody, on freshly isolated colon epithelium from *Trp53^+/+^* mice because p53 is known to act a a tumor suppressor in this tissue (Figure 1A). Merging two biological replicates, we called a total of 1126 high-confidence p53 peaks in murine non-neoplastic colon epithelium under normal homeostasis. Given the technical challenges of performing ChIP-seq on tissue, especially p53 which exhibits low protein levels under basal conditions, we first assessed our generated datasets for stringent quality control. Using the HOMER algorithm, we identified the p53 consensus binding site as the most significantly enriched transcriptional factors within called peaks, indicating fidelity of our generated ChIP-seq datasets towards p53 (Figure 1B). p53 occupancy was observed to be most prevalent at promoters or the first intron of target genes as previously described[12]. We thus assigned putative target genes to each p53 peak based on the nearest transcriptional start site using the HOMER algorithm. We observed basal p53 binding to well-characterized p53 target genes, including *Cdkn1a, Bax,* and *Bbc3* (Figure 1C).

**Figure 1.**
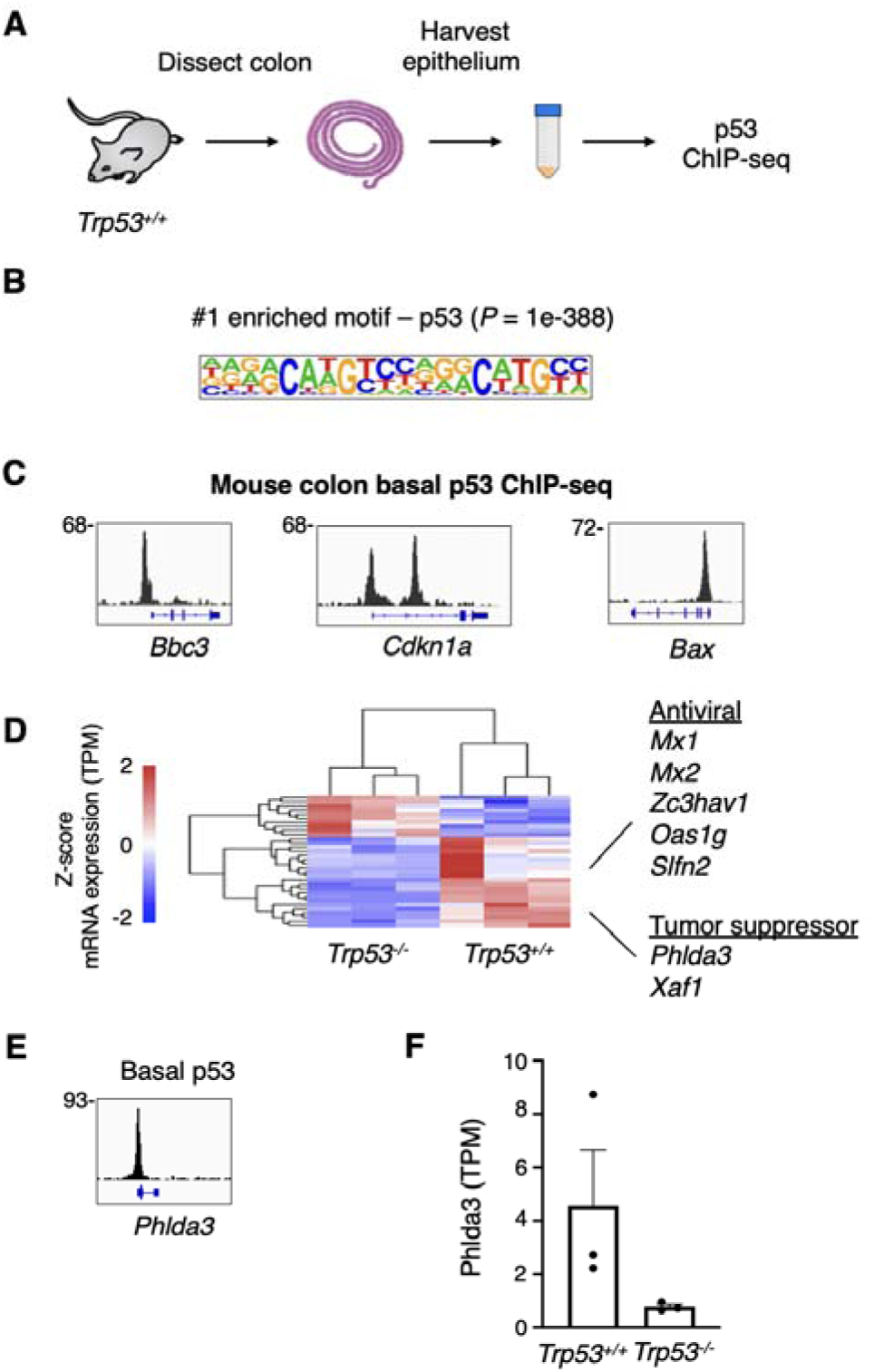
An optimized ChIP-seq protocol for basal p53 in homeostatic murine tissues. (A) Experiment overview. (B) HOMER *de novo* motif analysis performed on a total of 1,126 called basal p53 peaks from murine colon epithelium. (C) p53 ChIP-seq tracks at well-established p53 target genes. (D) Heatmap of a basal p53 bona fide target gene signature in murine colon epithelium. Gene expression via RNA-seq (TPM normalized as Z-score) between *n* = 3 WT and *n* = 3 null mice. (E) p53 ChIP-seq tracks at *Phlda3*, a known p53 target tumor suppressor in murine colon epithelium. (F) Gene expression of *Phlda3* in physiologic murine colon epithelium via RNA-seq between *n* = 3 WT and *n* = 3 null mice.

We next wished to characterize whether p53 occupancy at promoters and other regulatory elements of these genes affect mRNA expression under basal conditions. Towards this end, we performed RNA-sequencing (RNA-seq) on colon epithelium between *Trp53^+/+^* and *Trp53^-/-^* mice, hereafter referred to at WT and null respectively. Despite identifying 1,126 basal colon p53 peaks, only 32 (2.9%) were linked to genes that were significantly differentially expressed between WT and null colon. Although inaccurate peak-to-gene assignment may contribute to this observation, it strongly suggests basal p53 occupancy largely does not affect gene expression. We did not observe any difference in mRNA expression at well-characterized p53 target genes *Cdkn1a, Bax,* and *Bbc3* under between *Trp53^+/+^* and *Trp53^-/-^* mice despite p53 occupancy under basal conditions (Figure 1C; Supplementary Figure S1A-C).

### Basal p53 target genes in colon

Among the 32 genes with basal p53 occupancy and significant differential expression (*P_adj_* < 0.05) between WT and null colon epithelium, 22 genes exhibited higher expression in the setting of WT p53 (Figure 1D). p53 down-regulation of genes tended to be mild (median log_2_ fold change −0.49) compared to activated genes (median log_2_ fold change 2.21).

We shortlisted genes that were upregulated >1 log_2_ fold change as a threshold and identified fifteen p53 bona fide target genes with this criteria, most of which are novel (Supplementary Figure 1D-E; Supplementary Figure 2). Eleven out of fifteen genes were found to have a p53 consensus binding sequence using TRAP *de novo* motif discovery of DNA sequences underlying their basal p53 peaks (Supplementary Figure 2). These include previously described p53-regulated tumor suppressor genes, including *Phlda3* which encodes a competitive inhibitor of Akt (Figure 1E-F)[13]. This corroborated our previously observed basal p53 regulation of *Phlda3* in cultured MCF10A cells[9]. Notably, null mice did not express *Phlda3* in their colon epithelium (<1 TPM by RNA-seq), suggesting that basal p53 may be critical for expression of this tumor suppressor in this setting. We also observe basal p53 regulation of *Xaf1,* a pro-apoptotic zinc finger that competes with Mdm2 to stabilize p53 protein levels (Supplementary Figure 2B)[14–16]. This suggests that basal p53 maintain this positive-feedback loop in physiologic colon epithelium. Taken together, we observe that p53 activation of tumor suppressors occurs even under physiologic tissue conditions, as an extension beyond our previous study of epithelial cells in replete media, and identify novel p53 targets as candidates for further functional studies[9].

Other p53 target genes in basal colon epithelium notably enriched for genes involved in interferon and anti-viral response, including *Mx1, Mx2,* and *Oas1g* (Figure 1D, Supplementary Figure 2). Among the basal p53 targets not previously described in the literature, we identified *Slfn2* as a novel bona fide target, encoding a protein recently identified as essential for T cell-mediated immunity by protecting tRNAs from oxidative stress induced ribonuclease cleavage (Supplementary Figure 2N)[17, 18]. Another target, *Lgals3bp,* was previously described to inhibit growth of CRC cell lines as xenografts and its expression positively prognosticates overall survival of CRC patients (Supplementary Figure 2E)[19, 20]. *Lgals3bp,* which encodes a protein that regulates centriole biogenesis and centrosome hypertrophy, is also known to induce interferon responses and pro-inflammatory cytokines[21, 22].

### Basal p53 targets in other tissues

p53 is known to exhibit context-dependent activity[23]. We next assessed whether the basal p53 transcriptional program, including its regulation of tumor suppressors, differs across murine tissue types. Towards this end, we expanded to perform p53 ChIP-seq on basal spleen, thymus, and liver, all in parallel with each tissue type in duplicate. To allow for optimal direct comparison of p53 ChIP-seq signal, we excluded the basal colon p53 ChIP-seq datasets generated earlier for pooled analyses to exclude batch effect.

Basal p53 in murine thymus, liver, and spleen met quality control criteria with high signal-to-background ratios at established p53 target genes. Among all replicates from all three tissue types, we called a total of 821 high confidence basal p53 peaks which significantly enriched for the p53 consensus sequence in *de novo* motif discovery (Supplementary Figure 3). Most peaks were shared between all three tissue types and 291 of these peaks (35.4%) were specific to one tissue type, with specificity defined by a mean log_2_ fold change p53 ChIP-seq signal > 1 cut-off threshold (Figure 2A). Spleen appeared to be an exception, with nearly all basal p53 peaks shared by the other two tissues (Figure 2A).

**Figure 2.**
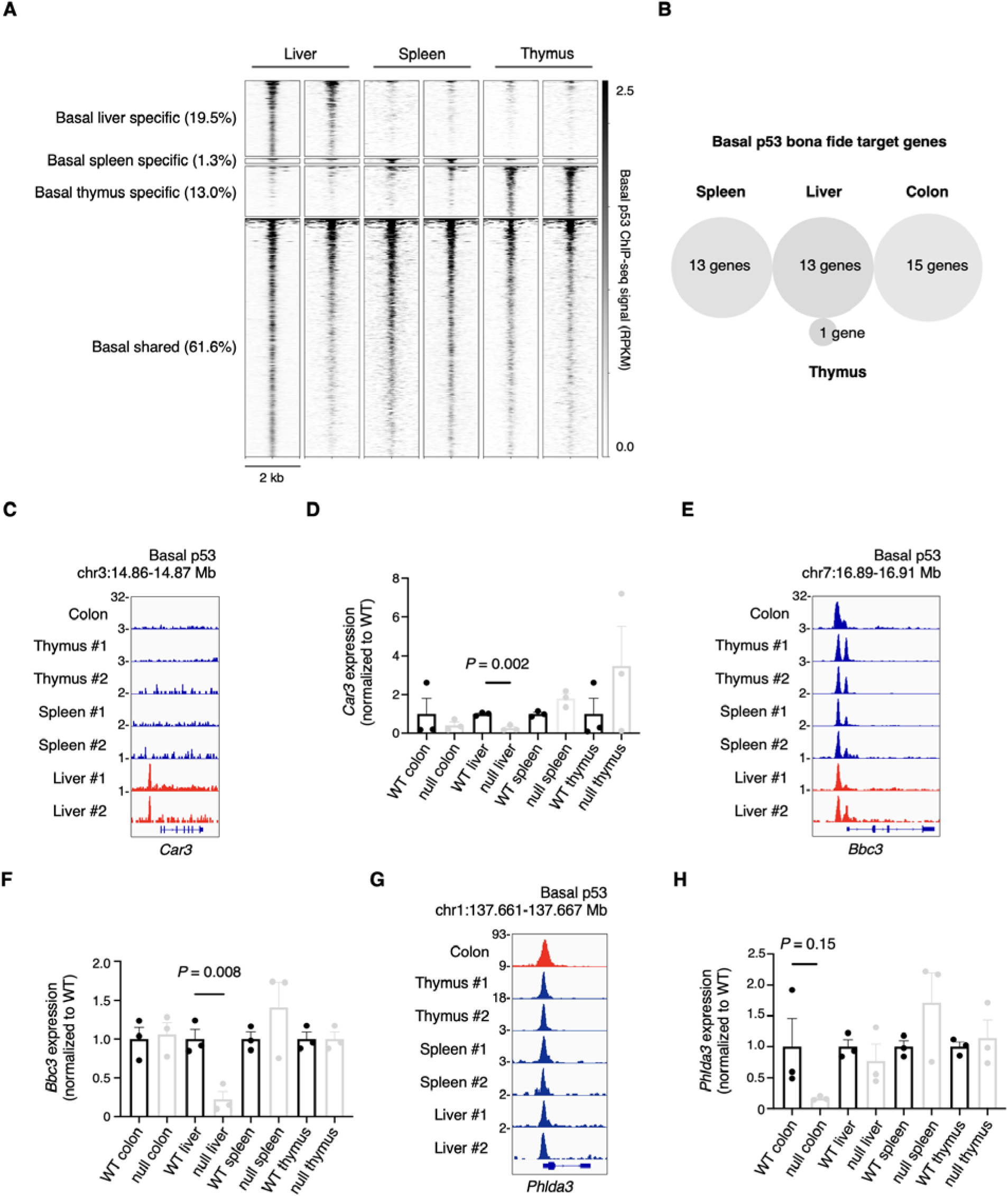
A comparison of basal p53 occupancy and *bona fide* target genes between tissue types. (A) Heatmap of basal p53 ChIP-seq signal (RPKM) in murine liver, spleen, or thymus. Tissue specific genes are those where p53 signal is > 1 log_2_ fold change greater than other tissue types. Each tissue type shown in biological replicates. (B) Venn diagram of basal p53 *bona fide* target genes, defined as those proximate to called p53 peaks and differentially > 1 log_2_ fold change expressed between p53 WT and null mice. (C) Basal p53 ChIP-seq track at the locus of *Car3.* (D) Expression of *Car3* in various mouse tissue between p53 WT (*n* = 3) or null (*n* = 3) mice; data expressed as mean with S.E.M, significance by Student’s *t* test. For each tissue type, *Car3* expression is normalized to WT. (E) Basal p53 ChIP-seq track at the locus of *Bbc3.* (F) Expression of *Bbc3* in various mouse tissue between p53 WT (*n* = 3) or null (*n* = 3) mice; data expressed as mean with S.E.M, significance by Student’s *t* test. For each tissue type, *Bbc3* expression is normalized to WT. (G) Basal p53 ChIP-seq track at the locus of *Phlda3.* (H) Expression of *Phlda3* in various mouse tissue between p53 WT (*n* = 3) or null (*n* = 3) mice; data expressed as mean with S.E.M, significance by Student’s *t* test. For each tissue type, *Phlda3* expression is normalized to WT.

To assess effects of p53 binding patterns on gene expression, we similarly performed tissue type-matched RNA-seq comparing WT and null mice. We generated basal p53 transcriptional programs for spleen, thymus, and liver utilizing the same criteria as before. Interestingly we found that none of basal p53 target genes were shared between tissue types (Figure 2B). Although some may be explained by tissue-specific p53 occupancy, such as *Car3,* most exhibit some degree of p53 occupancy in all assayed tissues (Figure 2C-D). The pro-apoptotic gene *Bbc3* (also known as Puma) is bound by p53 in all assayed tissues, but appears only dependent upon basal p53 for expression in liver, not in thymus, spleen, or colon (Figure 2E-F). Similarly, the Akt repressor gene *Phlda3* appears only maintained by basal p53 in colon, but not spleen, thymus, or liver despite basal occupancy in all these tissues (Figure 2G-H).

In liver, we identified *Bbc3, Trp53, S100a10, Ddias,* and *Ckap2* as previously described p53 targets (Supplementary Figure 4). We further propose eight other genes as *bona fide* targets, including several involved in genomic stability. We show that basal p53 binds to and maintains expression of *Xrcc2,* a gene involved in double stranded DNA break repair via homologous recombination and protects against chromosomal abnormalities (Supplementary Figure 4L-M)[24–27]. Two out of the three p53 null murine livers assayed did not meet minimal criteria for mRNA expression (TPM < 1 by RNA-seq), suggesting p53 may play an important role in its expression (Supplementary Figure 4L-M). We also identified *Fignl1,* which similarly regulates efficient homologous recombination, as a target gene of basal p53 in murine liver (Supplementary Figure 4T-U)[28]. Like *Xrcc2,* p53 null mice livers did not meet minimal criteria for mRNA expression for *Fignl1.* We observed basal p53 to also directly regulate *Cdt1,* which licenses DNA replication (Supplementary Figure 4V-W)[29, 30].

Basal p53 in normal murine spleen and thymus did not appear to directly maintain expression of many genes. In spleen, we found basal p53 to regulate known target gene *Polh,* a translesion polymerase capable of UV-induced DNA damage repair that is known to be deficient in xeroderma pigmentosa (Supplementary Figure 5A)[31–33]. In basal thymus, a poorly characterized gene *Lyrm7* was identified as a single and novel target (Supplementary Figure 5B). Taken together, our data show that basal p53 in the liver also regulates genes with known tumor inhibiting function, such as the well-described pro-apoptotic gene *Bbc3* and genes that maintain chromosomal stability in the basal liver and spleen.

### Comparison of basal and irradiated p53

We next wished to compare p53 binding patterns in tissue under basal and irradiated conditions. p53 ChIP-seq in basal murine liver, thymus, and spleen was performed in parallel with litter-matched control mice that were irradiated. As anticipated, irradiation resulted in significantly greater p53 occupancy genome-wide as detected by ChIP-seq (Figure 3A-B). The pattern of tissue-specific p53 peaks between liver, spleen, and thymus were similar between irradiated and basal settings (Figure 2A, 3C).

**Figure 3.**
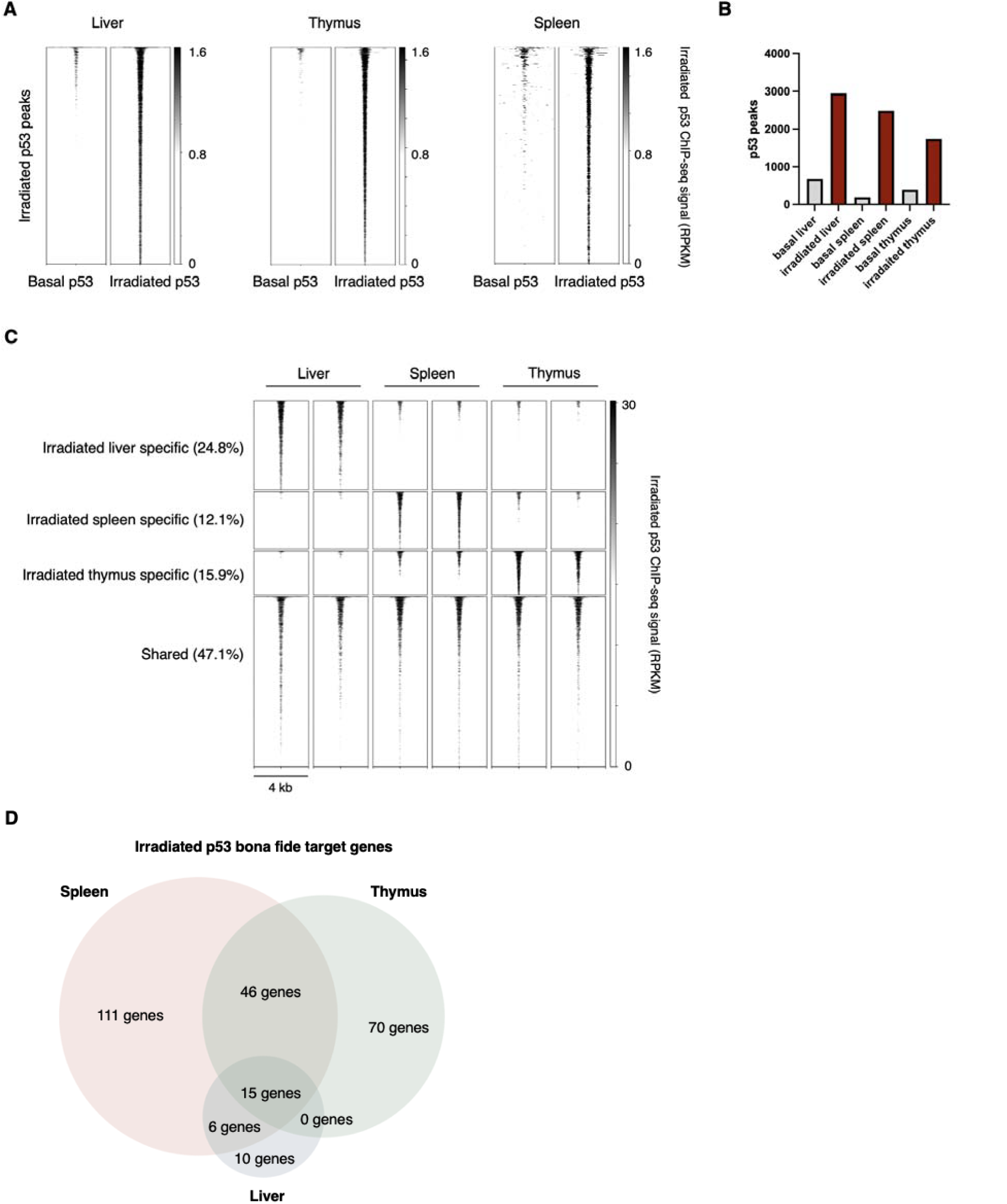
A comparison of irradiated p53 occupancy and *bona fide* target genes between tissue types. (A) Heatmap of p53 ChIP-seq signal (RPKM) in basal or irradiated murine liver, spleen, or thymus. (B) Histogram of number of p53 peaks. (C) Heatmap of irradiated p53 ChIP-seq signal (RPKM) in murine liver, spleen, or thymus. Tissue specific genes are those where p53 signal is > 1 log_2_ fold change greater than other tissue types. Each tissue type shown in biological replicates. (D) Venn diagram of irradiated p53 target genes shared between various tissues.

Regarding peaks associated with mRNA expression changes, we observed more of a consensus among the different irradiated tissues compared to basal (Figure 3D). Whereas none of the 42 basal p53 targets we identified were shared among different tissues, ∼28% (73 out of 258 target genes) were shared among irradiated tissues (Figure 2B, 3D). This suggests basal p53 may regulate more tissue-specific transcriptional programs, as compared to a more common transcriptional program activated by multiple tissues in response to irradiation.

In contrast to basal conditions, irradiated p53 target genes were nearly all previously described, with few exceptions. This data suggests the study of basal p53 reveals transcriptional targets previously missed by studies focusing on DNA damage response. 15 genes were observed to be activated in all three tissues upon irradiation and 67 genes were shared in two or more irradiated tissues, many of which were previously described as p53 target genes including *Cdkn1a, Bbc3, Bax, Cd80,* and *Mdm2* (Supplementary Table 1, 2). Pathway and gene ontology analyses showed an enrichment for DNA damage response, known p53 target genes, and p53 regulation (Supplementary Figure S6A-B, 7A-B).

Two of the 15 target genes shared in all irradiated tissues were, to our knowledge at the time of writing, not previously described as p53 target genes. They are *Dglucy,* a poorly characterized D-glutamate cyclase, and *Zfp958,* the murine ortholog of the human zinc finger protein *ZNF117* (Supplementary Figure 6C-J)[34]. Similarly, only three of the 67 target genes shared in two or more tissues were not previously described—the stearoyl-CoA desaturase *Scd2,* which converts saturated fatty acids into mono-unsaturated fatty acids during lipid metabolism, the mitochondrial protein glutaryl-CoA dehydrogenase *Gcdh,* and the uncharacterized zinc finger protein *Zfp688* (Supplementary Figure 7D-H)[35].

Among the individual tissues, we identify four out of the ten liver-specific irradiated p53 target genes to be previously undescribed—the glutaminyl cyclase *Qpct,* the lymphatic vessel endothelial hyaluronan receptor *Lyve1,* the neurodevelopmental gene *Nrxn1,* and the glutamine transporter *Slc1a4* (Supplementary Table 3; Supplementary Figure S8). We similarly identify seven out of the 99 spleen-specific irradiated p53 target genes to be previously undescribed, including the poorly characterized *Pnma1,* the chromatin modulator *Brpf3,* the carbonic anhydrase *Car2,* the voltage gated chloride channel *Clcn3,* the kinesin protein *Kifc5b,* the chemokine receptor *Ccrl2,* and the transcriptional activator *Mef2b* (Supplementary Table 4; Supplementary Figure S9). Lastly, seven of the 70 irradiated thymus-specific p53 target genes were not previously described, including *F8a, Tmem131l* a negative regulator of thymocyte proliferation, the methyl-lysine reader and transcriptional repressor *L3mbtl3* observed to be deleted in medulloblastoma, the antiviral gene *Slfn2* which we previously observed as a basal colon p53 target, *Ndrg3,* the NF-κB activator *Unc5cl,* and *Axdnd1* (Supplementary Table 5; Supplementary Figure S10)[36–38].

### Most basal p53 target genes are not activated upon irradiation

We next wished to assess how irradiation affects the expression of our identified basal p53 target genes, in our three tissues with matched basal-irradiation pairs, processed in parallel. Unsupervised hierarchical clustering analysis revealed two types of basal targets: one group that was up-regulated between p53 WT and null in both basal and irradiated conditions, and a second group that was up-regulated only under basal conditions (Figure 4A-B). Surprisingly, most of the basal p53 target genes fell into the latter category, including *Bbc3,* which was upregulated under basal conditions, but not upon irradiation in liver despite increased p53 binding (Figure 4A-B, Supplementary Figure 11). This agrees with previous observations that liver is not a radiosensitive tissue[39]. We only observed three p53 target genes, all previously known, that were up-regulated under both basal and irradiated settings in their respective tissues: *Ddias* which encodes a DNA damage induced apoptosis suppressor, *Ckap2* which encodes a microtubule stabilizing protein that promotes cell cycle arrest and apoptosis, and *Trp53* itself[40, 41].

**Figure 4.**
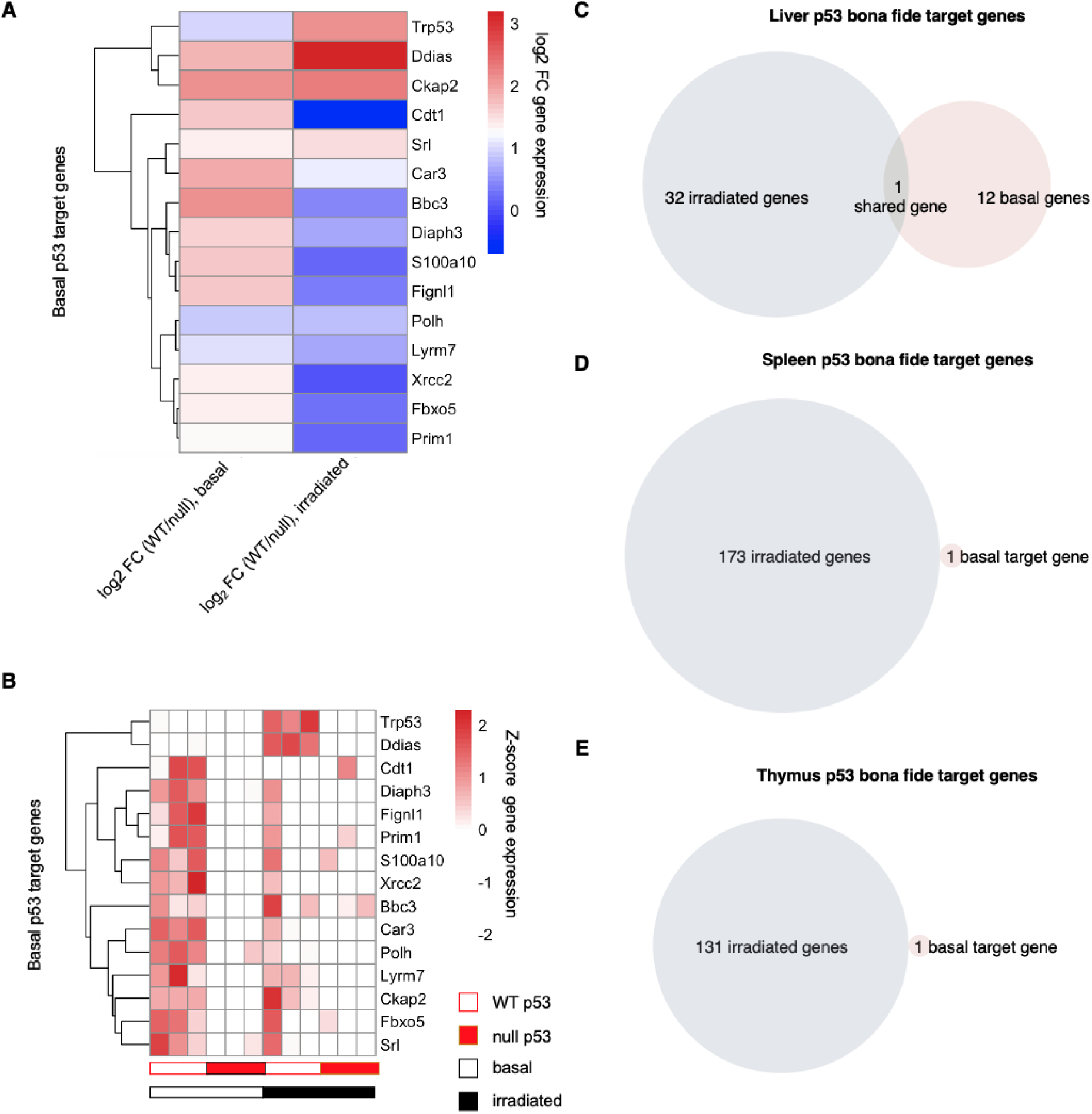
Irradiation largely does not activate basal p53 target genes *in vivo*. (A) Heatmap of basal p53 target genes (spleen, liver, and thymus combined) log_2_ fold change gene expression between WT and null under irradiated and basal settings. (B) Heatmap of basal p53 target genes raw expression, normalized by row. (C-E) Venn diagram showing overlap of basal and irradiated p53 target genes in spleen, thymus, and liver.

Indeed, when comparing p53 targets, we observed virtually no overlap in p53 target genes in tissues between basal and irradiated states. Between the 33 target genes in irradiated liver and 13 genes in basal liver, only one gene was shared, while none were shared between irradiated and basal thymus and spleen (Figure 4C-E). Taken together, these data suggest the basal p53 program to be unique compared to its DNA-damage program, and highlights the significance of using basal stress in physiologic tissue to uncover p53 function.

### Transcription factor motifs within p53 peaks

An important area of discovery remains how other transcription factors modulate p53 binding and activation of target genes. We first separated called p53 peaks into two bins, depending on whether a p53 consensus sequence was identified within the peak. In the basal setting, we observed p53 signal to be largely comparable regardless of whether a p53 consensus site was present (Supplementary Figure 12). However, upon irradiation, p53 peaks with a consensus binding sequence clearly exhibited stronger signal than those without (Supplementary Figure 12).

We also observed a difference in p53 occupancy locations between these two groups. p53 peaks with a consensus motif site were predominantly located at intergenic sites and introns, whereas peaks without a consensus motif site were predominantly in promoters (Supplementary Figure 13).

In the basal setting, most tissues such as liver, spleen, and thymus, only a minority of p53 peaks (26-32%) were found to contain a consensus binding sequence (Figure 5A). The exception to this was basal colon, where 76% of p53 peaks were observed to contain a consensus binding sequence. Unsupervised hierarchical clustering showed distinct transcription factor motifs present within p53 peaks with versus without a p53 consensus binding sequence in the basal setting (Figure 5B). In basal thymus, spleen, and liver, p53 peaks without a consensus sequence were notably enriched for *Hif1a, Myc,* and *Egr1* motifs (Figure 5B). Indeed, p53 is known to bind HIF-1α and “piggyback” onto HIF-1α at HIF-1α response elements, which validate our motif analyses[42–44]. Similarly, EGR1 and p53 protein-protein interactions have been described[45]. Conversely, p53 peaks with a consensus sequence were enriched for *Rora, Tead1,* and *Nfic* motifs (Figure 5B). The exception was basal colon, with similar transcription factor motif distribution regardless of whether a p53 consensus binding sequence was present (Figure 5B). This may reflect batch effect, as basal colon ChIP-seq was processed separately from other tissues versus a true tissue-specific difference.

**Figure 5.**
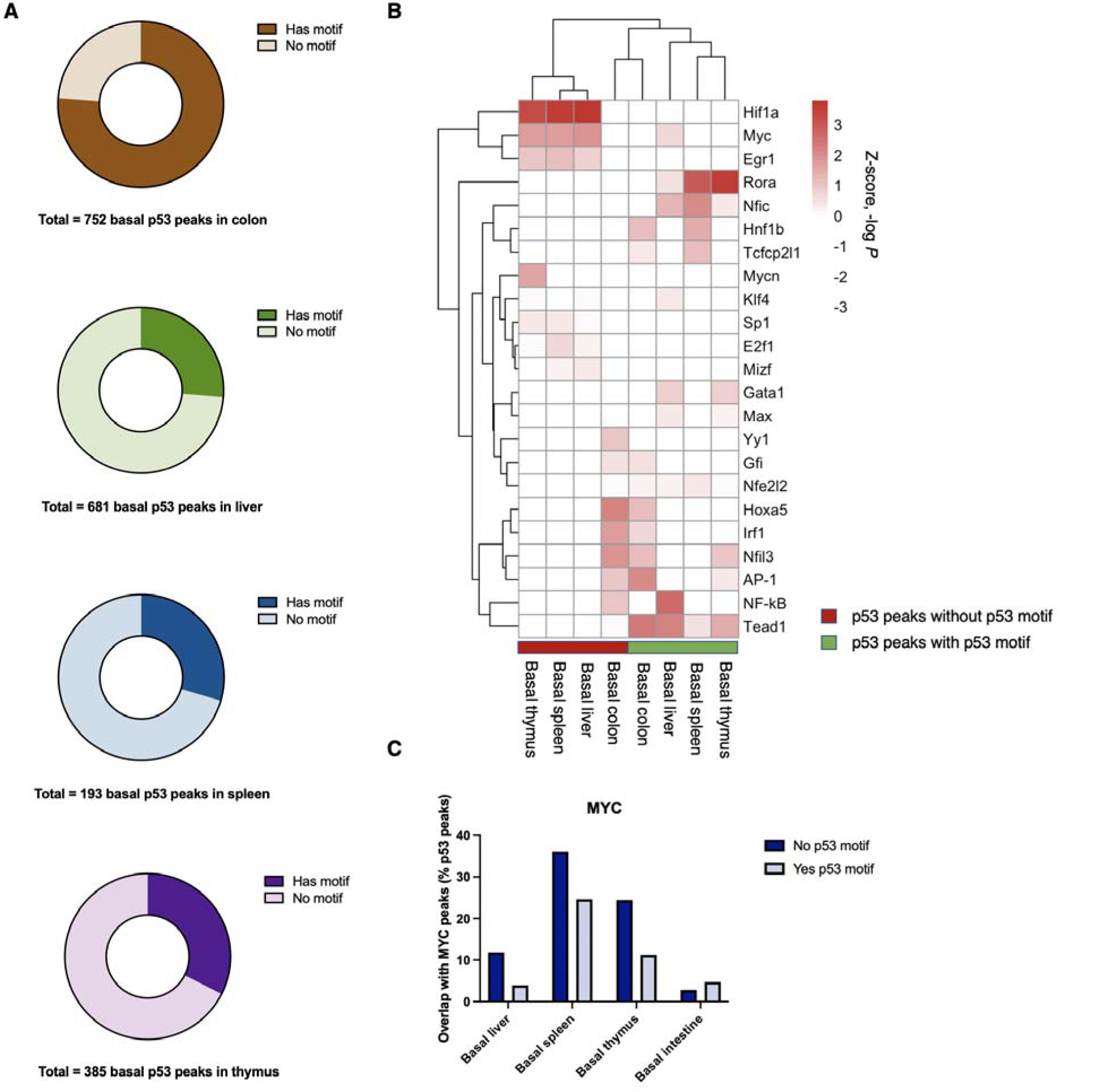
Transcription factor motif analysis at p53 peaks. (A) Pie chart of basal p53 peaks based on whether a p53 consensus binding sequence is present. (B) Heatmap and unsupervised hierarchical analysis of transcription factor motif analysis (by significance of motif enrichment) within p53 peaks depending on whether a consensus binding sequence is present. Only expressed transcription factors are included—a non-expressed transcription factor (TPM < 1) is assigned a *P* = 1. (C) Overlap of p53 and MYC peaks in various basal tissues depending on whether a p53 consensus binding sequence is present.

Upon irradiation, the proportion of p53 peaks containing a canonical binding motif rose in all tissues, to as high as 82% and 86% in irradiated spleen and thymus, respectively (Supplementary Figure 14A). The same trend is observed in the irradiated tissue setting as well: *Hif1a* and *Myc* motifs are enriched in peaks without a canonical consensus site whereas *Tead1* and *Gata1* are enriched in peaks with a canonical consensus site (Supplementary Figure 14B). Similarly, EGR1 and p53 as well as GATA-1 and p53 protein-protein interactions have been described[45, 46].

To validate our motif analyses, we queried appropriate, tissue-matched public ChIP-seq datasets. We did not find tissue-matched public ChIP-seq datasets for HIF-1α, NFIC, TEAD1, RORα, and GATA1, but found suitably complete datasets for MYC. We observed a greater overlap of MYC and p53 peaks if no underlying p53 consensus sequence was present, corroborating our motif analyses (Figure 5C). Furthermore, this trend was not observed in basal colon, where we observed a poor enrichment of MYC motifs among p53 peaks (Figure 5C). An alternative explanation is the prevalence of MYC and its E-box sites at promoters, as we observe p53 preferentially binding to promoters if no consensus sequence is present (Supplementary Figure 13)[47]. Taken together, these data provide insight to guide future studies dissecting whether these candidate factors indeed influence or guide basal p53 occupancy. Taken together, our analyses highlight alternative methods and hypotheses for non-canonical p53 DNA binding.

## Conclusion

Our aim is to further assess whether basal p53 maintains tumor suppressor expression *in vivo* and, given the well-described context-dependent function of p53, to further expand the known list of p53 target genes. In this study, we observed basal p53 regulation of several established tumor suppressors, including *Phlda3, Bbc3, Xaf1,* and itself. We did not observe significant overlap between tumor suppressors regulated by basal p53 in murine tissues versus MCF10A cells in culture from our previous study, which may reflect human-mouse differences or tissue-specific differences. We find basal p53 maintaining the expression of several anti-viral genes, including *Mx1, Mx2,* as well as *Slfn2* a gene required for T cell mediated immunity in basal colon. We also find p53 regulation of DNA integrity genes such as *Xrcc2, Fignl1,*and *Cdt1* in basal liver, and *Polh* in basal spleen. Furthermore, the vast majority of basal p53 target genes were not induced upon irradiation, suggesting basal p53 operates a transcriptional program that is largely distinct from its DNA damage response. Similarly, the basal p53 target gene repertoire is unique to each tissue type.

In most basal tissues, except for colon, the majority of p53 peaks were not found to have an underlying consensus sequence. We find that basal, but not irradiated, p53 binds with largely equal signal strength regardless of whether an underlying consensus sequence was present. We show that, interestingly, basal p53 without a consensus sequence favors binding at promoters whereas those with a consensus sequence favored introns and intergenic sites. Furthermore, co-enrichment of other transcription factor motifs at basal p53 peaks differed depending on whether a consensus sequence was present. Taken together, we expand upon our previous study and characterize the basal p53 landscape, transcriptional programs, and binding preferences in the setting of physiologic mouse tissue via fresh tissue ChIP-seq.

## Materials and Methods

### Mice

All animal work was conducted in accordance with an Icahn School of Medicine at Mount Sinai Institutional Care and Use Committee (IACUC) protocol. For the initial colon p53 ChIP-seq and RNA-seq experiments, 8-week-old C57/Bl6 mice were used. For the subsequent liver, spleen, and thymus p53 ChIP-seq and RNA-seq experiments, 3-week-old C57/Bl6 mice were used. Both male and female mice were used. For irradiation, mice were exsposed to 2Gy irradiation in a Precision X-ray, X-RAD 32. Pie chambers were used for immobilization with no shielding. X-rayed and control mice were held in holding cages for 3 hours at which point they were sacrificed by regulated and approved exposure to carbon dioxide for 4 minutes compliant with euthanasia guidelines.

### Mouse colon epithelium isolation

The following section of mouse colon was dissected: distal to the cecum and proximal to the rectum. A plastic catheter was threaded through one opening of the colon and stool was flushed out using ice cold PBS. The colon was then dissected longitudinally with scissors to expose the luminal mucosa. This was rinsed three times with ice cold PBS. To specifically isolate the mucosal epithelium, a glass microscope slide was used to gently scrape the mucosal surface and the top layer of cells collected. The rest of the colon, including the fibrous mesenchyme, was discarded.

### ChIP-seq and library construction

ChIP and library construction was performed as previously described[11]. In brief, bulk tissues were pressed through a 70 μM cell strainer (BD Falcon #352350) to generate a single cell suspension and crosslinked in 1% formaldehyde in PBS for 10 minutes at room temperature. Given difficulty in sonicating colon epithelium, we treated them with 0.25% trypsin (Corning) in rotation for three minutes at room temperature before quenching with 10% FBS in RPMI media prior to passing through the cell strainer. Whereas mouse spleen, thymus, and liver were single-crosslinked with 10% formaldehyde, mouse colon epithelium was double-crosslinked with 0.25 M disuccinimidyl glutarate for 45 minutes at room temperature in addition to 1% formaldehyde. Cross-linked cells were sonicated on a Diagenode Bioruptor for 15 cycles on the low setting for 30s ON 30s OFF. Rest of protocol as previously described. For ChIP, 10 μg the mouse p53 polyclonal antibody CM5 was used from Leica (#P53-PROTEIN-CM5) or control mouse IgG. For pulldown, Protein A Dynabeads were used (Millipore Sigma #16-661). For mouse colon epithelium, library construction as previously described and sequenced by the Mount Sinai Oncological Sciences Sequencing Core Facility on an Illumina NextSeq 550 instrument as 75 bp single end reads[11]. For mouse liver, thymus, and spleen, library construction and sequencing were performed by the Epigenomics Core Facility at Weill Cornell Medicine.

### ChIP-seq data analysis

The cutadapt script was used to trim barcode contamination from Illumina sequencing reads. Computational analyses were performed on the using the MINERVA supercomputing system at the Icahn School of Medicine at Mount Sinai. Single-end ChIP-seq reads were mapped using bowtie (version 1.3.0) to the mm9 mouse reference genome under the following parameters[48]. Paired-end ChIP-seq reads (from those sequenced at Weill Cornell) were mapped using bowtie2 (version 2.4.1) under the default parameters[48]. Bigwig tracks were generated using deeptools bamCoverage with RPKM nornmalization. Heatmaps were generated using deepTools computeMatrix and plotProfile[49, 50]. For the heatmap in Figure 3B, RPKM signal was normalized across samples to best allow for comparison. This was done using deepTools bamCoverage with scaleFactor, normalizing to the mean RPKM signal at all called peaks for each sample. Please see Supplementary Table __ for quality control metrics for ChIP-seq datasets generated in this study. Motif analysis was performed using TRAP for single or multiple DNA sequences where appropriate[51–53]. Parameters used included JASPAR vertebrate motifs as the matrix file, mouse promoters as the background file, and Benjamini-Hochberg for multiple test correction.

### RNA isolation and library construction

Extracted organ pieces, or harvested colon epithelium, were immediately placed in RNAlater during dissection and stored at 4 degrees Celsius until RNA isolation using the RNeasy Mini Kit (QIAGEN) following the manufacturer’s instructions. Library construction and sequencing were performed by the Epigenomics Core Facility at Weill Cornell Medicine. For mouse spleen, thymus, and liver, sequencing libraries were prepared with the TruSeq Stranded Total RNA kit from Illumina, from 1 μg of total RNA. For mouse colon epithelium, the Mount Sinai Oncological Sciences Sequencing Core Facility prepared poly-A-capture RNA-seq libraries using the TruSeq RNA Library Prep Kit v2 (Illumina) according to manufacturer’s instructions. All libraries were sequenced as 75 bp single end reads by the Mount Sinai Oncological Sciences Sequencing Core Facility on an Illumina NextSeq 550 instrument.

### RNA-seq data analysis

Sequenced reads were mapped to the mm10 transcriptome using Salmon quasi-mapping under the default parameters and expression output in TPM was extracted[54]. Output files (expression in TPM) were directly integrated into the R package DESeq2 for differential gene expression analysis, again under default parameters[55]. Genes with expression >2 TPM were used for downstream analyses. Pathway analyses were performed using pre-ranked Gene Set Enrichment Analysis (GSEA), using the pre-ranked (by log_2_ fold change gene expression) function, as well as Enrichr (which includes many tools including ChEA)[56, 57]. Log_2_ fold change cut-offs for differentially expressed genes used in downstream analyses are stated in the text.

### Statistical analysis

All experiments were performed with at least three biological replicates. Significance was determined by unpaired two-tailed Student’s *t*-test with significance defined as *P* < 0.05. All error bars represent standard error around the mean.

## Data Availability

Irradiated mouse p53 ChIP-seq and RNA-seq datasets performed by our group in parallel with this study were previously published and accessed under the accession number GSE203643 and GSE204924[39]. Mouse liver MYC ChIP-seq data was accessed under the accession number GSE83869[58]. Mouse intestine MYC ChIP-seq data was accessed under the accession number GSE56008[59]. Mouse B cell from spleen MYC ChIP-seq data was accessed under the accession number GSE51004[60]. Mouse T cell MYC ChIP-seq data was accessed under the accession number GSE58075[61].

---

**Supplementary Figure 1.**
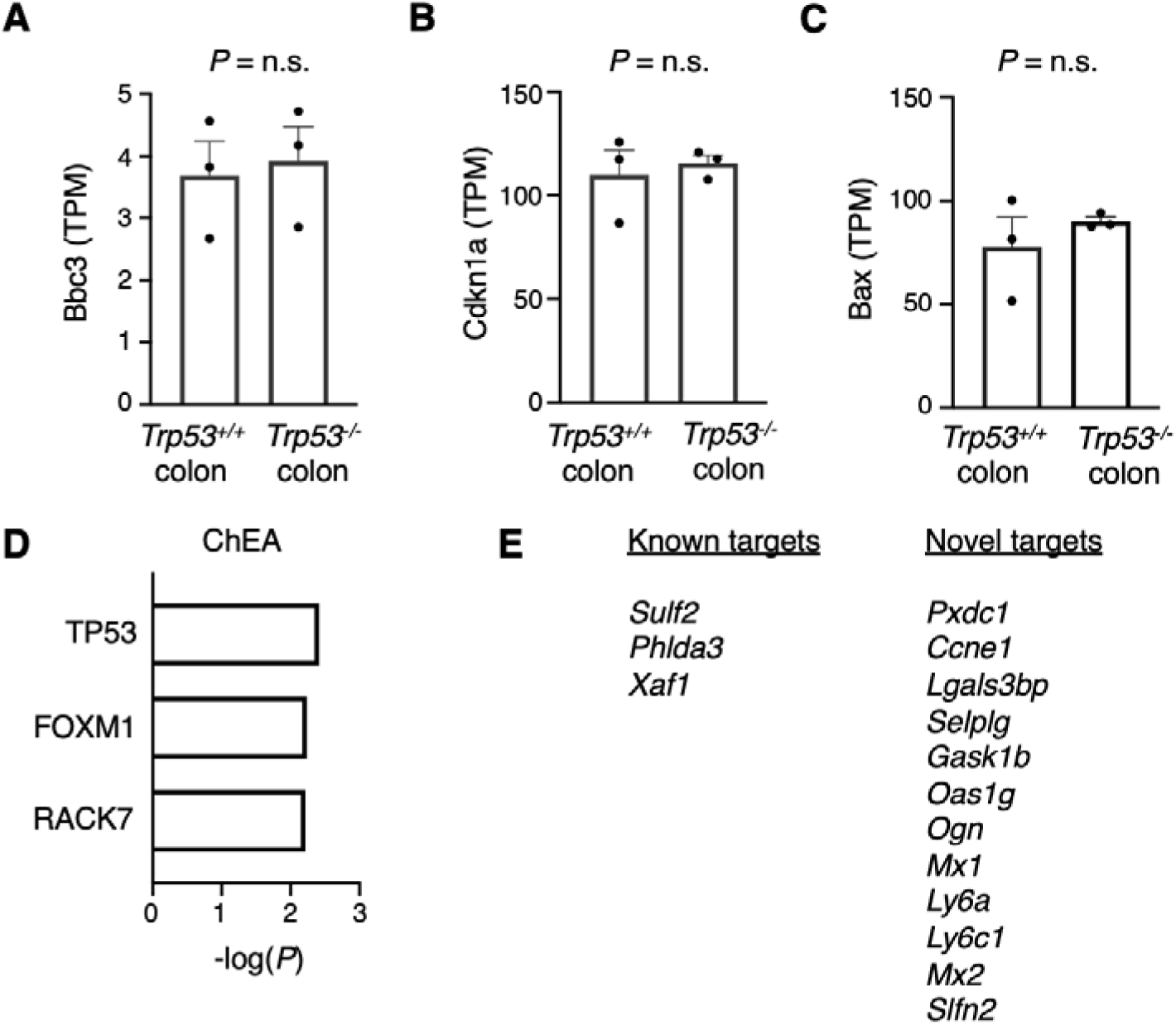
Basal p53 occupancy in physiologic colon identifies known and novel targets. (A-C) Gene expression of known p53 target genes in physiologic murine colon epithelium via RNA-seq between *n* = 3 WT and *n* = 3 null mice. (D) ChEA transcription factor analysis of identified p53 targets. (E) Bona fide basal p53 target genes in physiologic murine colon epithelium, defined by p53 occupancy by ChIP-seq and differential expression between WT and null colon by >1 log_2_ fold change.

**Supplementary Figure 2.**
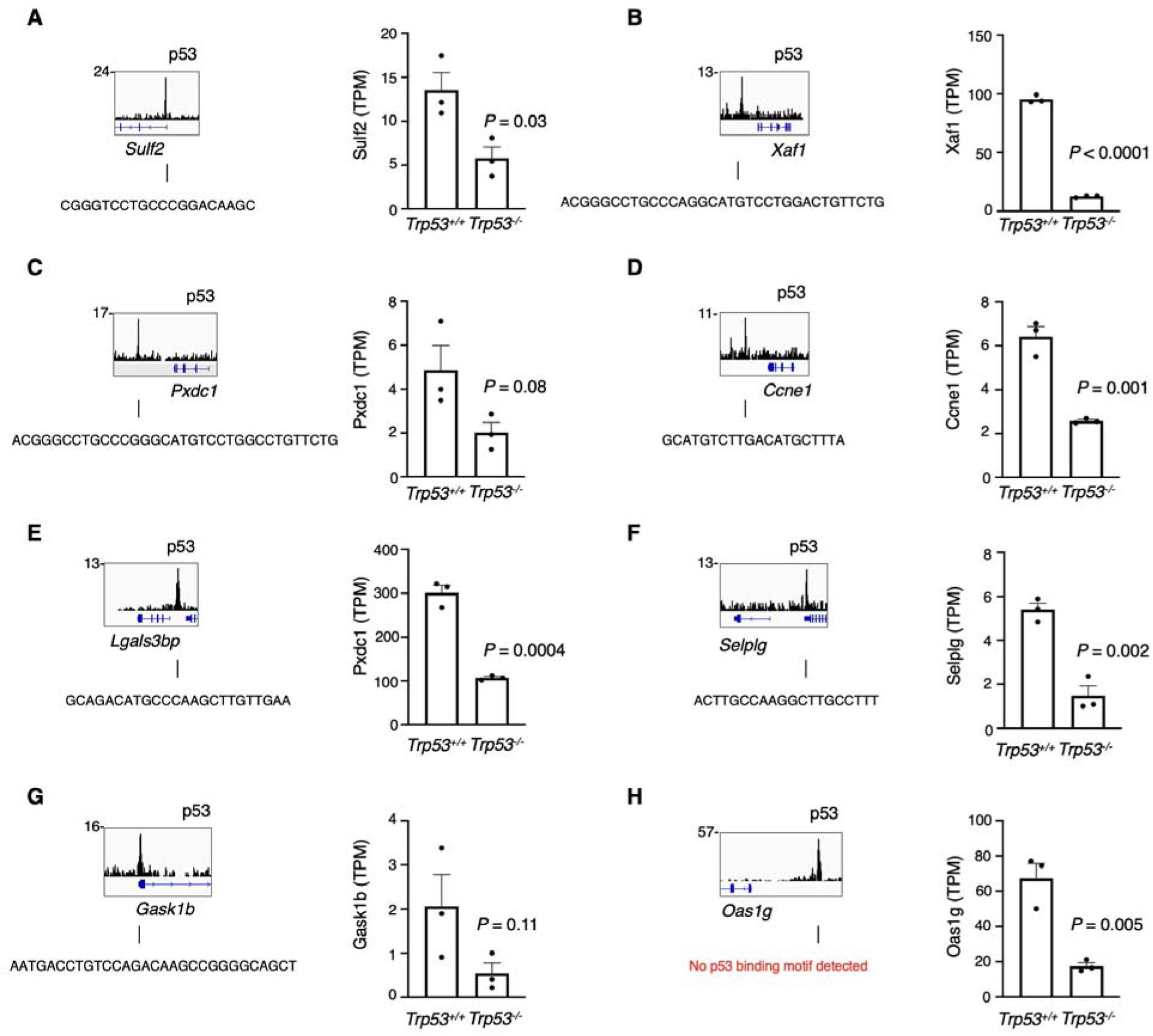
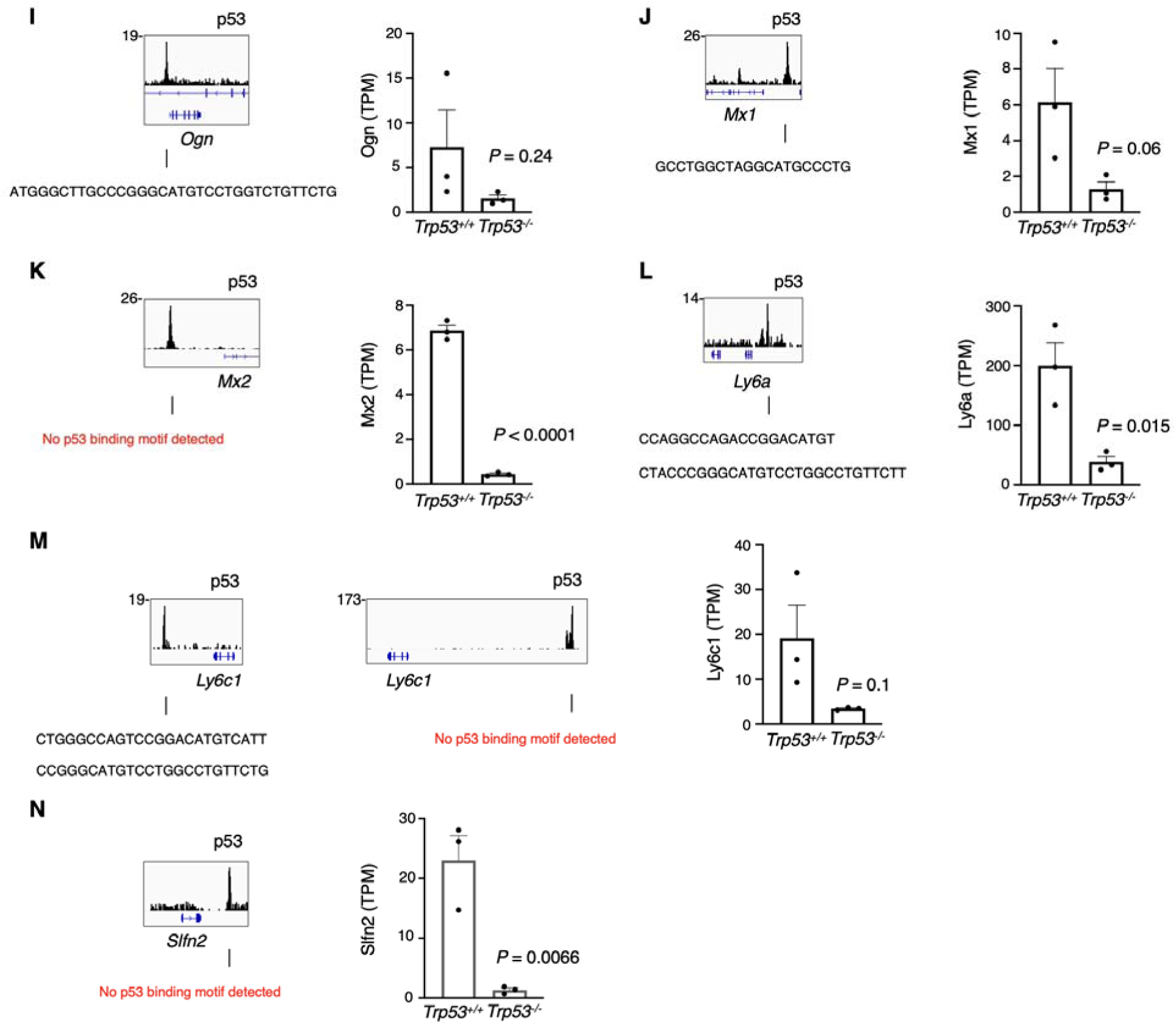
Basal p53 target genes in physiologic murine colon epithelium. p53 ChIP-seq tracks at basal p53 target genes from Figure 1. DNA sequences under called peaks were subject to TRAP motif analysis for *de novo* discovery of p53 binding motifs. Expression of target genes by RNA-seq between p53 WT (*n* = 3) or null (*n* = 3) murine colon epithelium.

**Supplementary Figure 3.**
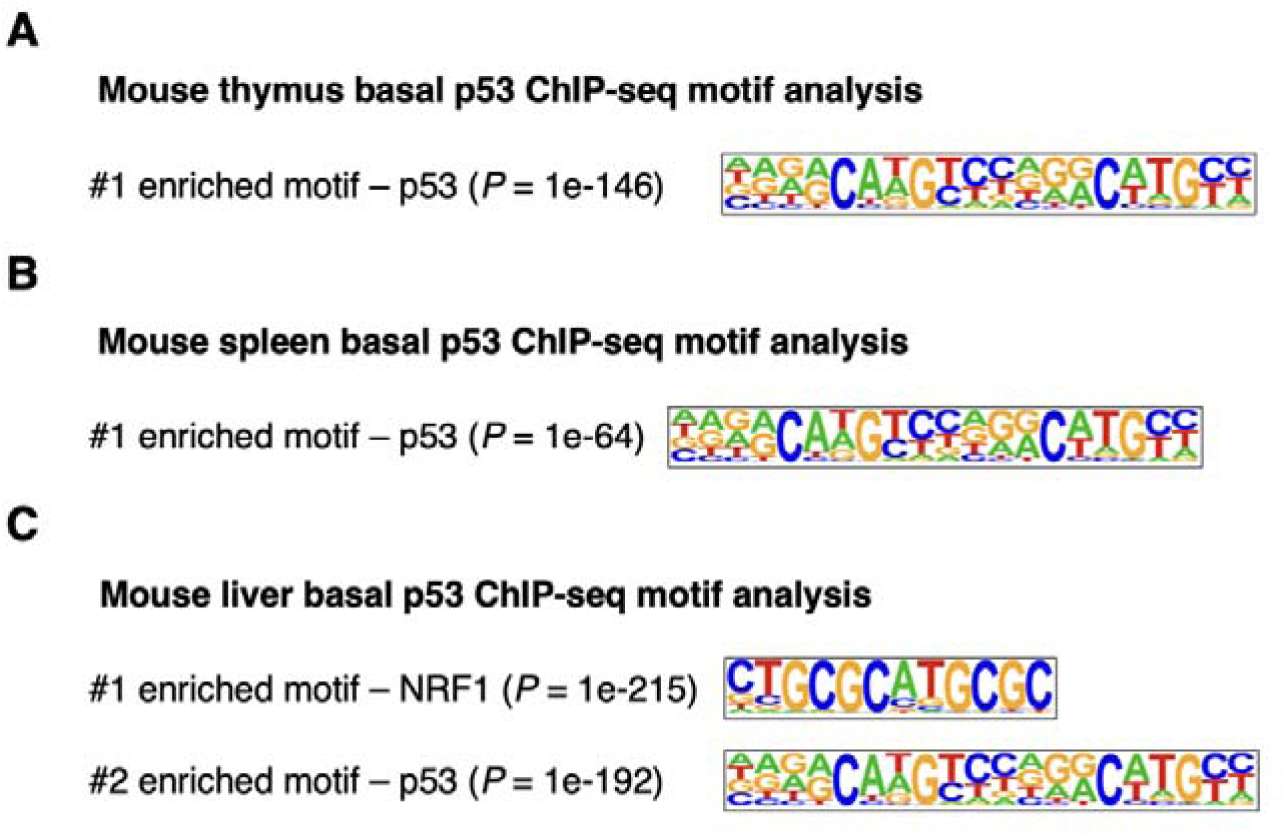
Basal p53 peaks are enriched for its consensus binding sequence. HOMER *de novo* transcription factor motif discovery of called basal p53 peaks in murine (A) thymus, (B) spleen, or (C) liver.

**Supplementary Figure 4.**
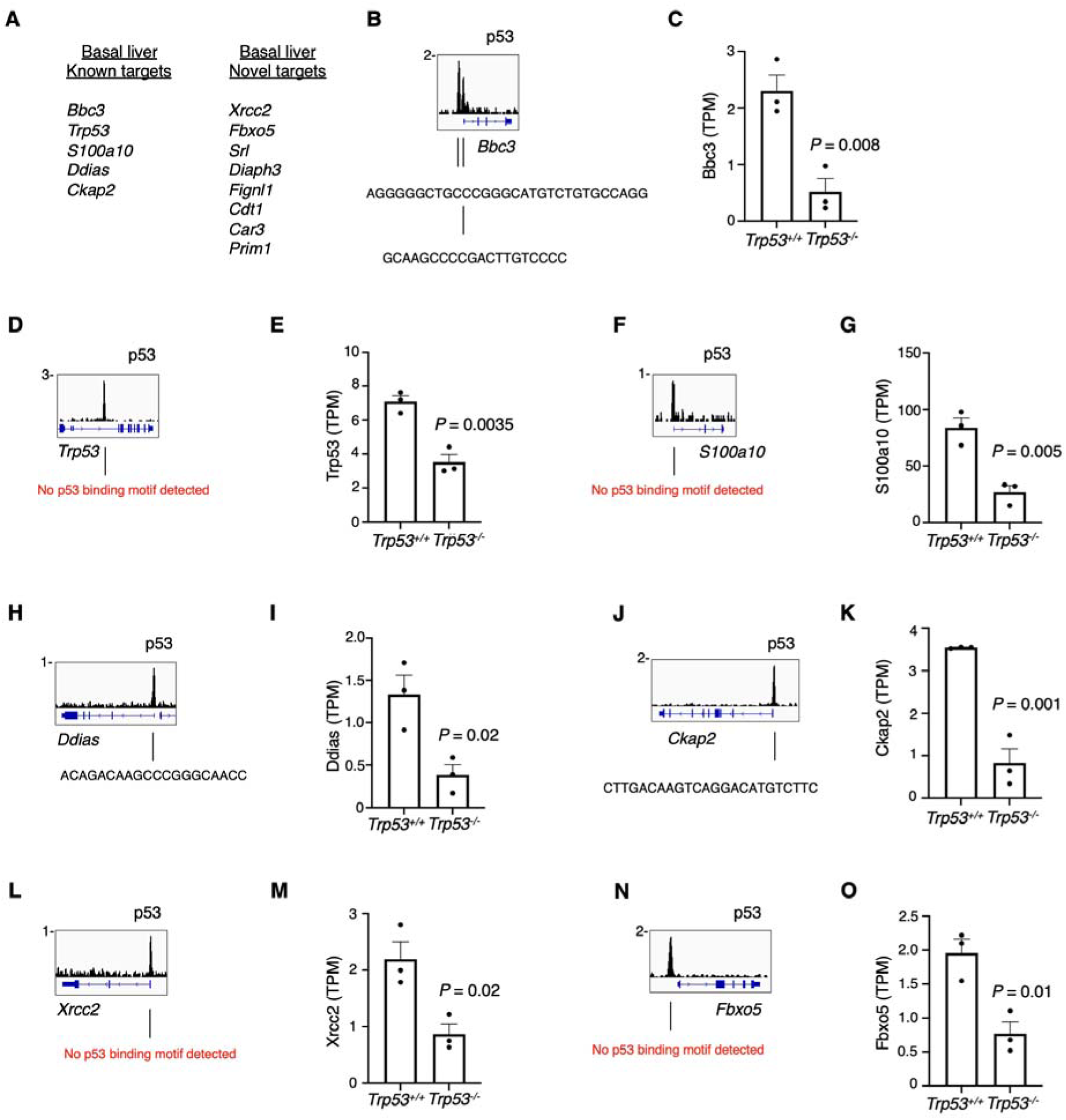
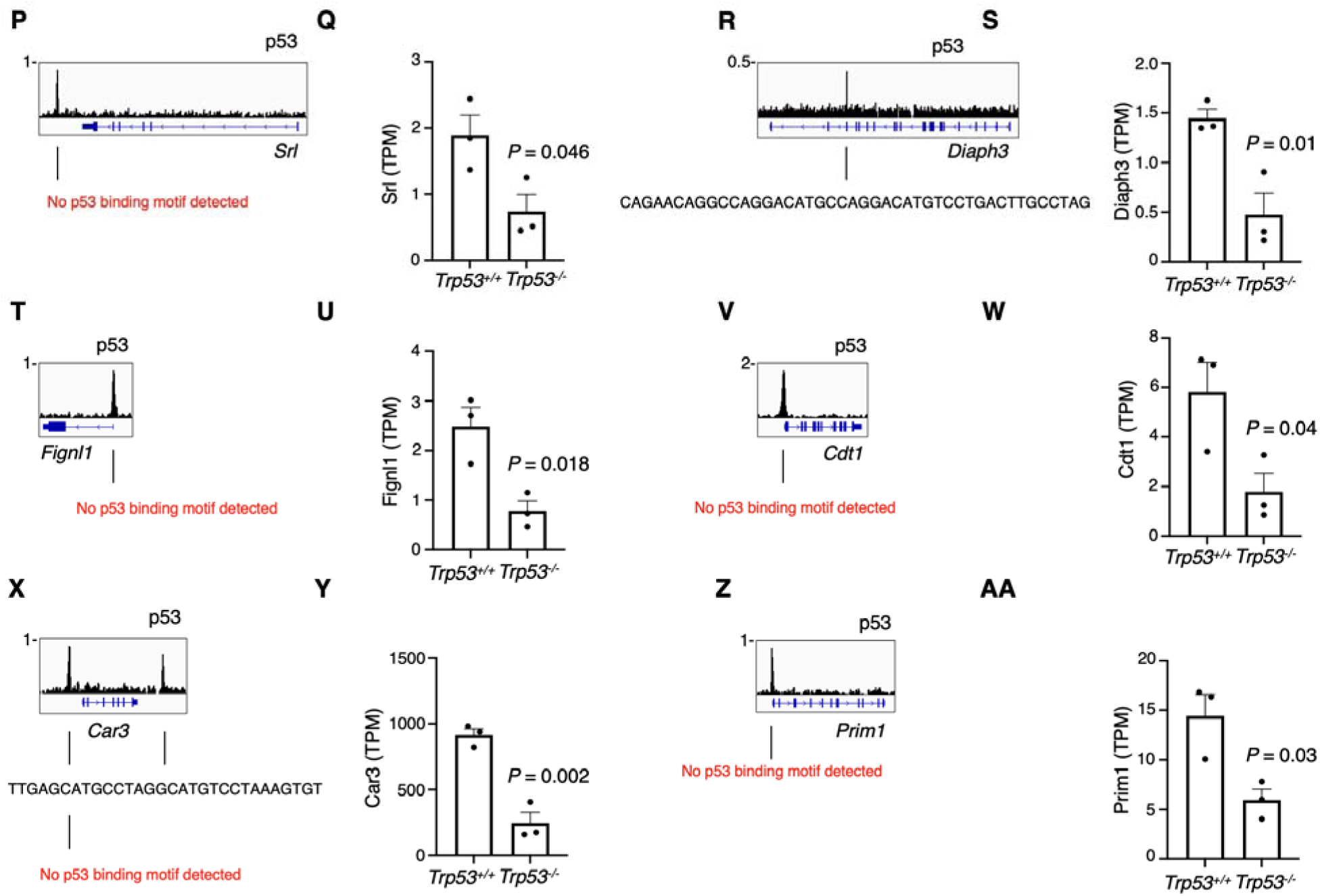
Basal p53 target genes in physiologic murine liver. p53 ChIP-seq tracks at basal p53 target genes identified in murine liver listed in panel (A). DNA sequences under called peaks were subject to TRAP motif analysis for *de novo* discovery of p53 binding motifs. Expression of target genes by RNA-seq between p53 WT (*n* = 3) or null (*n* = 3) murine liver.

**Supplementary Figure 5.**
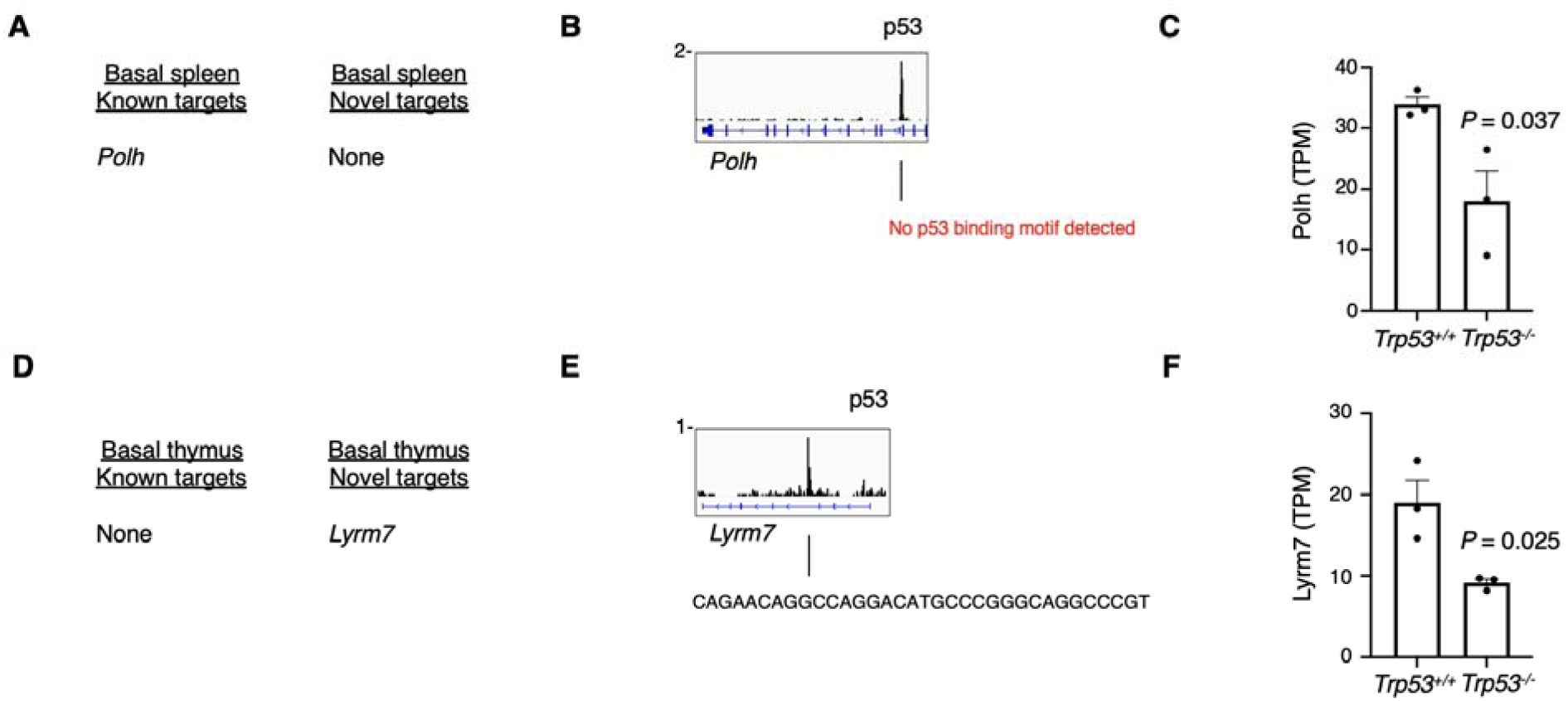
Basal p53 target genes in physiologic murine spleen and thymus. (A) p53 ChIP-seq tracks at basal p53 target genes identified in murine spleen. (B) DNA sequences under called peaks were subject to TRAP motif analysis for *de novo* discovery of p53 binding motifs. (C) Expression of target genes by RNA-seq between p53 WT (*n* = 3) or null (*n* = 3) murine spleen. (D) p53 ChIP-seq tracks at basal p53 target genes identified in murine thymus. (E) DNA sequences under called peaks were subject to TRAP motif analysis for *de novo* discovery of p53 binding motifs. (F) Expression of target genes by RNA-seq between p53 WT (*n* = 3) or null (*n* = 3) murine thymus.

**Supplementary Figure 6.**
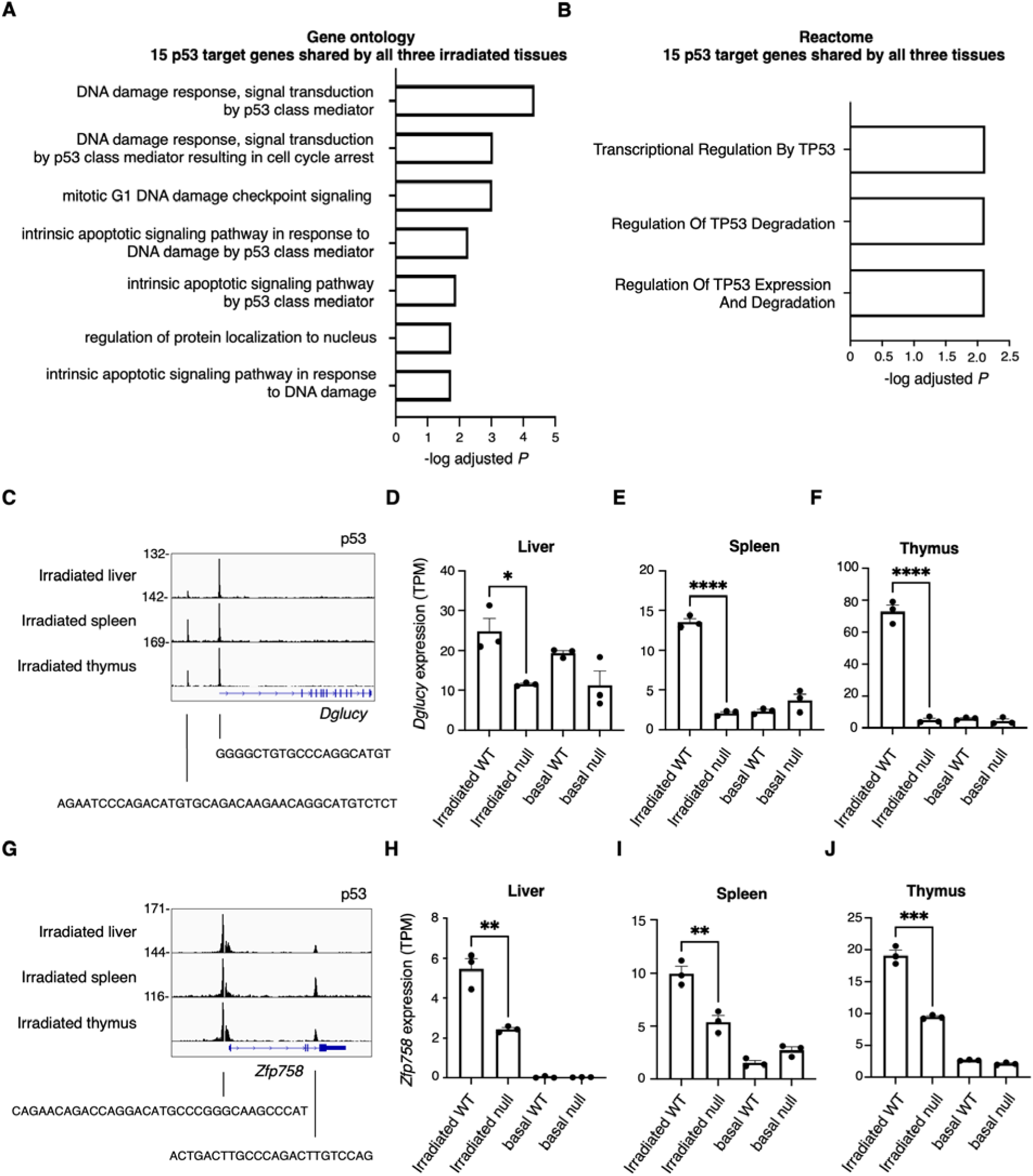
Irradiated p53 target genes activated in all three tissues. (A) Gene ontology and (B) Reactome pathway analysis of 15 p53 target genes activated in irradiated spleen, thymus, and liver. (C) Irradiated p53 ChIP-seq at the *Dglucy* locus. (D-F) *Dglucy* expression by RNA-seq in liver, spleen, and thymus. (G) Irradiated p53 ChIP-seq at the *Zfp958* locus. (H-J) *Zfp958* expression by RNA-seq in liver, spleen, and thymus.

**Supplementary Figure 7.**
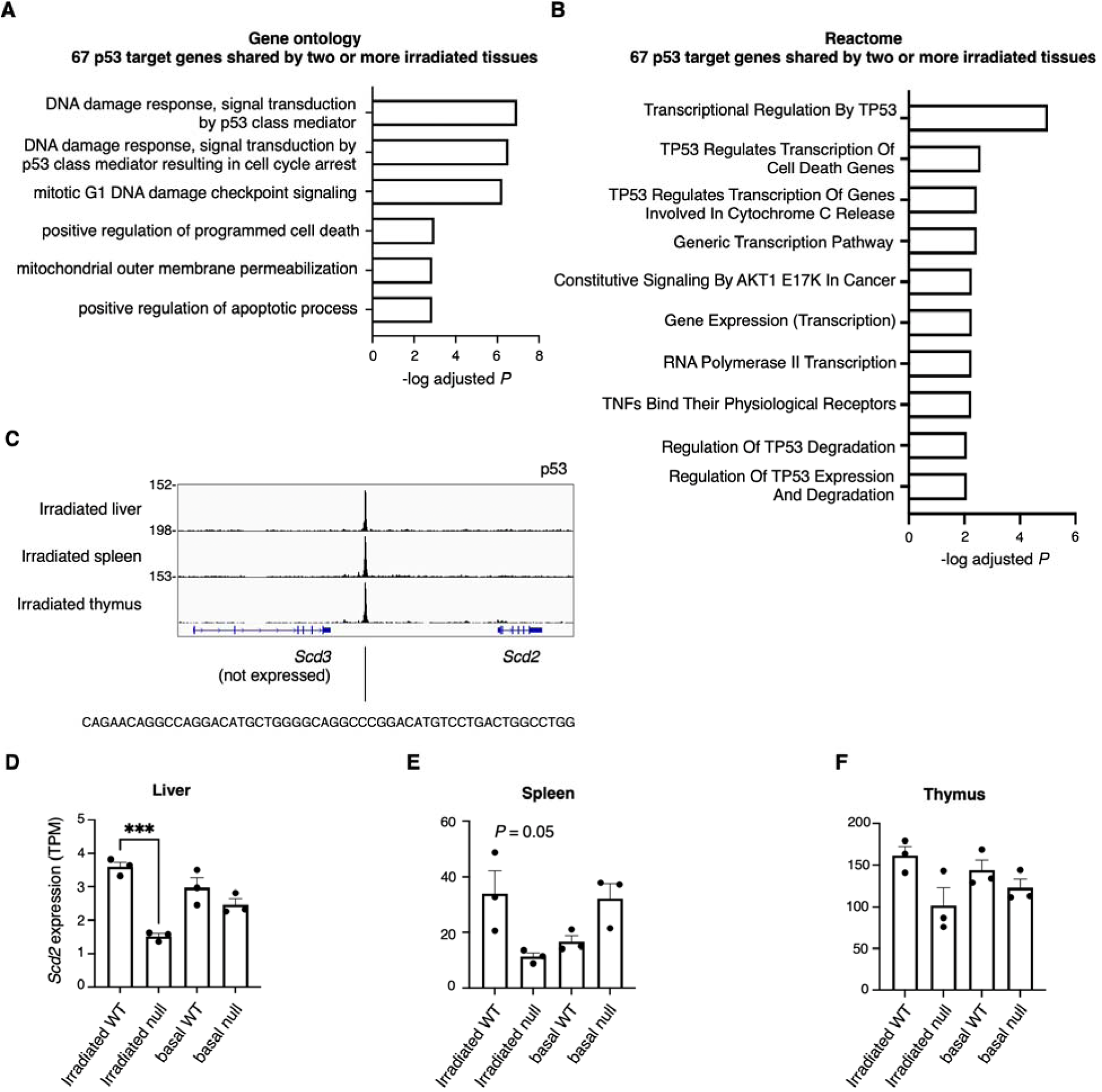
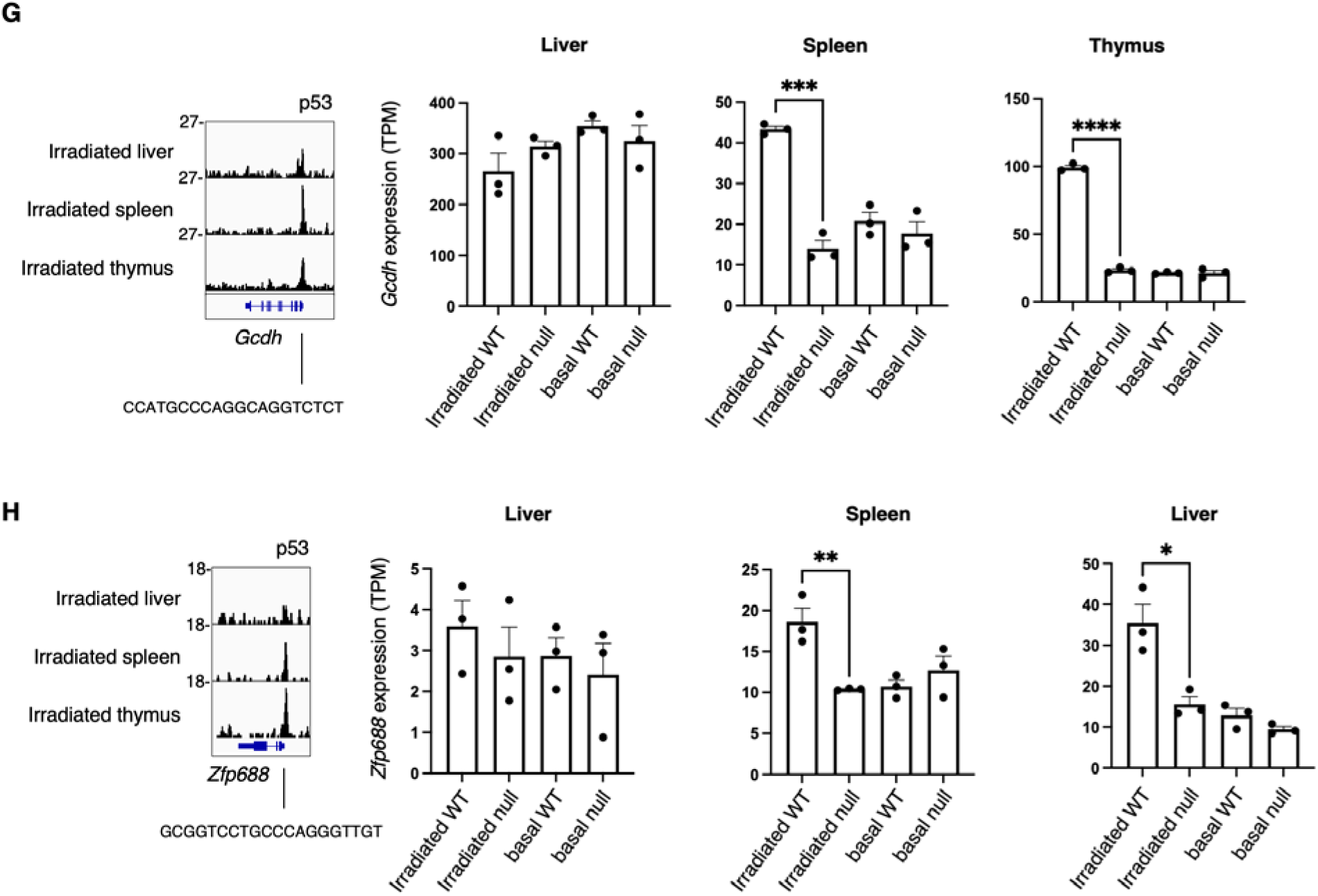
Irradiated p53 target genes activated in two or more tissues. (A) Gene ontology and (B) Reactome pathway analysis of 67 p53 target genes activated in two or more irradiated tissues. (C) Irradiated p53 ChIP-seq track at the *Scd2* locus. (D-F) *Scd2* expression by RNA-seq in liver, spleen, and thymus. (G) Irradiated p53 ChIP-seq track at th *Gcdh* locus, with RNA-seq gene expression. (H) Irradiated p53 ChIP-seq track at the *Zfp688* locus with RNA-seq gene expression.

**Supplementary Figure 8.**
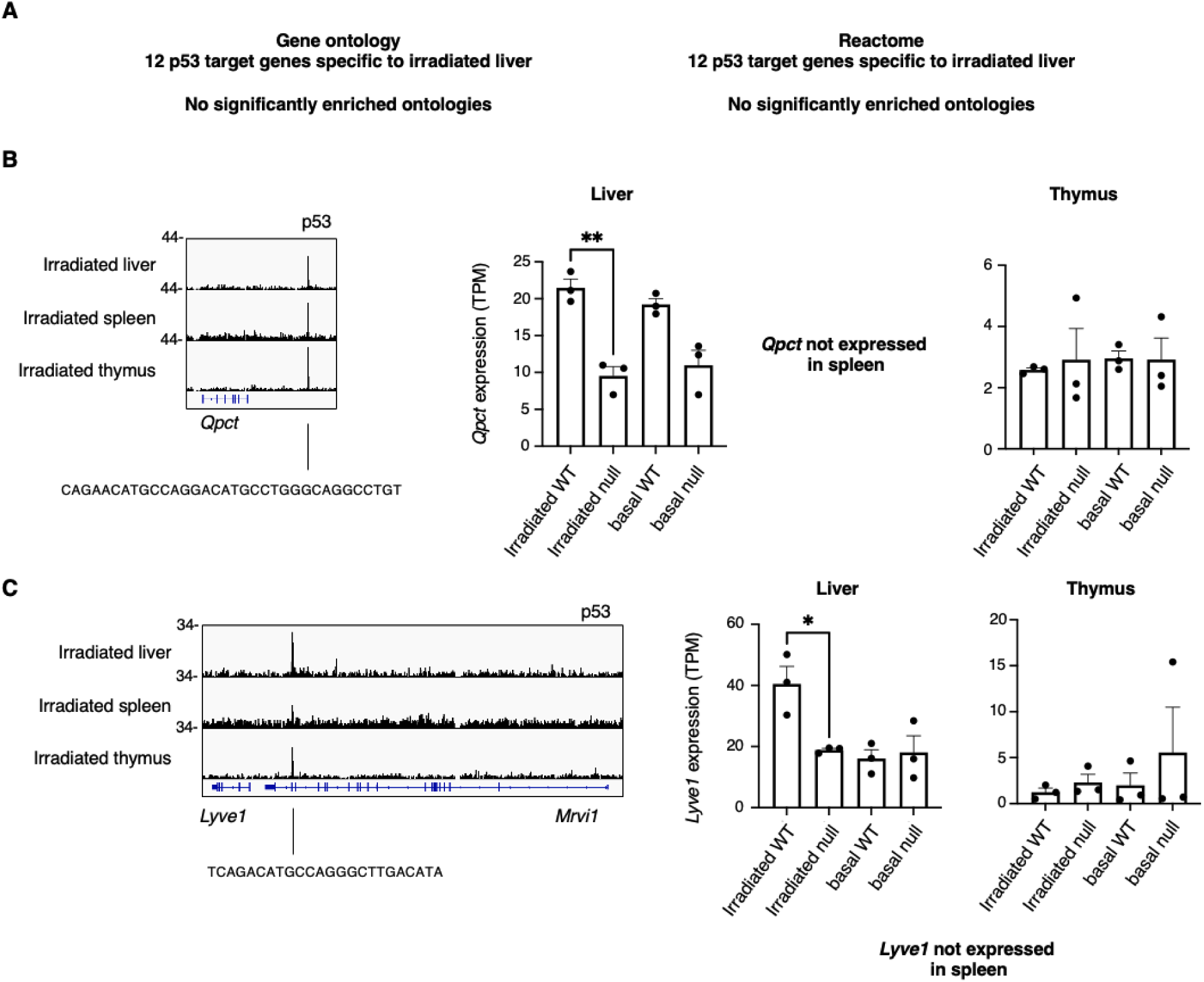
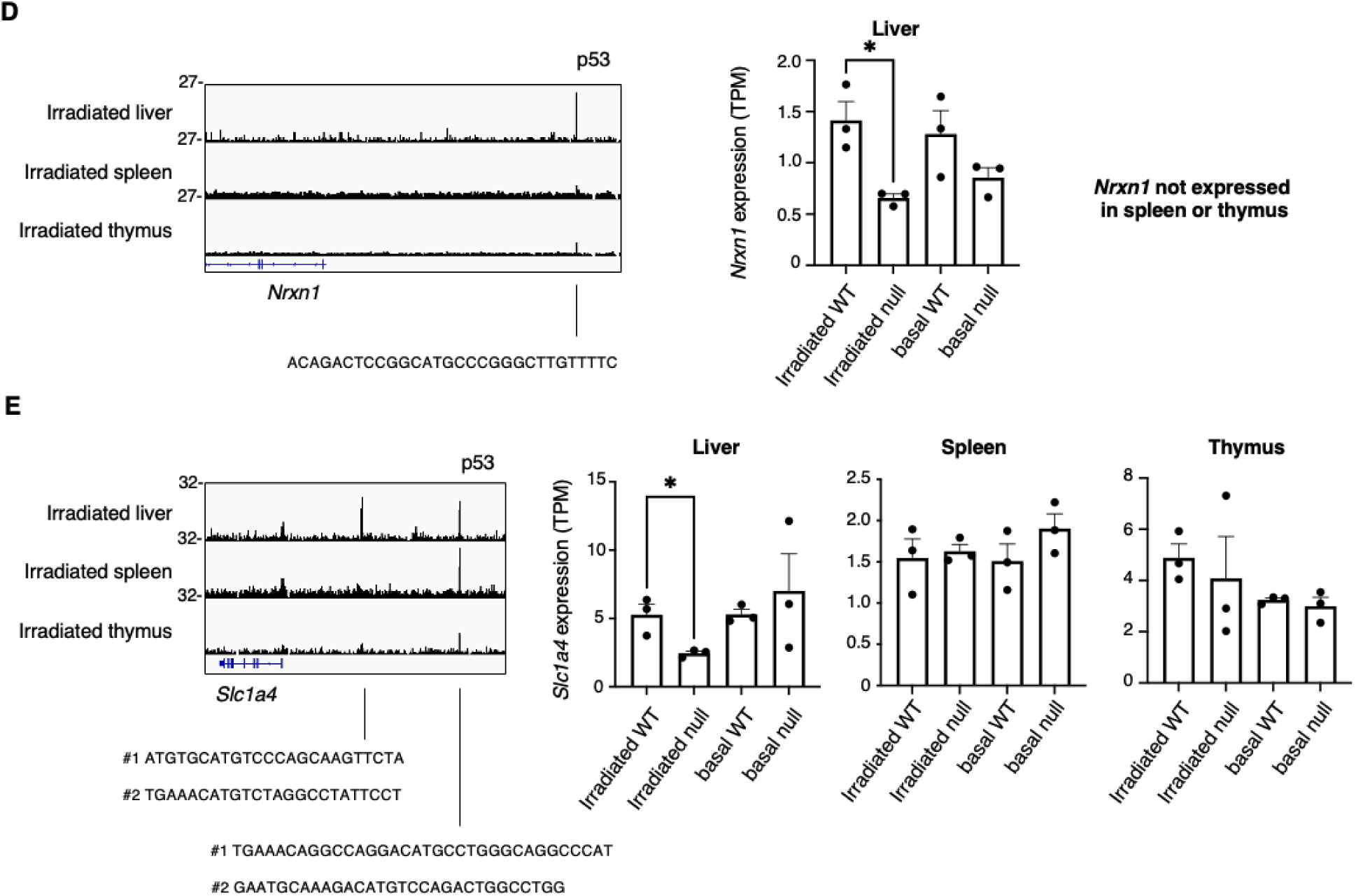
Irradiated p53 target genes specific to liver. (A) Gene ontology and Reactome pathway analysis of 12 p53 target genes activated inly in irradiated liver showed no significant enrichment. (B) Irradiated p53 ChIP-seq at the *Qpct* locu and *Qpct* expression by RNA-seq in liver, spleen, and thymus. (C) Irradiated p53 ChIP-seq at the *Lyve1* locus and *Lyve1* expression by RNA-seq in liver, spleen, and thymus. (D) Irradiated p53 ChIP-seq at the *Nrxn1* locus and *Nrxn1* expression by RNA-seq in liver, spleen, and thymus. (E) Irradiated p53 ChIP-seq at the *Slc1a4* locus and *Slc1a4* expression by RNA-seq in liver, spleen, and thymus.

**Supplementary Figure 9.**
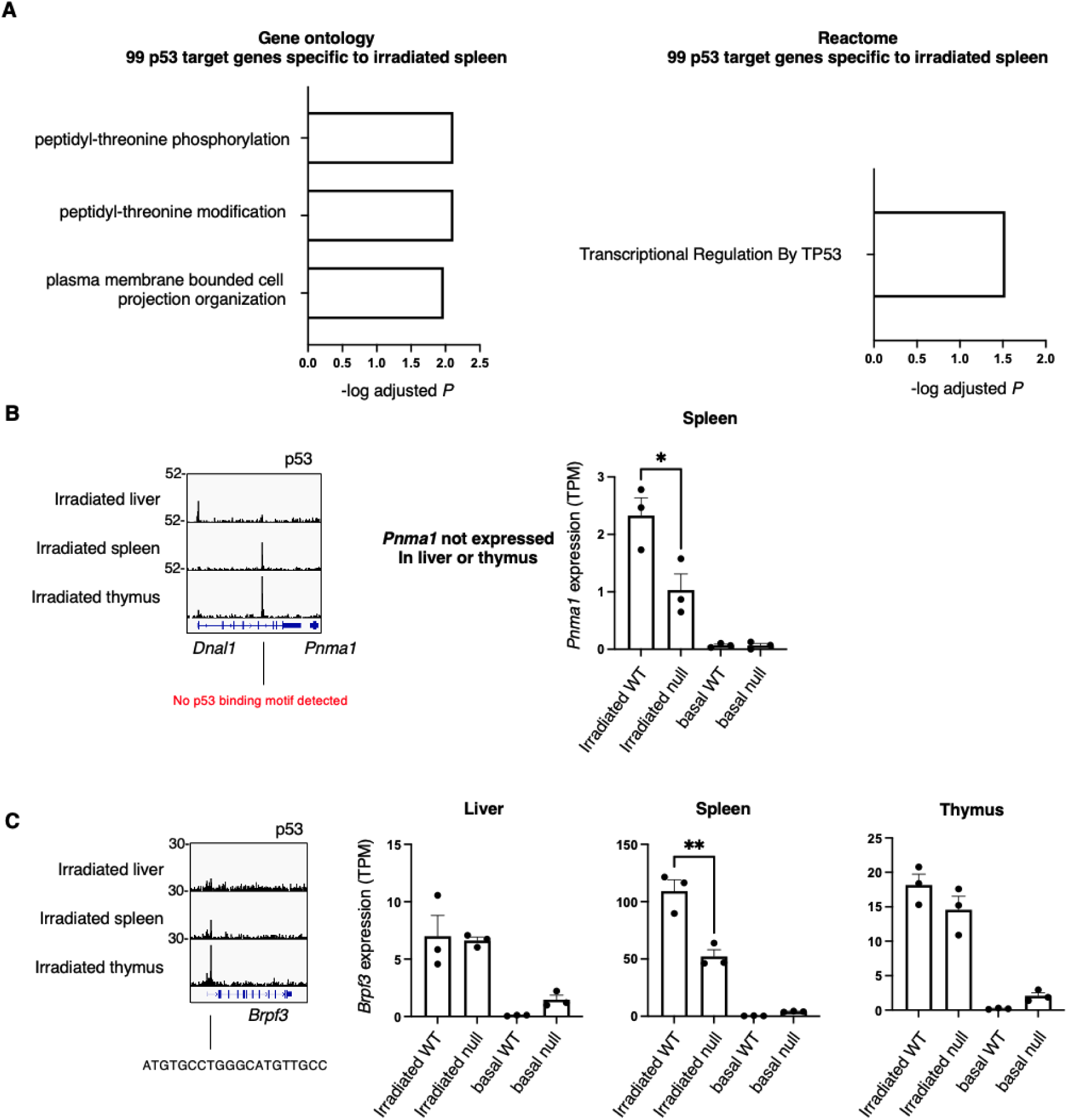
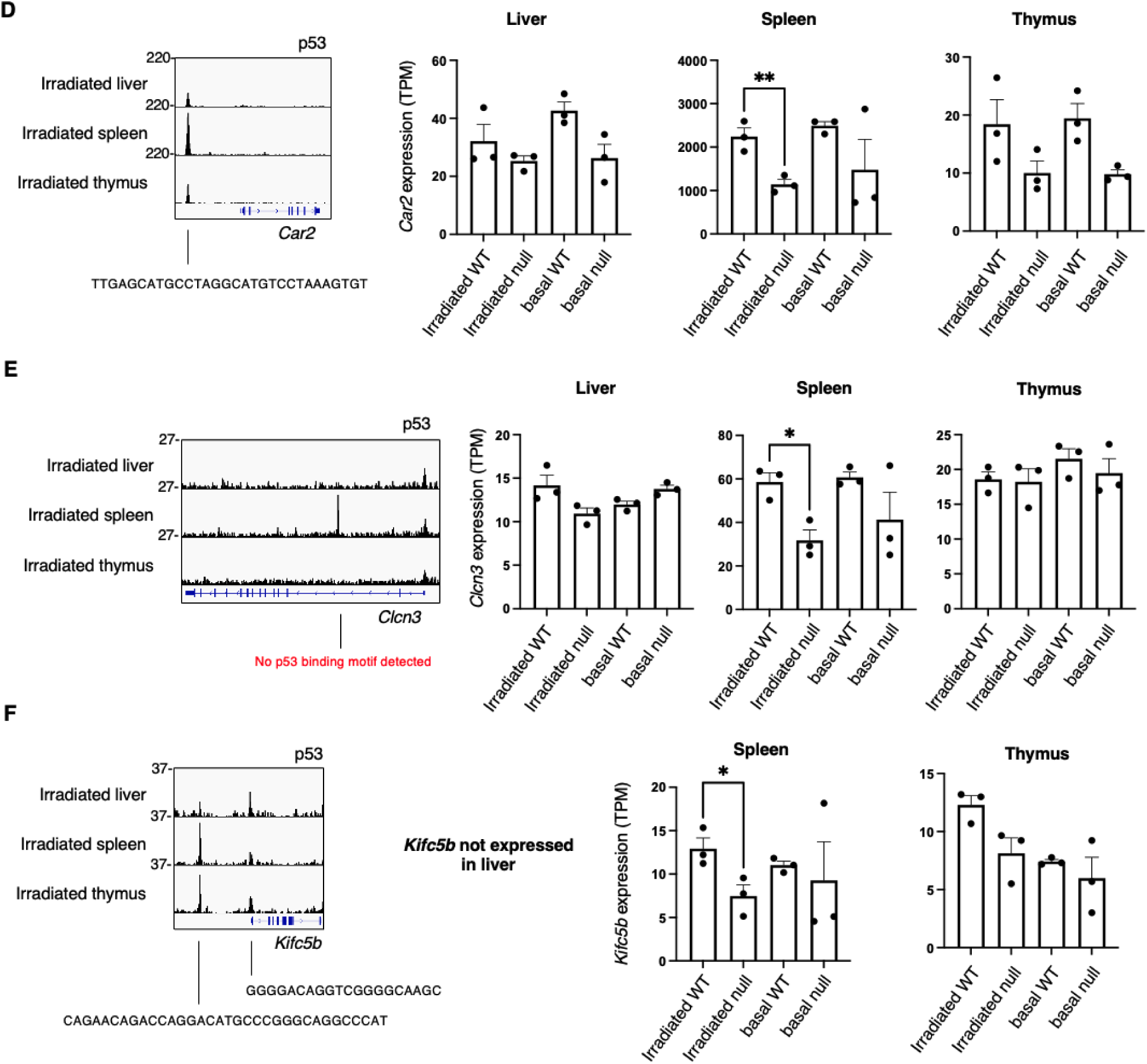
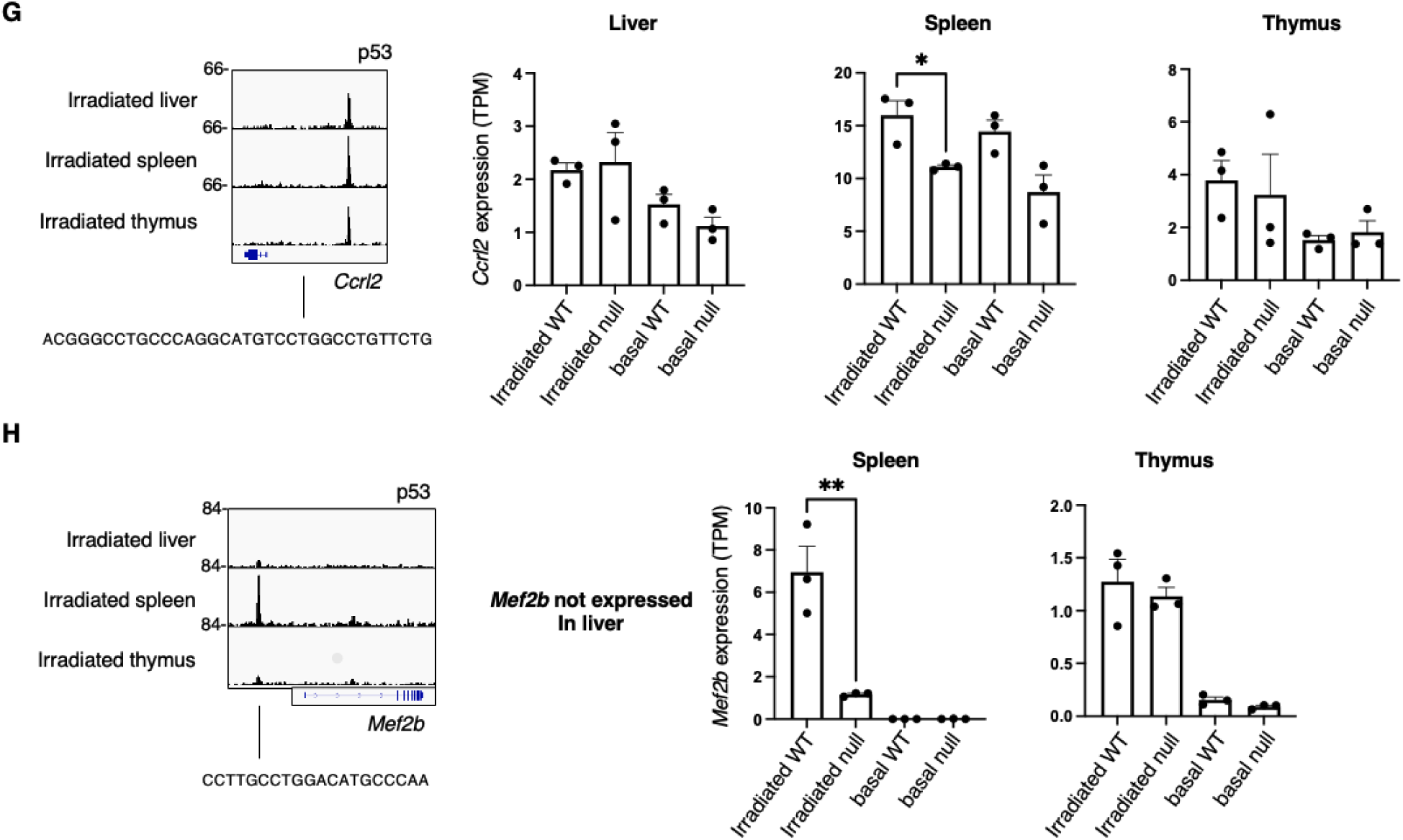
Irradiated p53 target genes specific to spleen. (A) Gene ontology and Reactome pathway analysis of 99 p53 target genes activated only in irradiated spleen. (B) Irradiated p53 ChIP-seq at the *Pnma1* locus and *Pnma1* expression by RNA-seq in liver, spleen, and thymus. (C) Irradiated p53 ChIP-seq at the *Brpf3* locus and *Brpf3* expression by RNA-seq in liver, spleen, and thymus. (D) Irradiated p53 ChIP-seq at the *Car2* locus and *Car2* expression by RNA-seq in liver, spleen, and thymus. (E) Irradiated p53 ChIP-seq at the *Clcn3* locus and *Clcn3* expression by RNA-seq in liver, spleen, and thymus. (F) Irradiated p53 ChIP-seq at the *Kifc5b* locus and *Kifc5b* expression by RNA-seq in liver, spleen, and thymus. (G) Irradiated p53 ChIP-seq at the *Ccrl2* locus and *Ccrl2* expression by RNA-seq in liver, spleen, and thymus. (H) Irradiated p53 ChIP-seq at the *Mef2b* locus and *Mef2b* expression by RNA-seq in liver, spleen, and thymus.

**Supplementary Figure 10.**
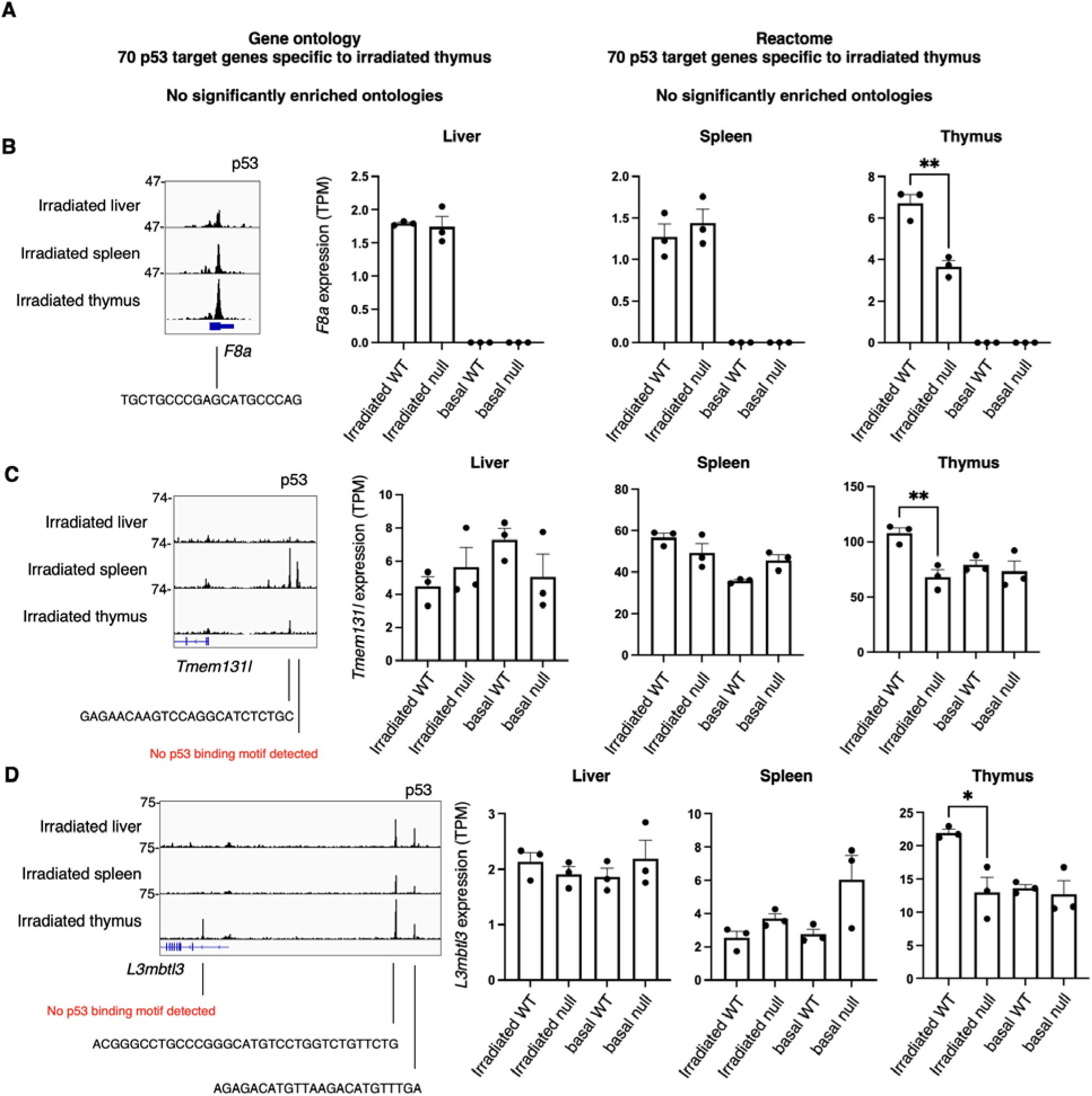
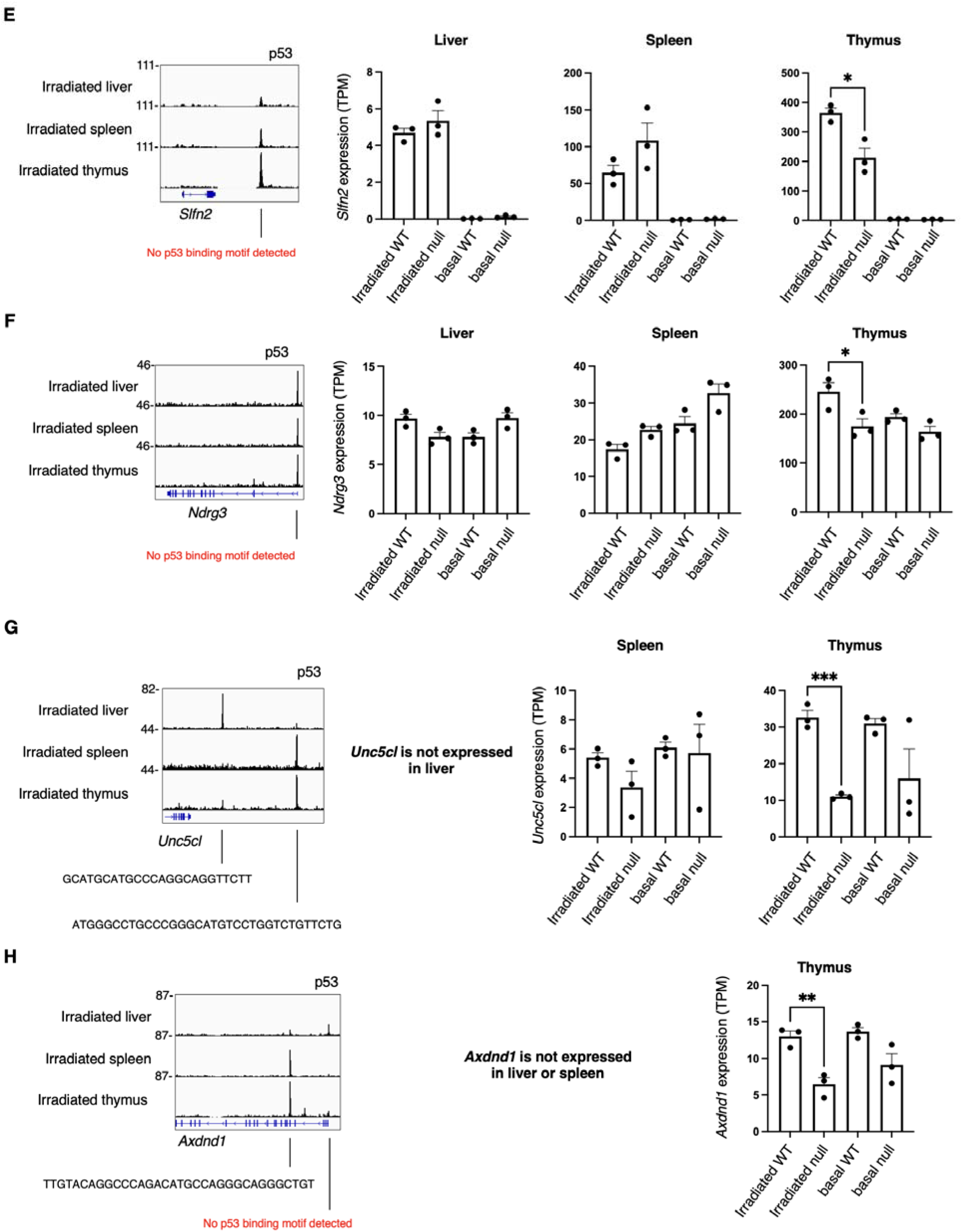
Irradiated p53 target genes specific to thymus. (A) Gene ontology and Reactome pathway analysis of 70 p53 target genes activated only in irradiated thymus shows no significant pathway enrichment. (B) Irradiated p53 ChIP-seq at the *F8a* locus and *F8a* expression by RNA-seq in liver, spleen, and thymus. (C) Irradiated p53 ChIP-seq at the *Tmem131l* locus and *Tmem131l* expression by RNA-seq in liver, spleen, and thymus. (D) Irradiated p53 ChIP-seq at the *L3mbtl3* locus and *L3mbtl3* expression by RNA-seq in liver, spleen, and thymus. (E) Irradiated p53 ChIP-seq at the *Slfn2* locus and *Slfn2* expression by RNA-seq in liver, spleen, and thymus. (F) Irradiated p53 ChIP-seq at the *Ndrg3* locus and *Ndrg3* expression by RNA-seq in liver, spleen, and thymus. (G) Irradiated p53 ChIP-seq at the *Unc5cl* locus and *Unc5cl* expression by RNA-seq in liver, spleen, and thymus. (H) Irradiated p53 ChIP-seq at the *Axdnd1* locus and *Axdnd1* expression by RNA-seq in liver, spleen, and thymus.

**Supplementary Figure 11:**
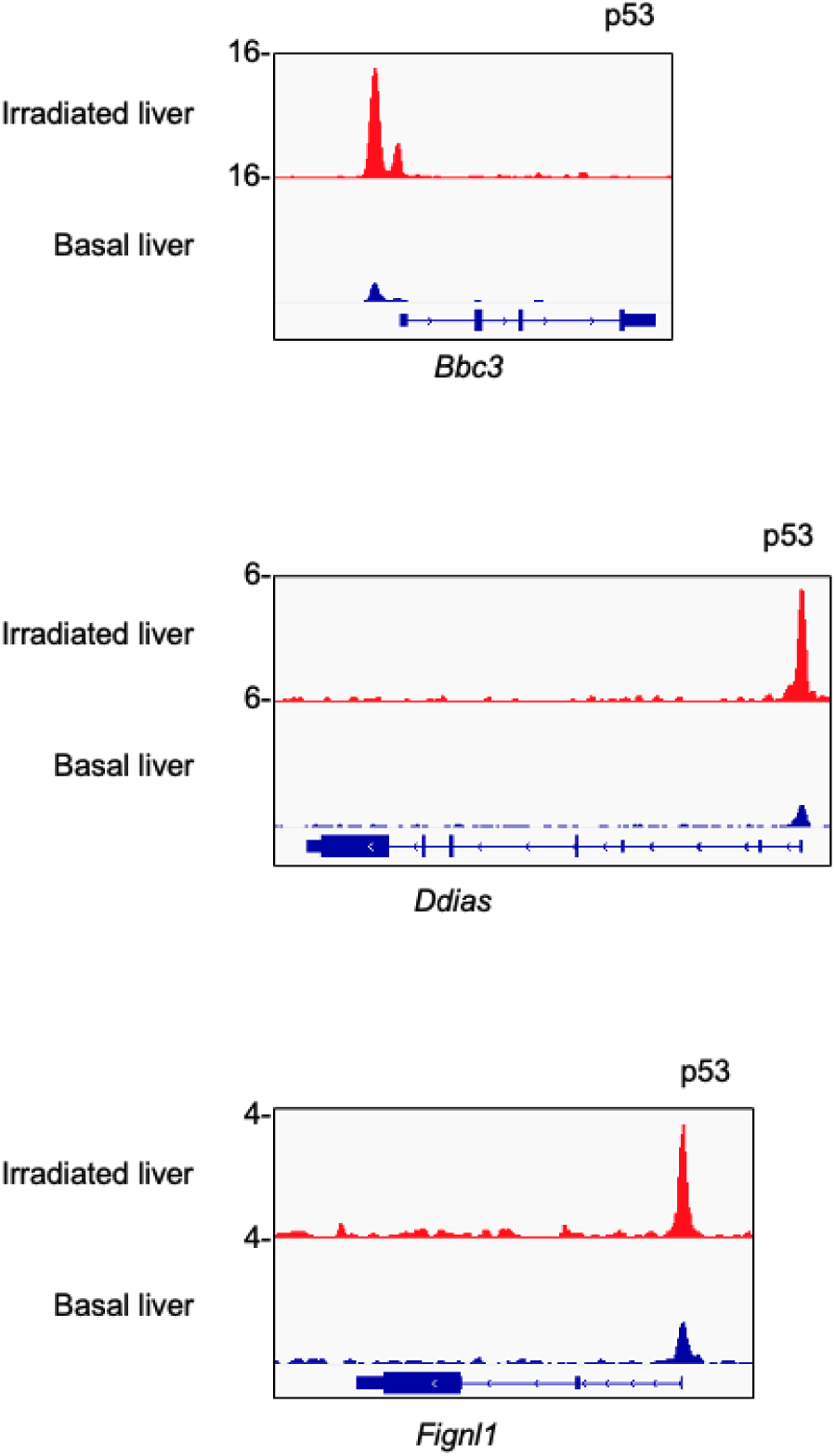
ChIP-seq tracks of p53 at basal target genes, under basal and irradiated conditions. p53 ChIP-seq tracks at the loci of *Bbc3, Ddias,* and *Fignl1* in irradiated versus basal tissues.

**Supplementary Figure 12.**
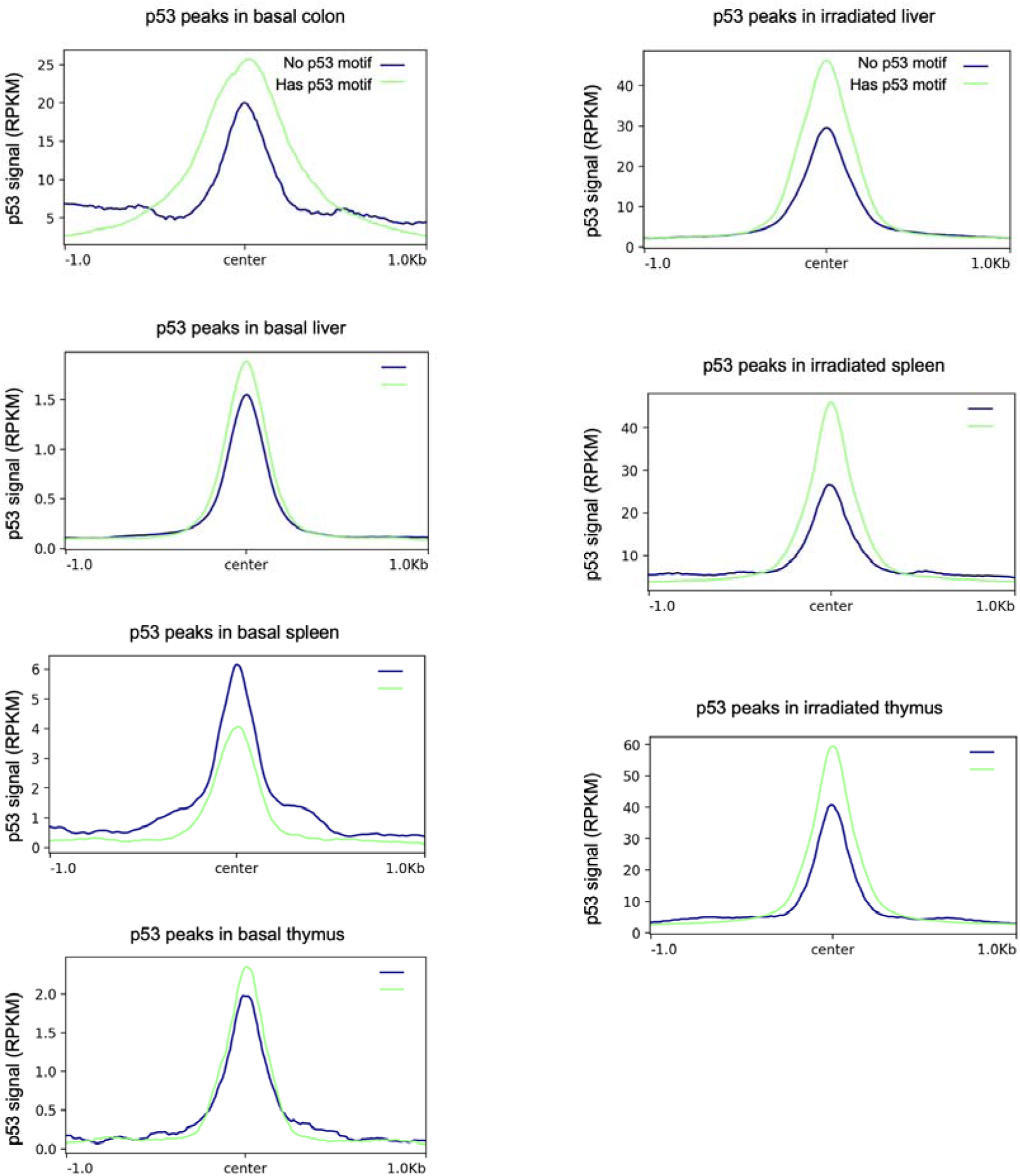
p53 signal at peaks with versus without a p53 consensus sequence. Summary plots of p53 ChIP-seq signal in basal and irradiated tissues. Green signifies p53 peaks containing a consensus binding sequence; blue signifies p53 peaks without a consensus binding sequence.

**Supplementary Figure 13.**
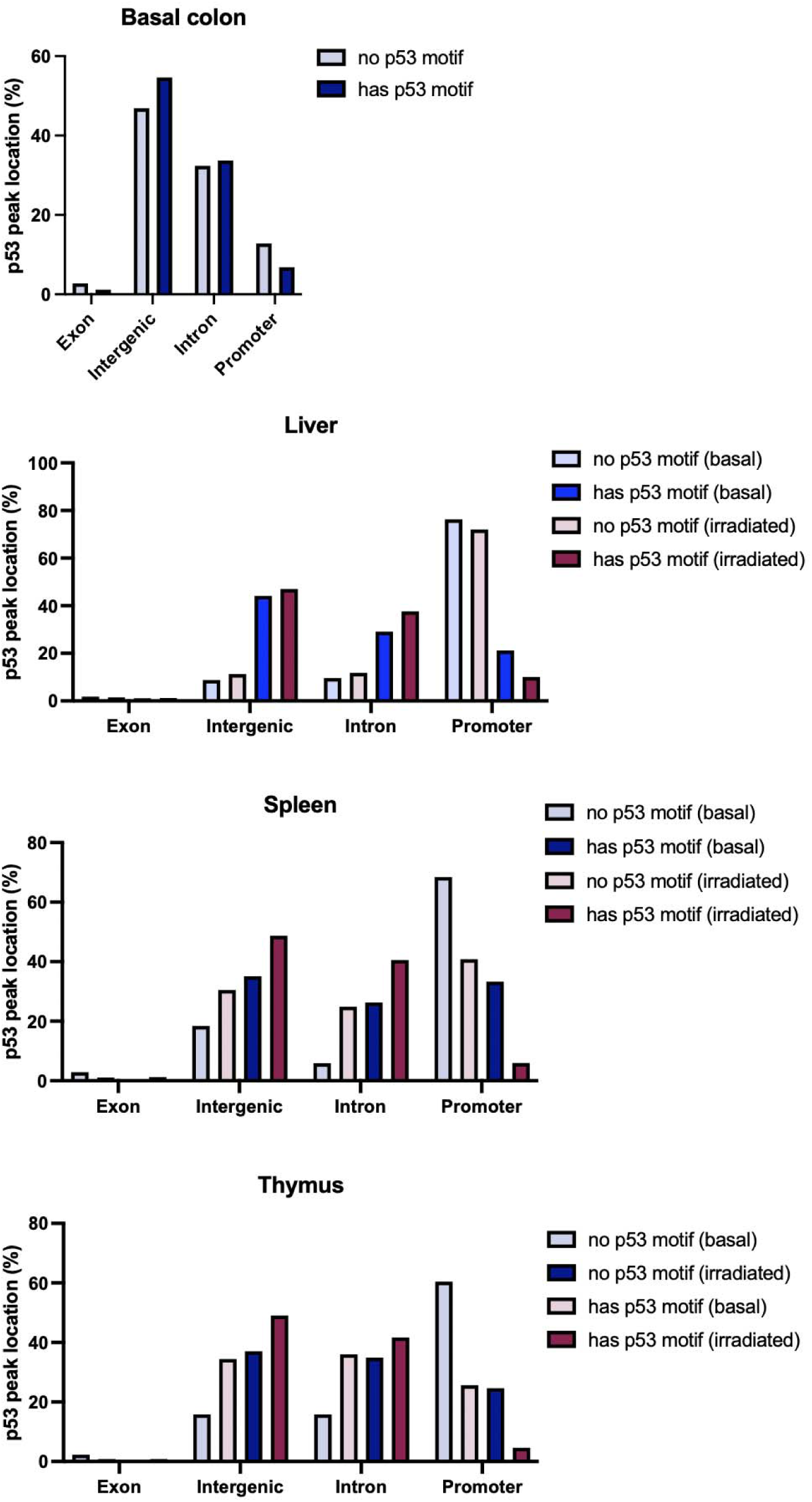
Location of p53 peaks depending on whether they contain a p53 consensus sequence. Location of p53 peaks in basal and irradiated tissues. p53 peaks are further divided between whether a consensus binding sequence is present.

**Supplementary Figure 14.**
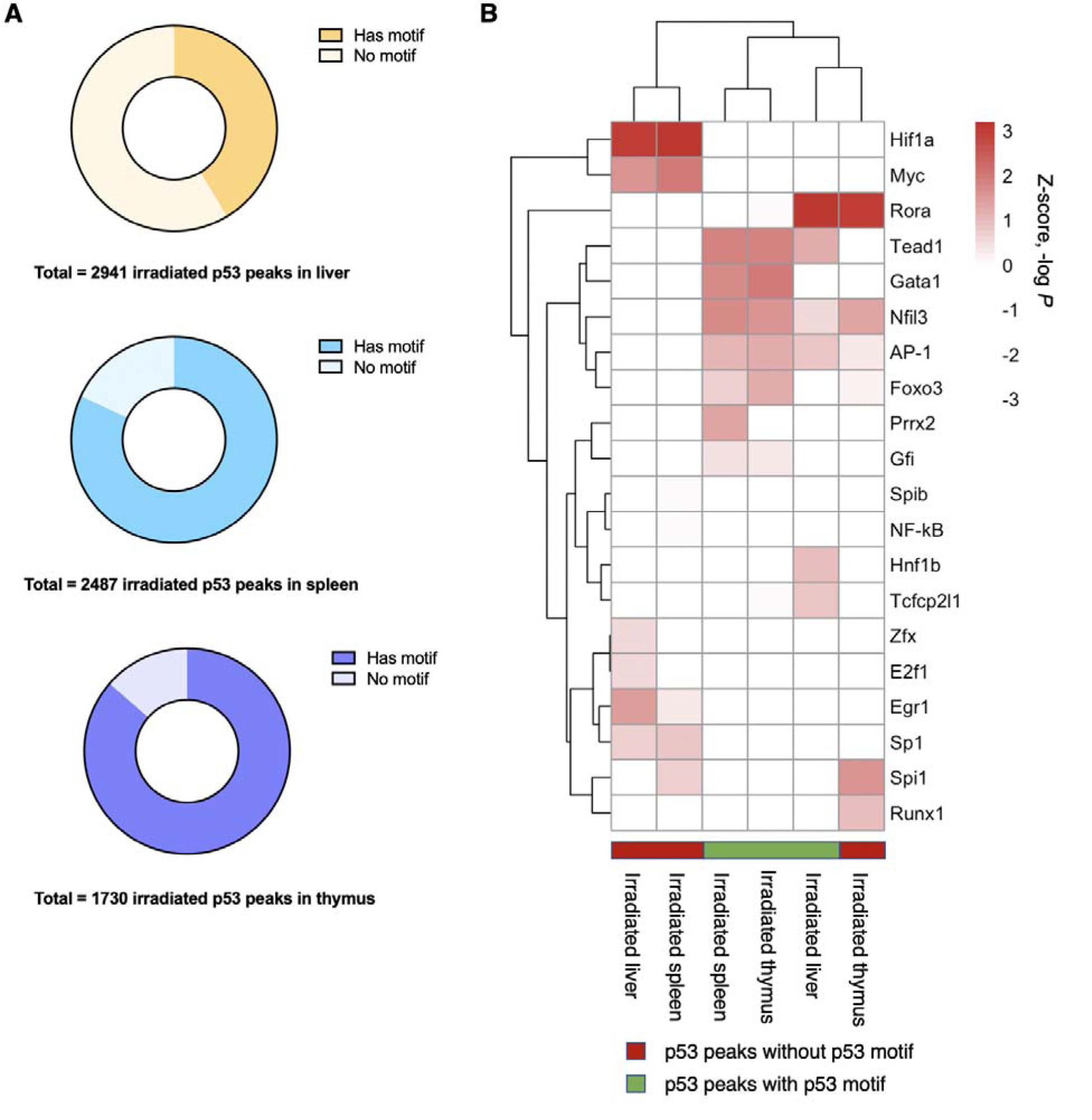
Transcription factor motif analysis of irradiated p53 peak depending on whether they contain a p53 consensus sequence. (A) Pie chart of irradiated p53 peaks depending on whether a p53 consensus binding sequence i present. (B) Heatmap and unsupervised hierarchical clustering of transcription factor motifs (by significance of motif enrichment) among irradiated tissues. Columns are further divided by whether a p53 consensus binding sequence is present. Only expressed transcription factors are included—a non-expressed transcription factor (TPM < 1) is assigned a *P* = 1.

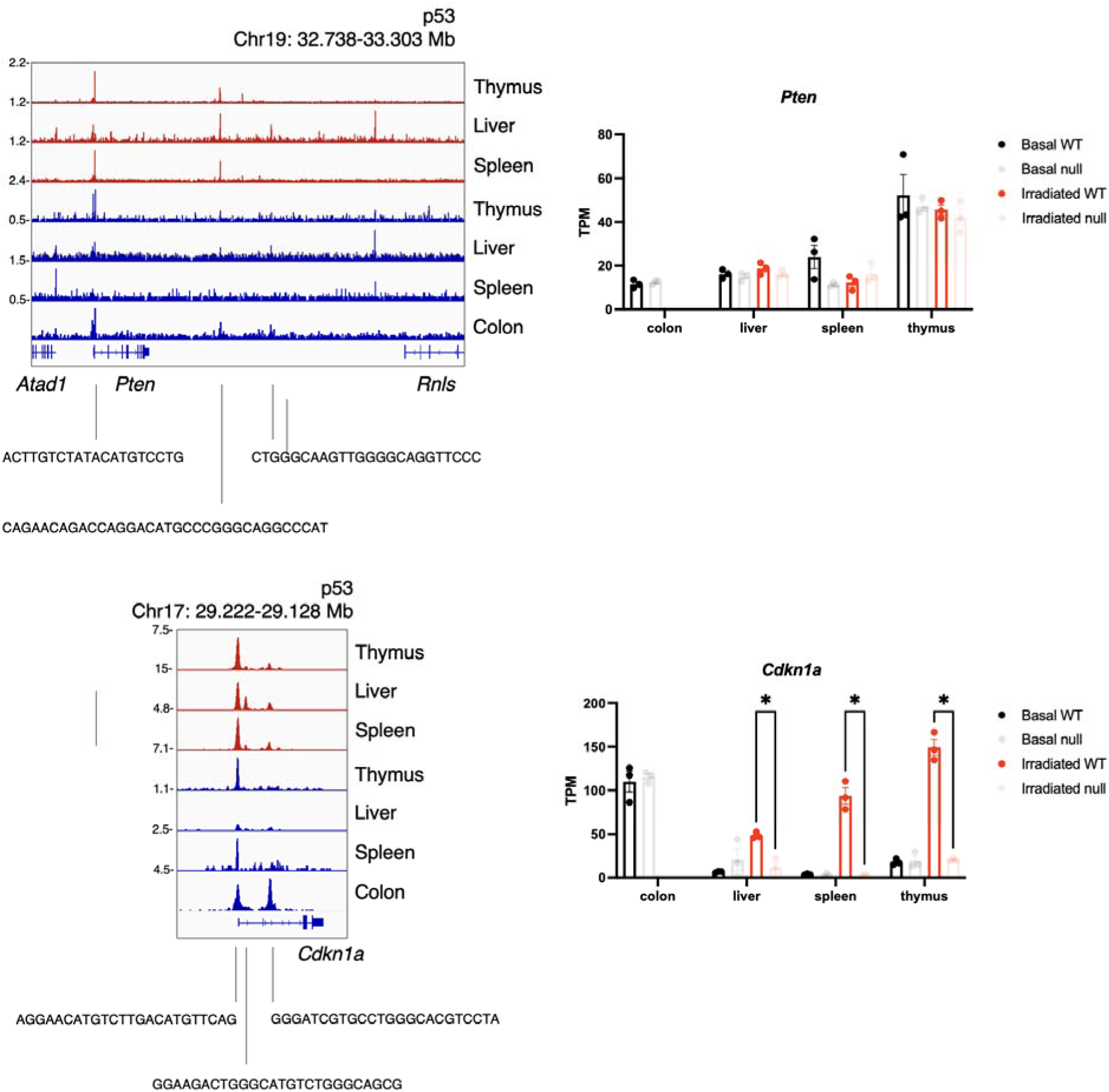
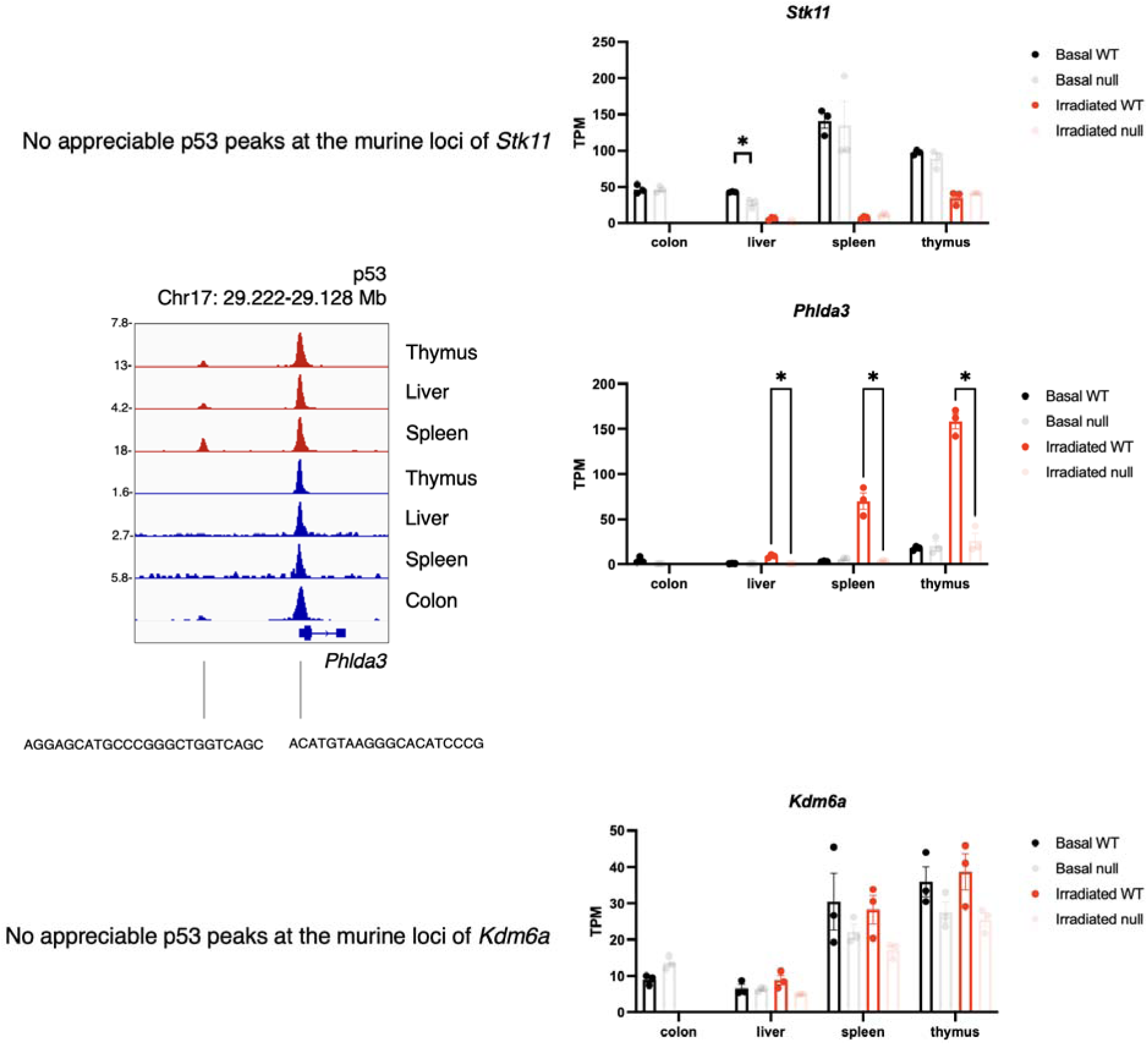
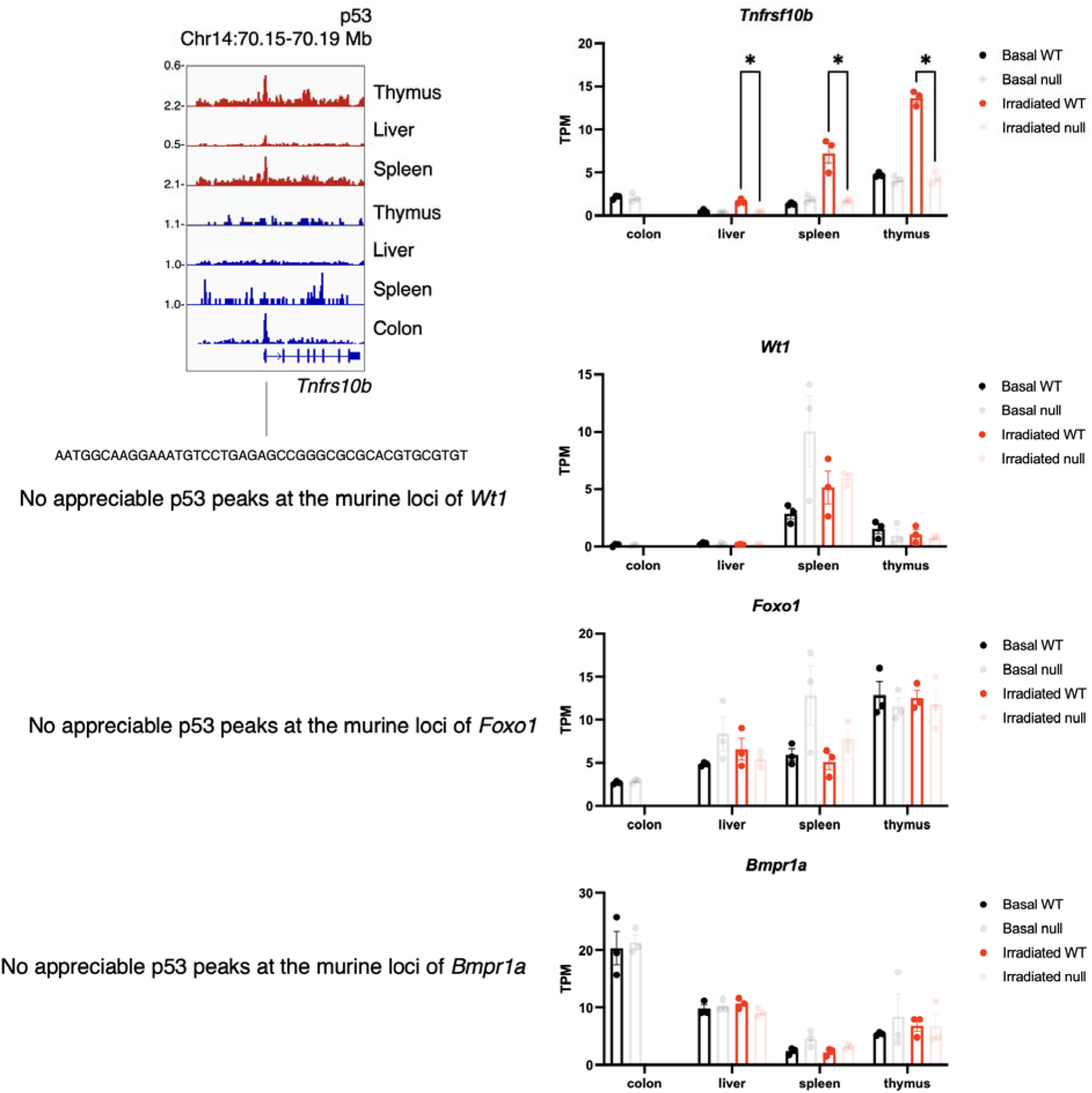
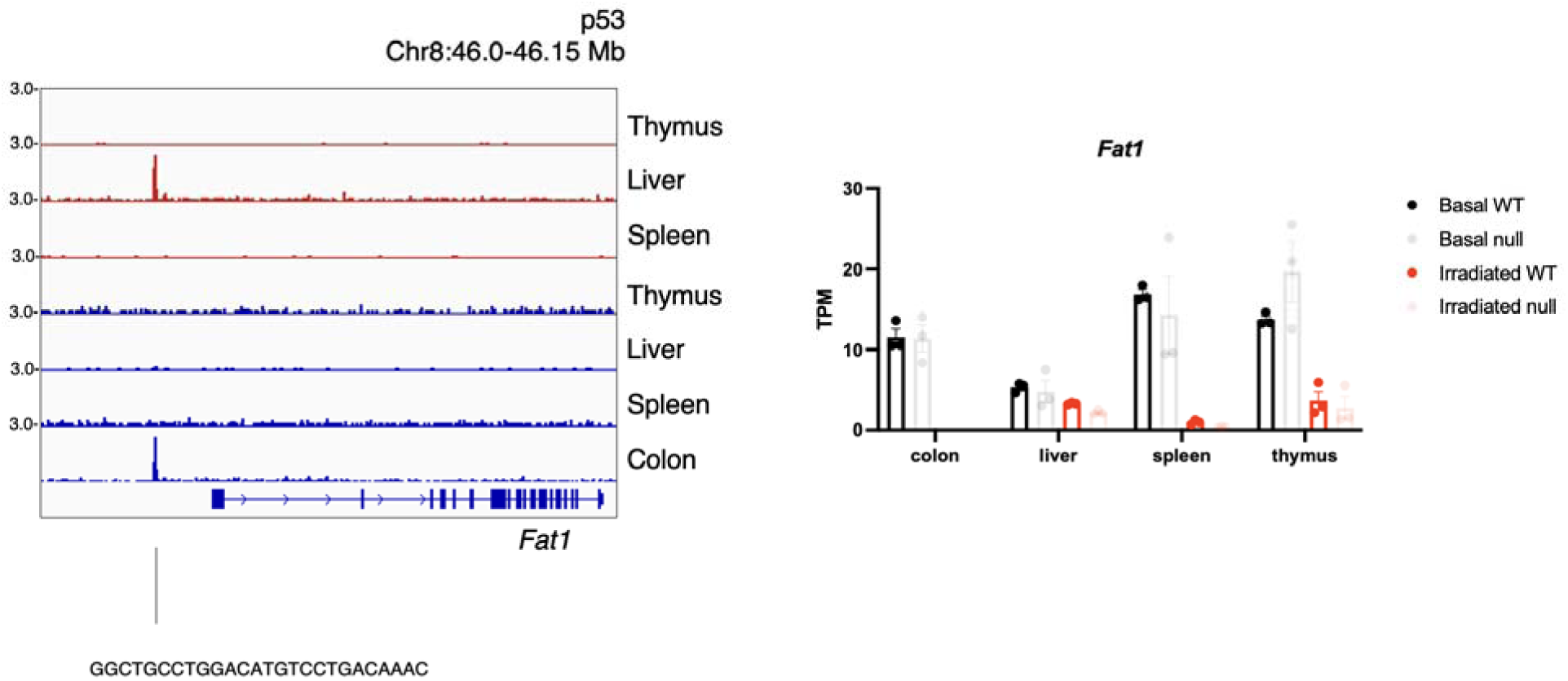

