## Supplemental Figures for "Basal p53 maintains a distinct transcriptional program from irradiated p53 in tissue, including tumor suppressors"

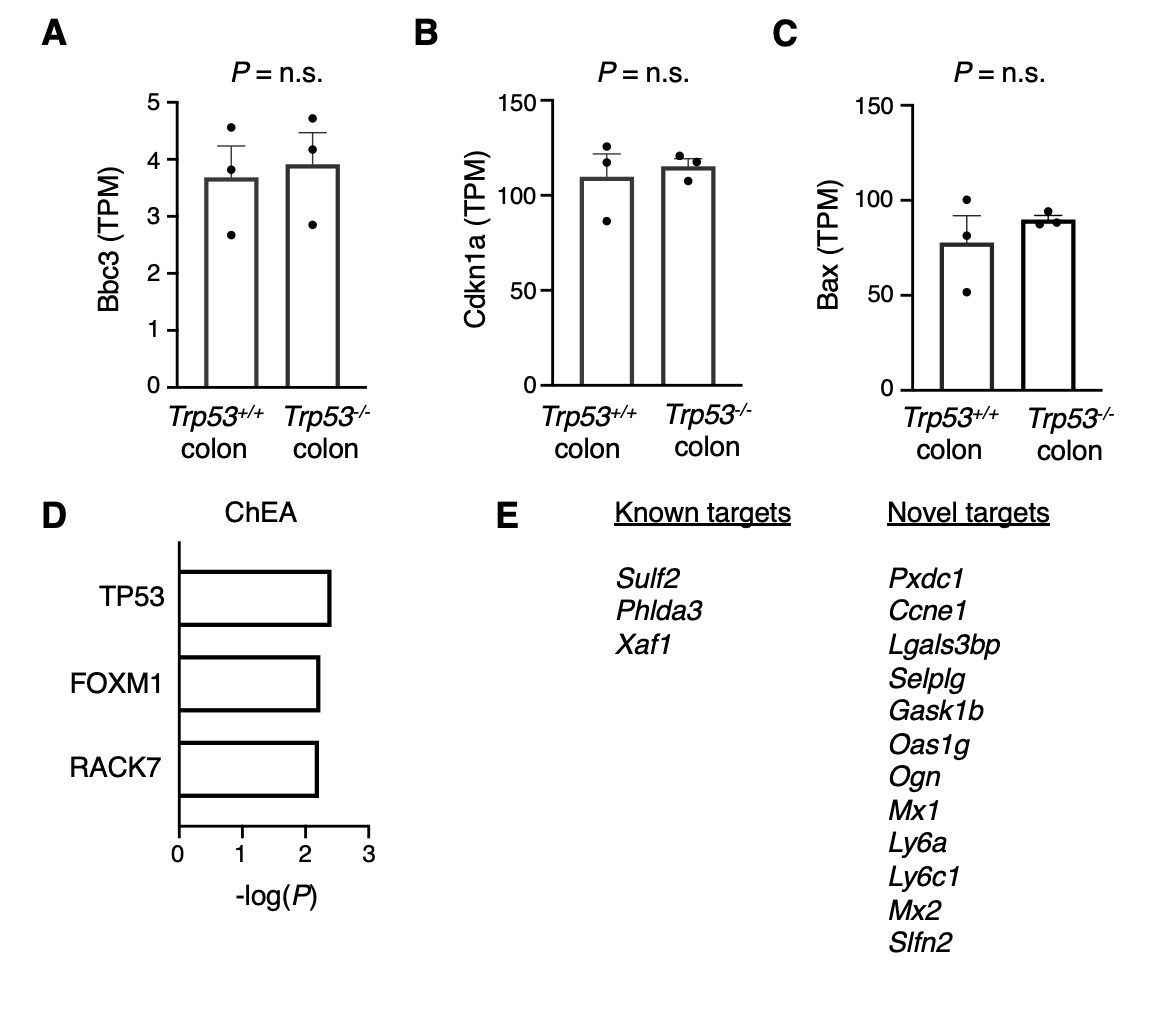

**Supplementary Figure 1. Basal p53 occupancy in physiologic colon identifies known and novel targets.**

(A-C) Gene expression of known p53 target genes in physiologic murine colon epithelium via RNA-seq between *n* = 3 WT and *n* = 3 null mice. (D) ChEA transcription factor analysis of identified p53 targets. (E) Bona fide basal p53 target genes in physiologic murine colon epithelium, defined by p53 occupancy by ChIP-seq and differential expression between WT and null colon by >1 log_2_ fold change.

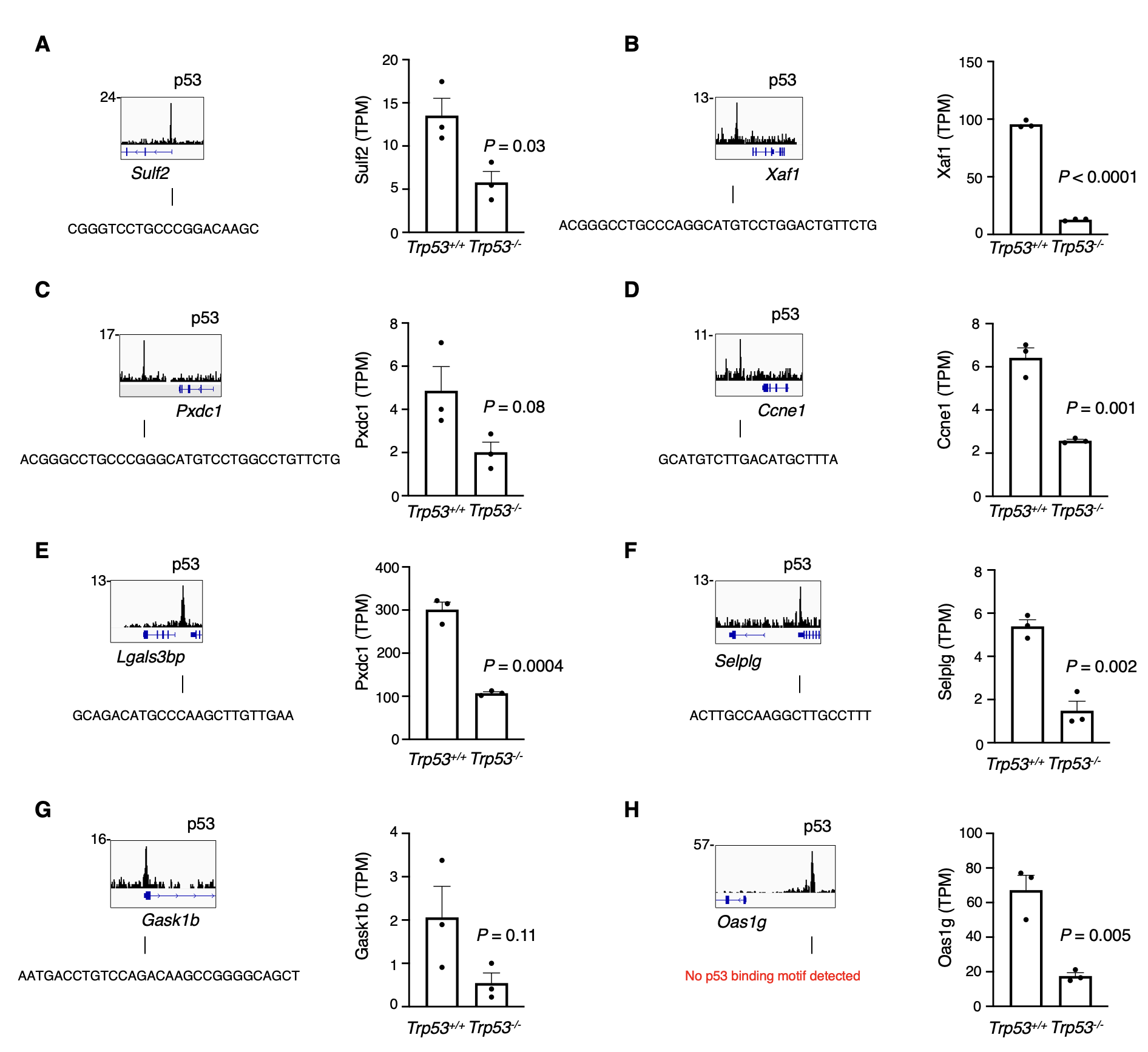

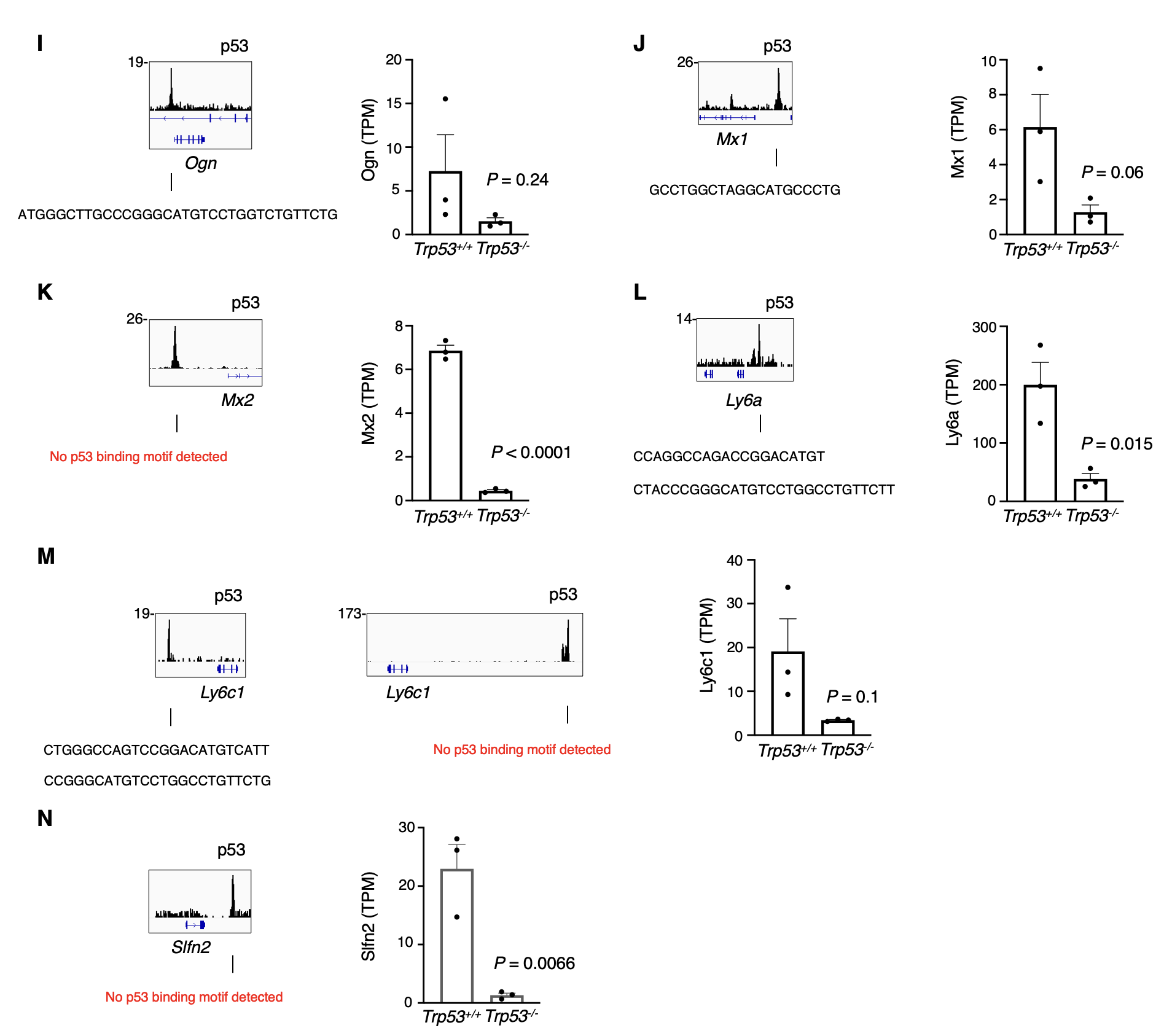

**Supplementary Figure 2. Basal p53 target genes in physiologic murine colon epithelium.**

p53 ChIP-seq tracks at basal p53 target genes from Figure 1. DNA sequences under called peaks were subject to TRAP motif analysis for *de novo* discovery of p53 binding motifs. Expression of target genes by RNA-seq between p53 WT (*n* = 3) or null (*n* = 3) murine colon epithelium.

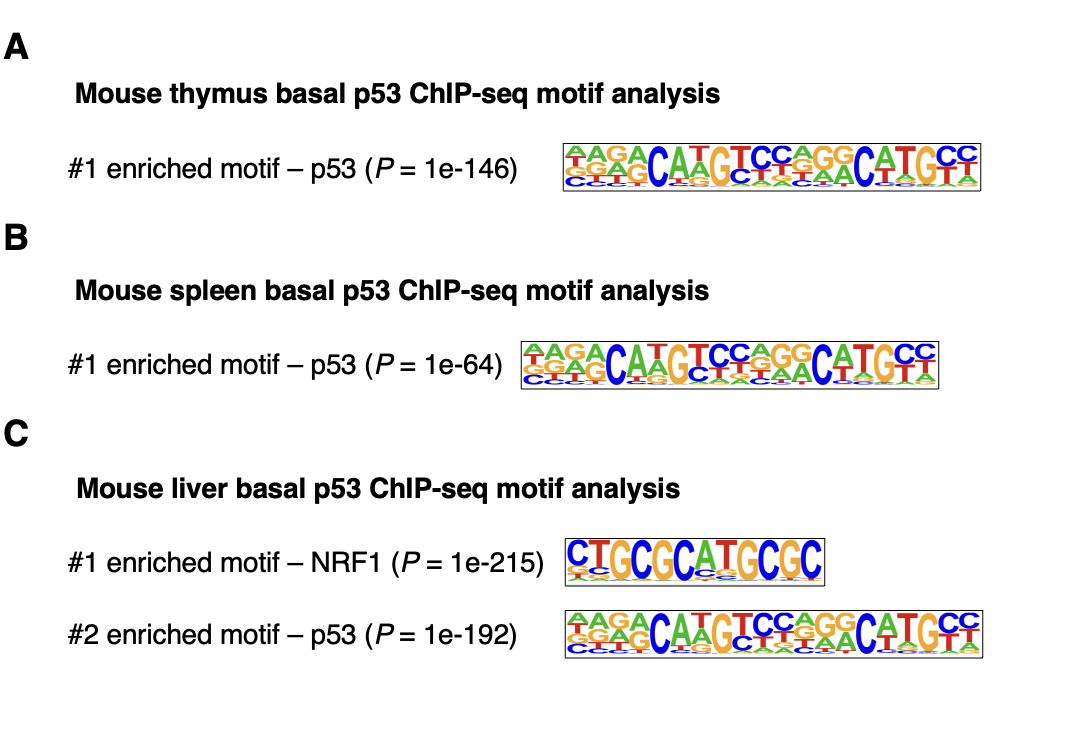

**Supplementary Figure 3. Basal p53 peaks are enriched for its consensus binding sequence.**

HOMER *de novo* transcription factor motif discovery of called basal p53 peaks in murine (A) thymus, (B) spleen, or (C) liver.

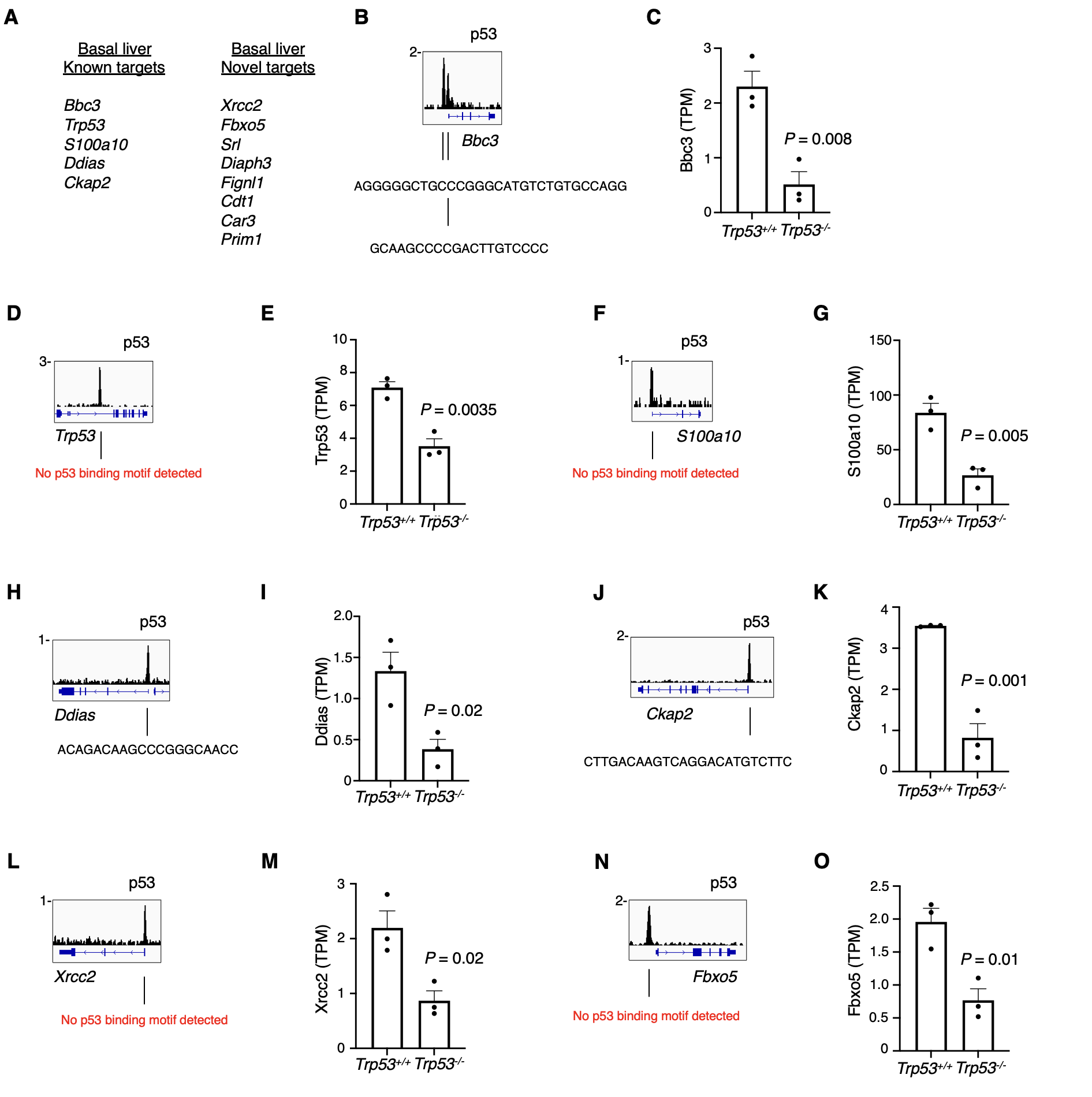

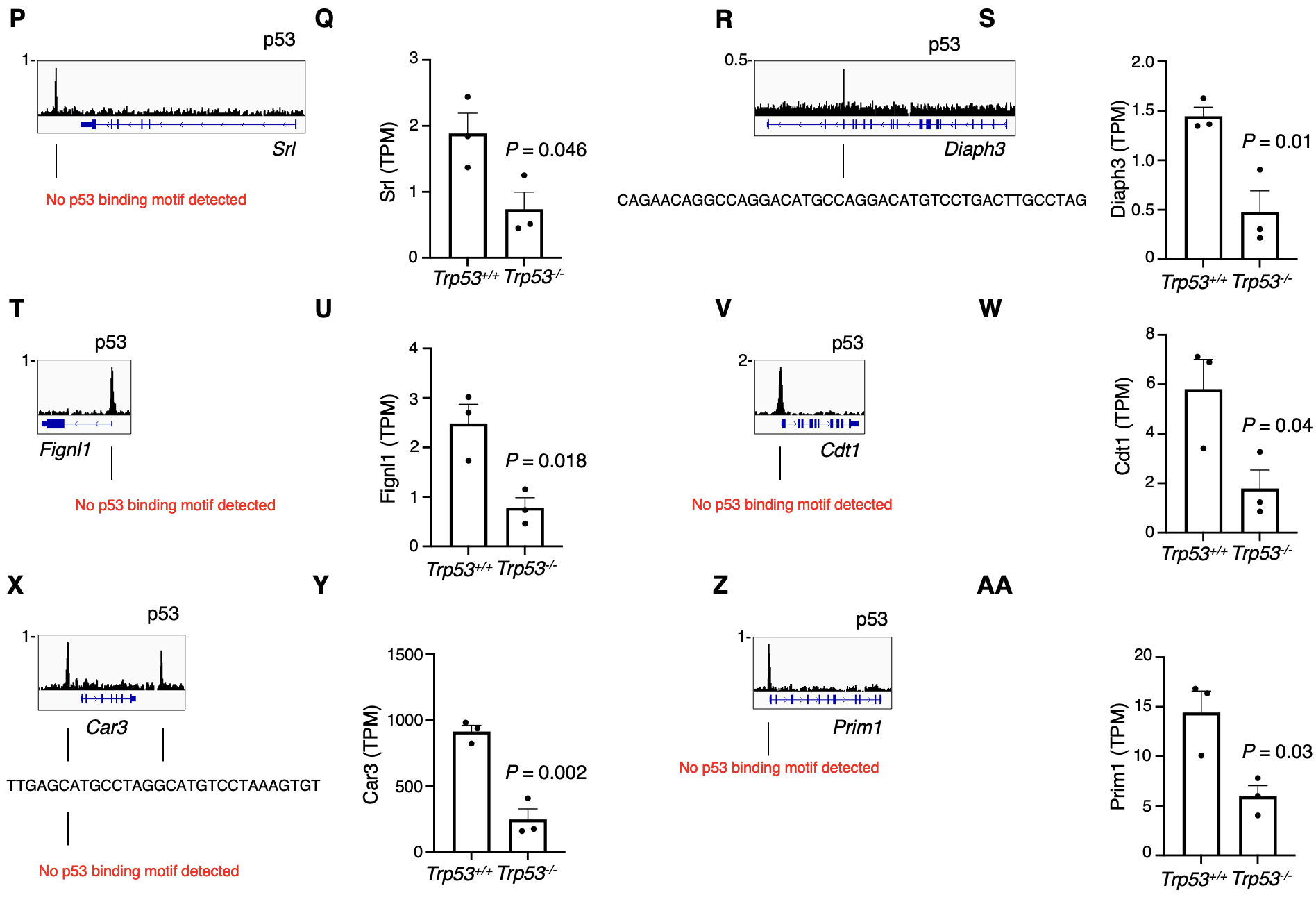

**Supplementary Figure 4. Basal p53 target genes in physiologic murine liver.**

p53 ChIP-seq tracks at basal p53 target genes identified in murine liver listed in panel (A). DNA sequences under called peaks were subject to TRAP motif analysis for *de novo* discovery of p53 binding motifs. Expression of target genes by RNA-seq between p53 WT (*n* = 3) or null (*n* = 3) murine liver.

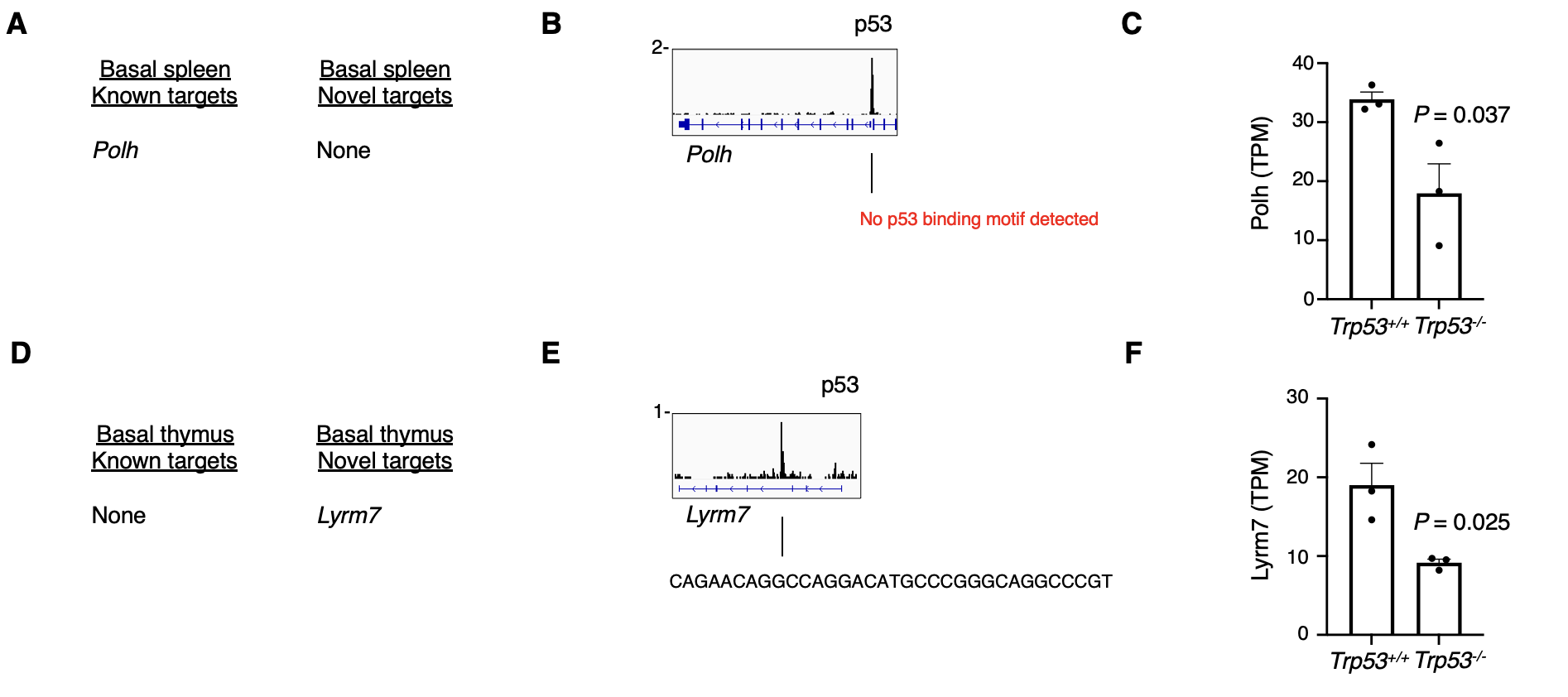

**Supplementary Figure 5. Basal p53 target genes in physiologic murine spleen and thymus.**

(A) p53 ChIP-seq tracks at basal p53 target genes identified in murine spleen. (B) DNA sequences under called peaks were subject to TRAP motif analysis for *de novo* discovery of p53 binding motifs. (C) Expression of target genes by RNA-seq between p53 WT (*n* = 3) or null (*n* = 3) murine spleen. (D) p53 ChIP-seq tracks at basal p53 target genes identified in murine thymus. (E) DNA sequences under called peaks were subject to TRAP motif analysis for *de novo* discovery of p53 binding motifs. (F) Expression of target genes by RNA-seq between p53 WT (*n* = 3) or null (*n* = 3) murine thymus.

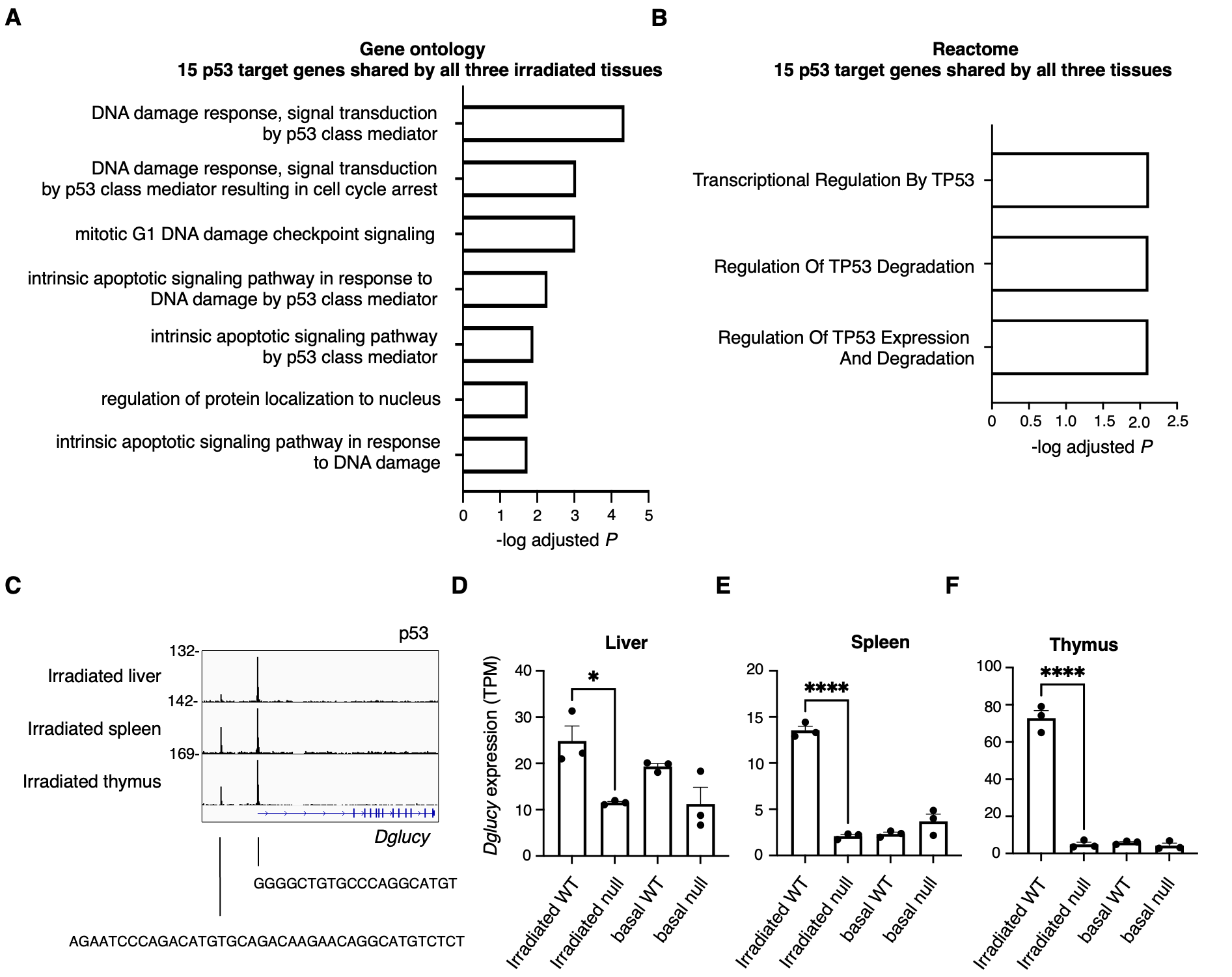

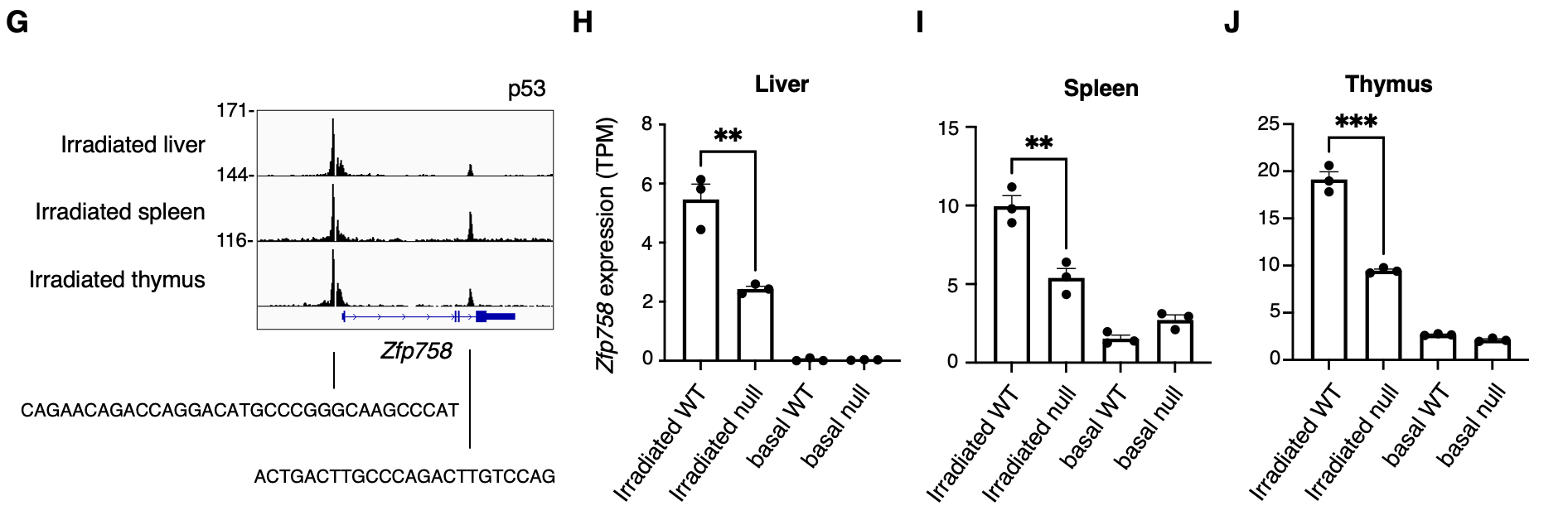

**Supplementary Figure 6. Irradiated p53 target genes activated in all three tissues.**

(A) Gene ontology and (B) Reactome pathway analysis of 15 p53 target genes activated in irradiated spleen, thymus, and liver. (C) Irradiated p53 ChIP-seq at the *Dglucy* locus. (D-F) *Dglucy* expression by RNA-seq in liver, spleen, and thymus. (G) Irradiated p53 ChIP-seq at the *Zfp958* locus. (H-J) *Zfp958* expression by RNA-seq in liver, spleen, and thymus.

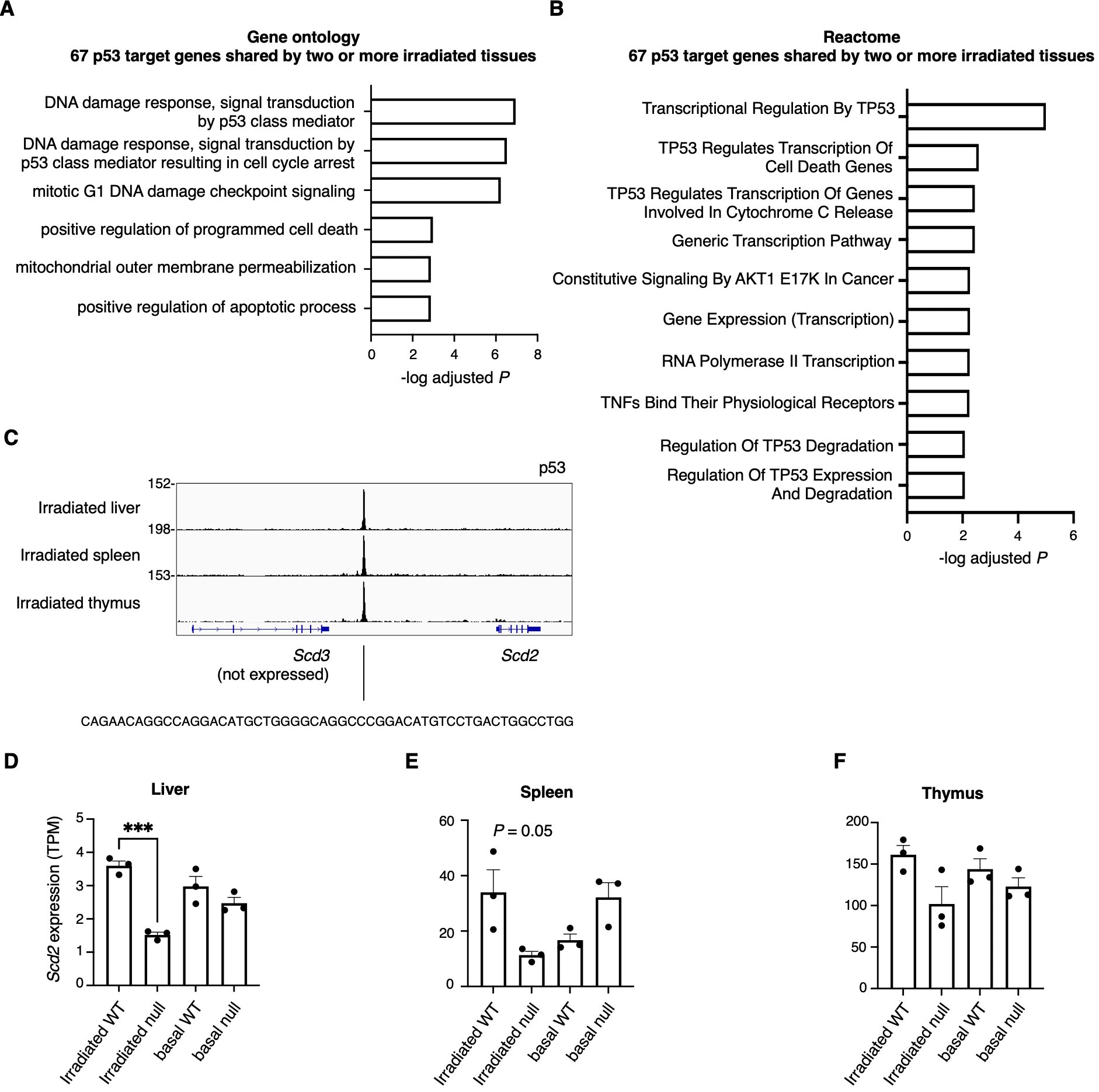

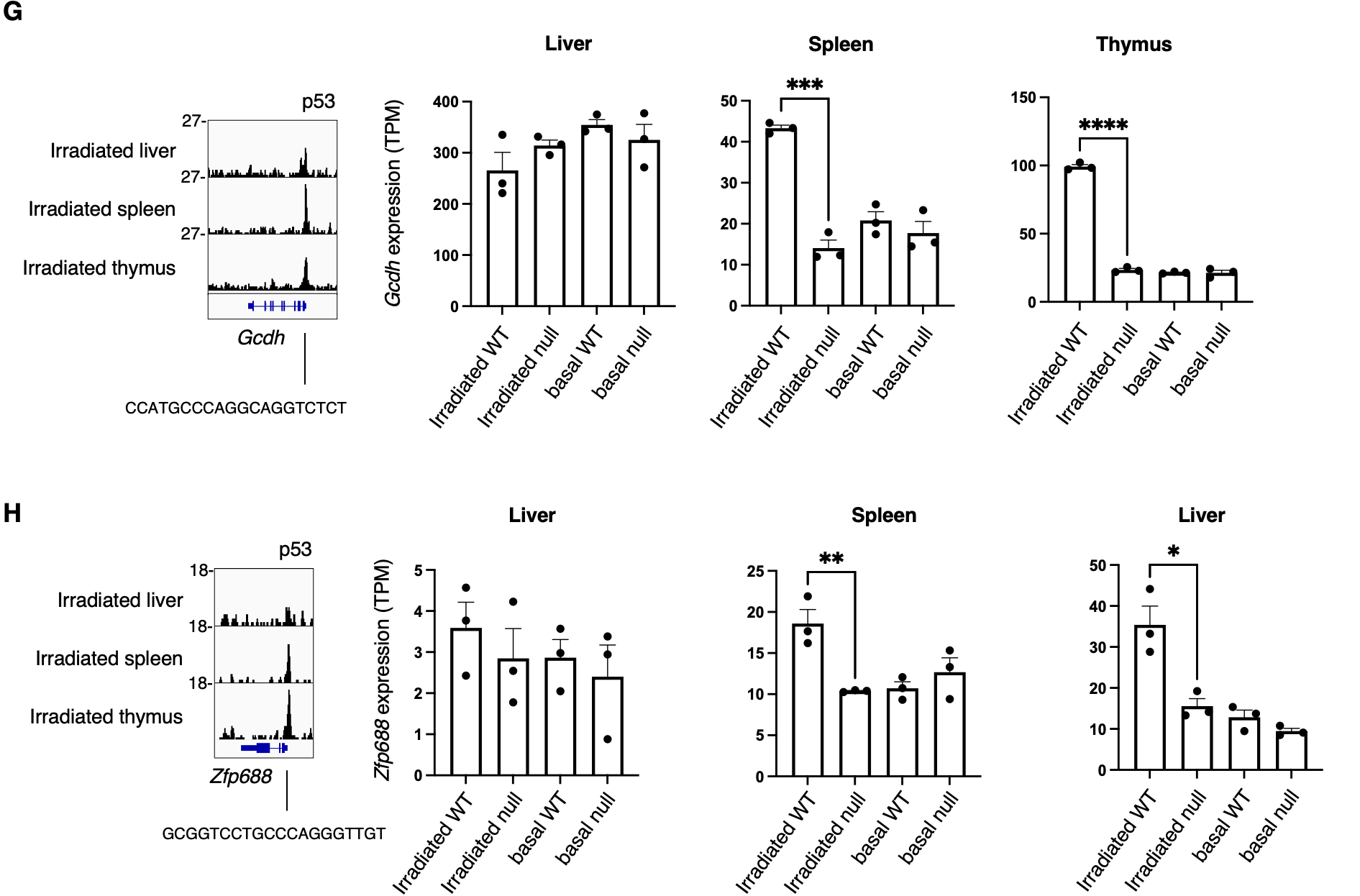

**Supplementary Figure 7. Irradiated p53 target genes activated in two or more tissues.**

(A) Gene ontology and (B) Reactome pathway analysis of 67 p53 target genes activated in two or more irradiated tissues. (C) Irradiated p53 ChIP-seq track at the *Scd2* locus. (D-F) *Scd2* expression by RNA-seq in liver, spleen, and thymus. (G) Irradiated p53 ChIP-seq track at the *Gcdh* locus, with RNA-seq gene expression. (H) Irradiated p53 ChIP-seq track at the *Zfp688* locus with RNA-seq gene expression.

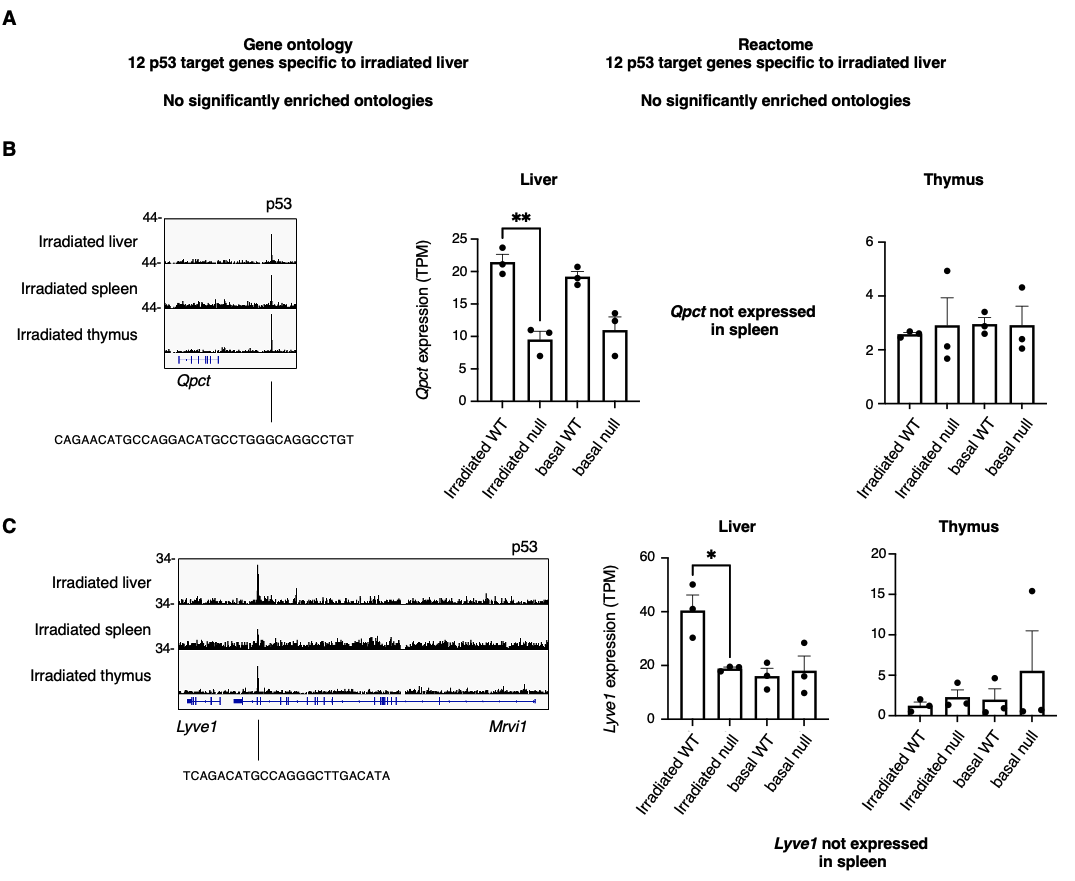

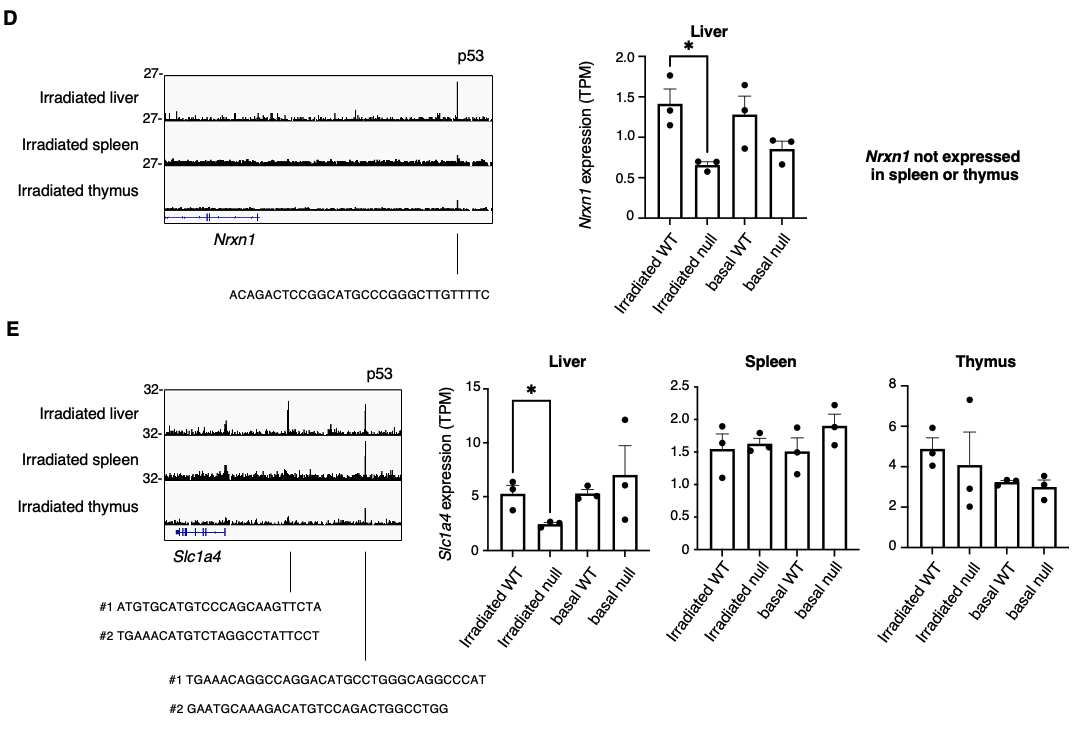

**Supplementary Figure 8. Irradiated p53 target genes specific to liver.**

(A) Gene ontology and Reactome pathway analysis of 12 p53 target genes activated inly in irradiated liver showed no significant enrichment. (B) Irradiated p53 ChIP-seq at the *Qpct* locus and *Qpct* expression by RNA-seq in liver, spleen, and thymus. (C) Irradiated p53 ChIP-seq at the *Lyve1* locus and *Lyve1* expression by RNA-seq in liver, spleen, and thymus. (D) Irradiated p53 ChIP-seq at the *Nrxn1* locus and *Nrxn1* expression by RNA-seq in liver, spleen, and thymus. (E) Irradiated p53 ChIP-seq at the *Slc1a4* locus and *Slc1a4* expression by RNA-seq in liver, spleen, and thymus.

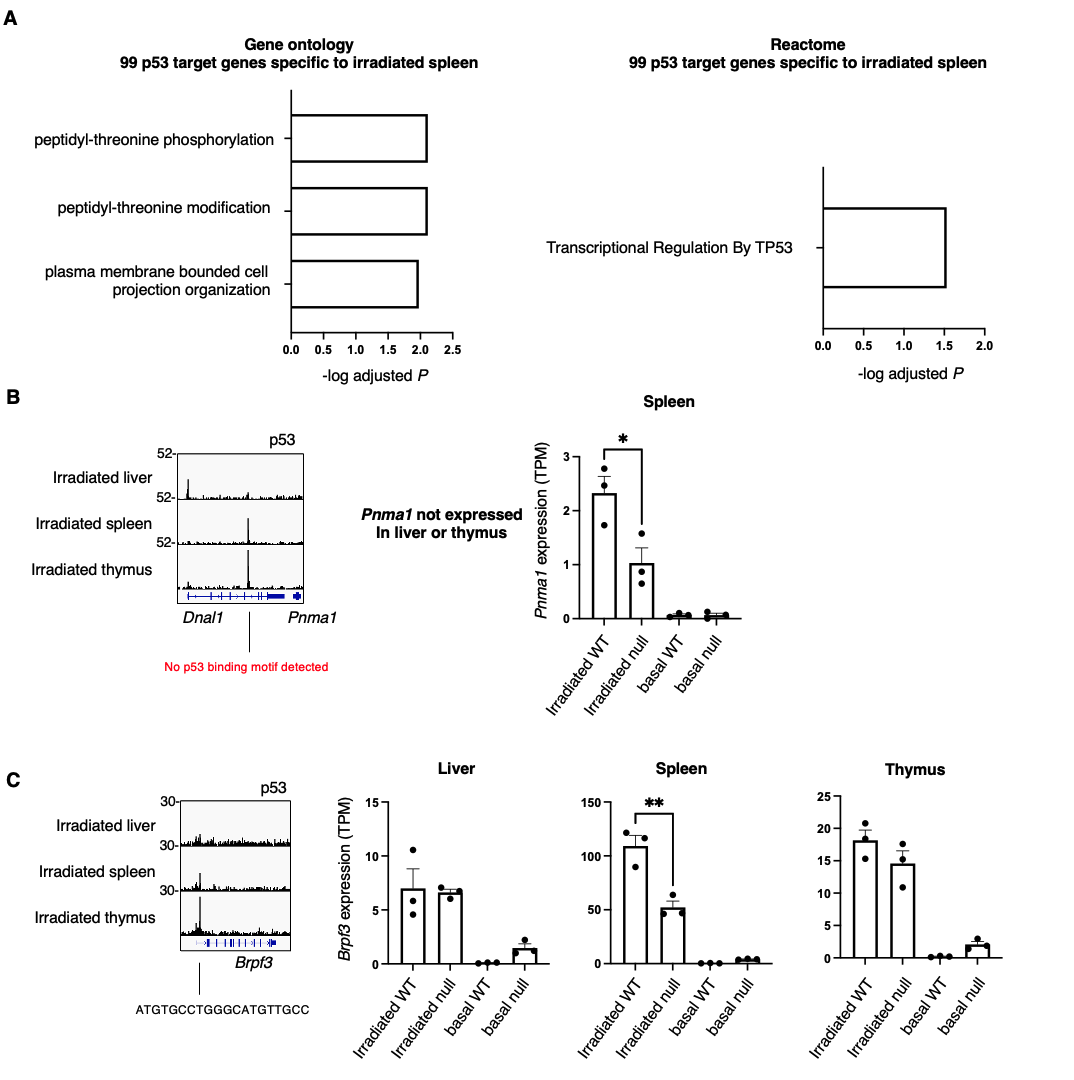

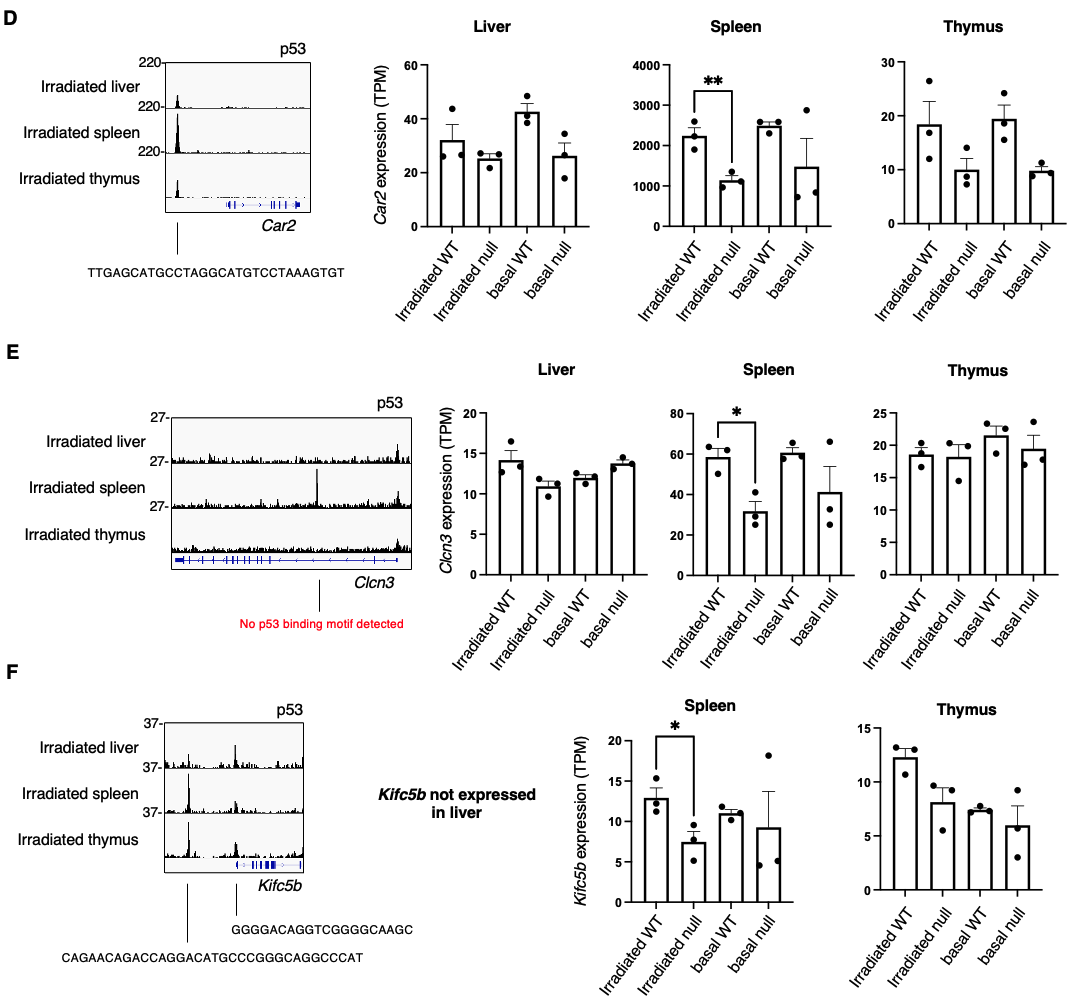

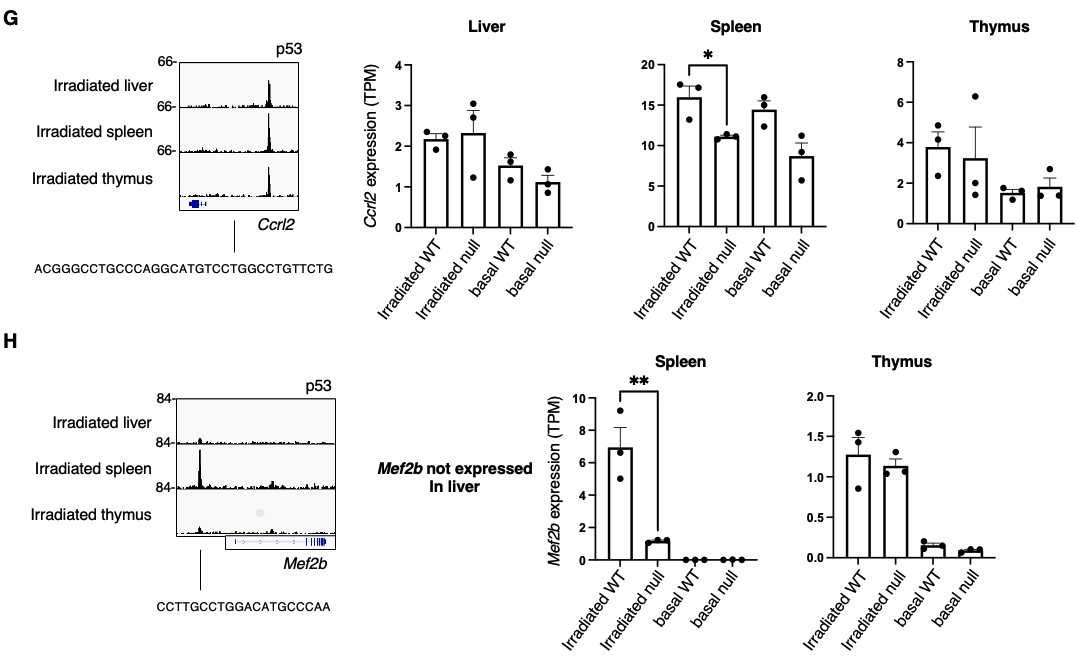
**Supplementary Figure 9. Irradiated p53 target genes specific to spleen.**

(A) Gene ontology and Reactome pathway analysis of 99 p53 target genes activated only in irradiated spleen. (B) Irradiated p53 ChIP-seq at the *Pnma1* locus and *Pnma1* expression by RNA-seq in liver, spleen, and thymus. (C) Irradiated p53 ChIP-seq at the *Brpf3* locus and *Brpf3* expression by RNA-seq in liver, spleen, and thymus. (D) Irradiated p53 ChIP-seq at the *Car2* locus and *Car2* expression by RNA-seq in liver, spleen, and thymus. (E) Irradiated p53 ChIP-seq at the *Clcn3* locus and *Clcn3* expression by RNA-seq in liver, spleen, and thymus. (F) Irradiated p53 ChIP-seq at the *Kifc5b* locus and *Kifc5b* expression by RNA-seq in liver, spleen, and thymus. (G) Irradiated p53 ChIP-seq at the *Ccrl2* locus and *Ccrl2* expression by RNA-seq in liver, spleen, and thymus. (H) Irradiated p53 ChIP-seq at the *Mef2b* locus and *Mef2b* expression by RNA-seq in liver, spleen, and thymus.

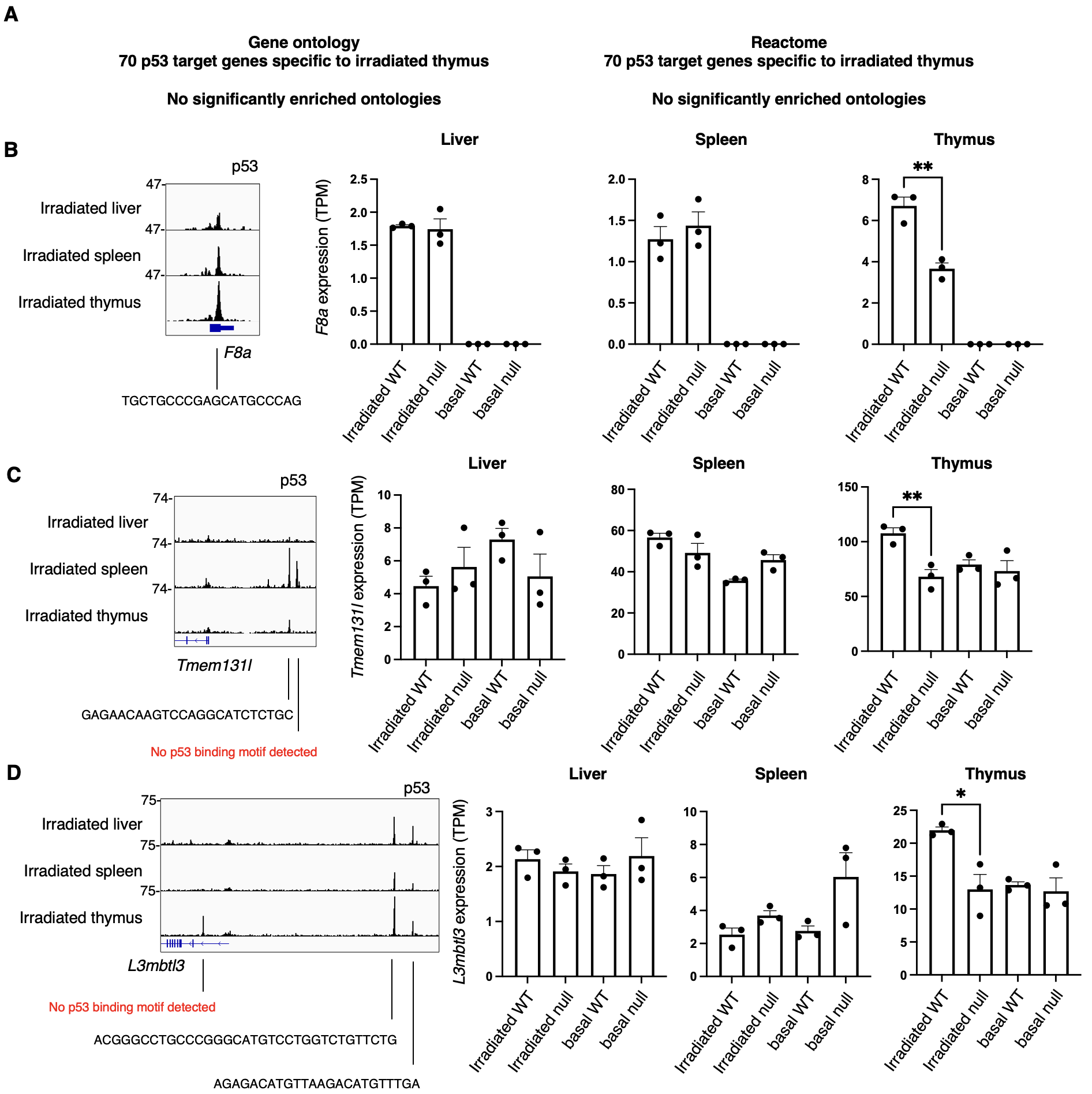

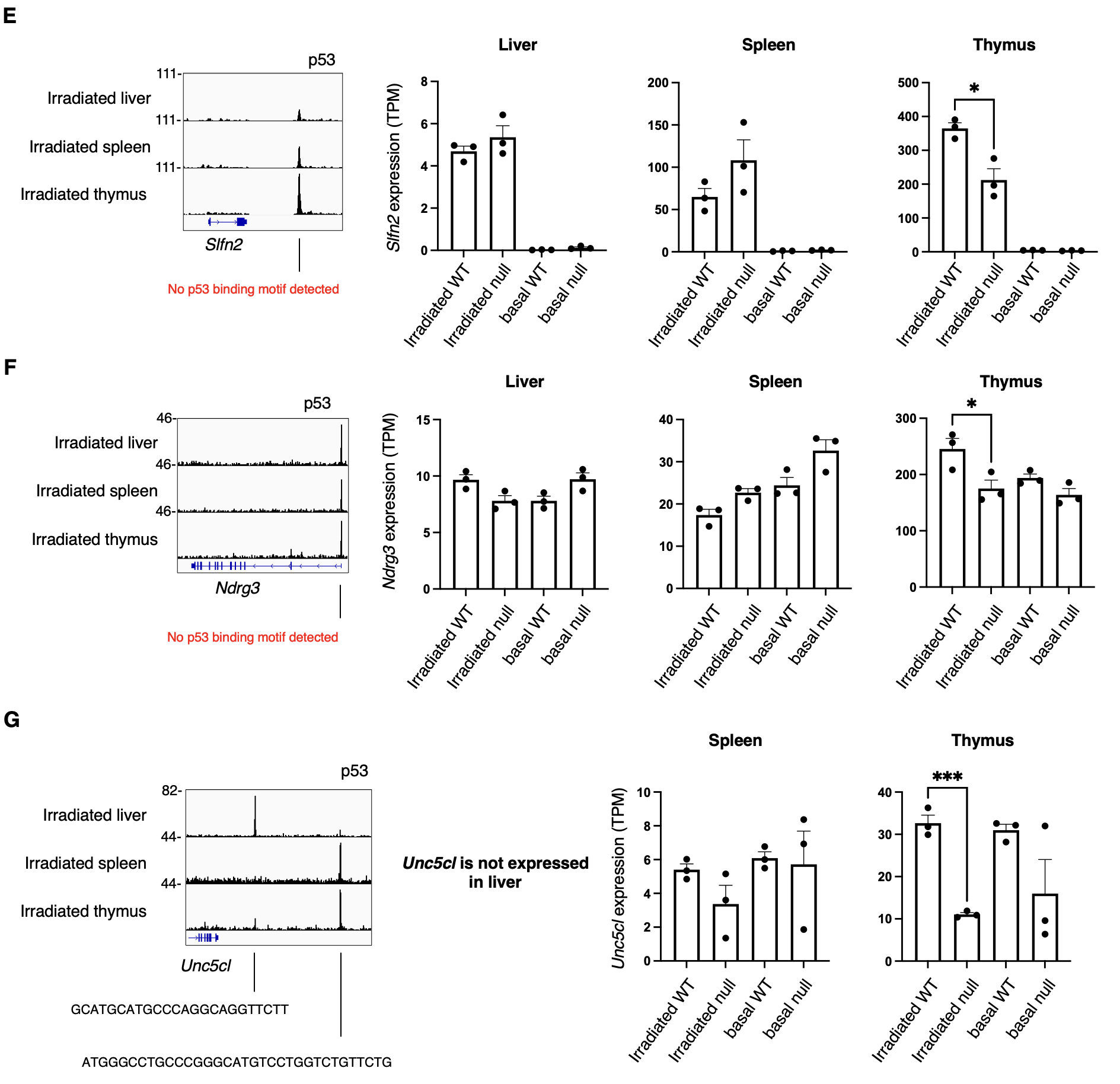

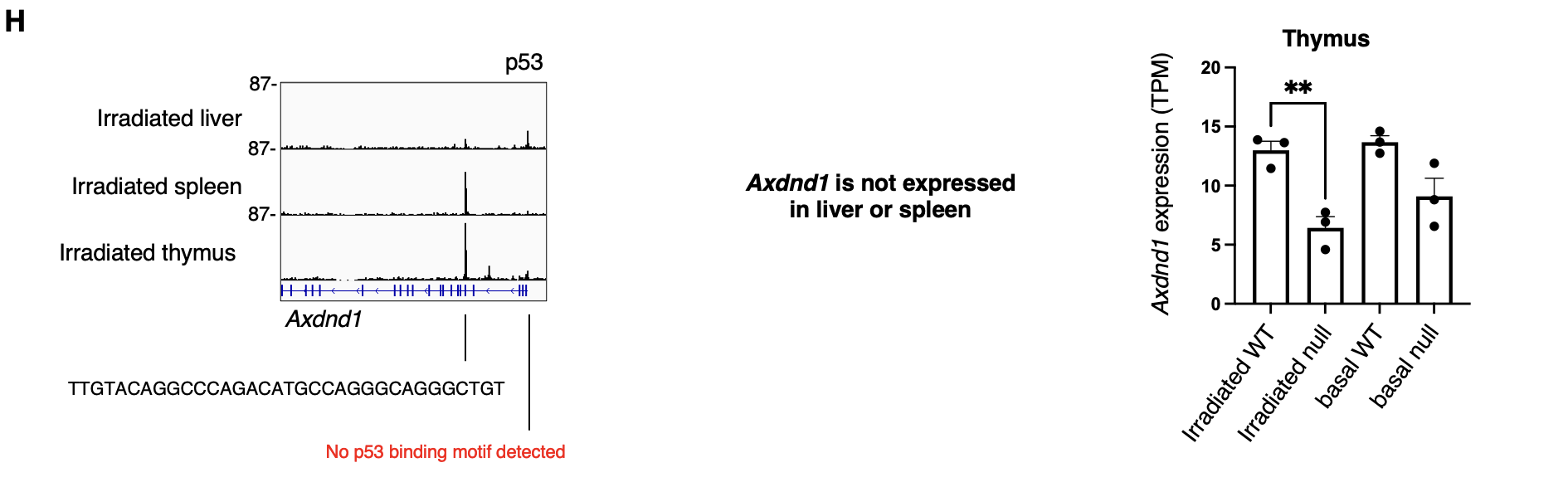

**Supplementary Figure 10. Irradiated p53 target genes specific to thymus.**

(A) Gene ontology and Reactome pathway analysis of 70 p53 target genes activated only in irradiated thymus shows no significant pathway enrichment. (B) Irradiated p53 ChIP-seq at the *F8a* locus and *F8a* expression by RNA-seq in liver, spleen, and thymus. (C) Irradiated p53 ChIP-seq at the *Tmem131l* locus and *Tmem131l* expression by RNA-seq in liver, spleen, and thymus. (D) Irradiated p53 ChIP-seq at the *L3mbtl3* locus and *L3mbtl3* expression by RNA-seq in liver, spleen, and thymus. (E) Irradiated p53 ChIP-seq at the *Slfn2* locus and *Slfn2* expression by RNA-seq in liver, spleen, and thymus. (F) Irradiated p53 ChIP-seq at the *Ndrg3* locus and *Ndrg3* expression by RNA-seq in liver, spleen, and thymus. (G) Irradiated p53 ChIP-seq at the *Unc5cl* locus and *Unc5cl* expression by RNA-seq in liver, spleen, and thymus. (H) Irradiated p53 ChIP-seq at the *Axdnd1* locus and *Axdnd1* expression by RNA-seq in liver, spleen, and thymus.

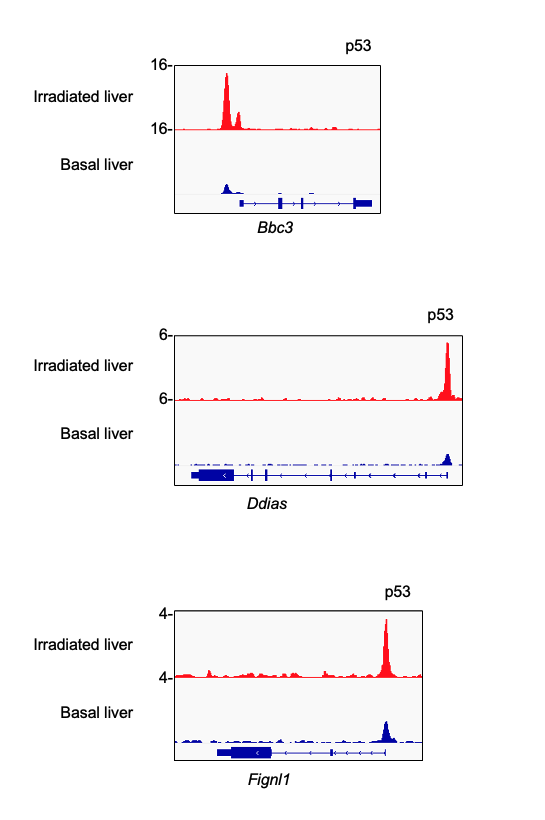

**Supplementary Figure 11: ChIP-seq tracks of p53 at basal target genes, under basal and irradiated conditions.**

p53 ChIP-seq tracks at the loci of *Bbc3, Ddias,* and *Fignl1* in irradiated versus basal tissues.

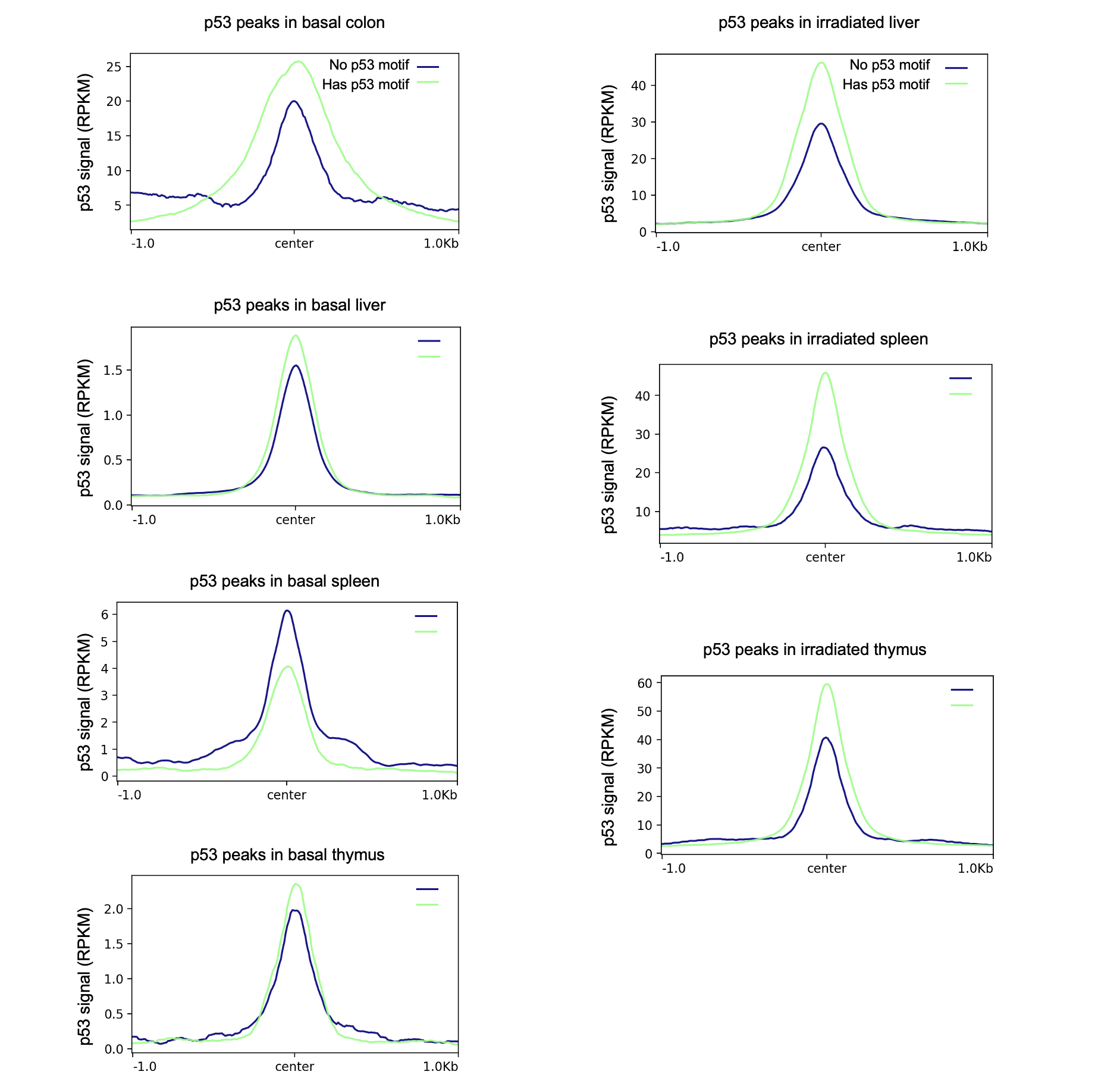

**Supplementary Figure 12. p53 signal at peaks with versus without a p53 consensus sequence.**

Summary plots of p53 ChIP-seq signal in basal and irradiated tissues. Green signifies p53 peaks containing a consensus binding sequence; blue signifies p53 peaks without a consensus binding sequence.

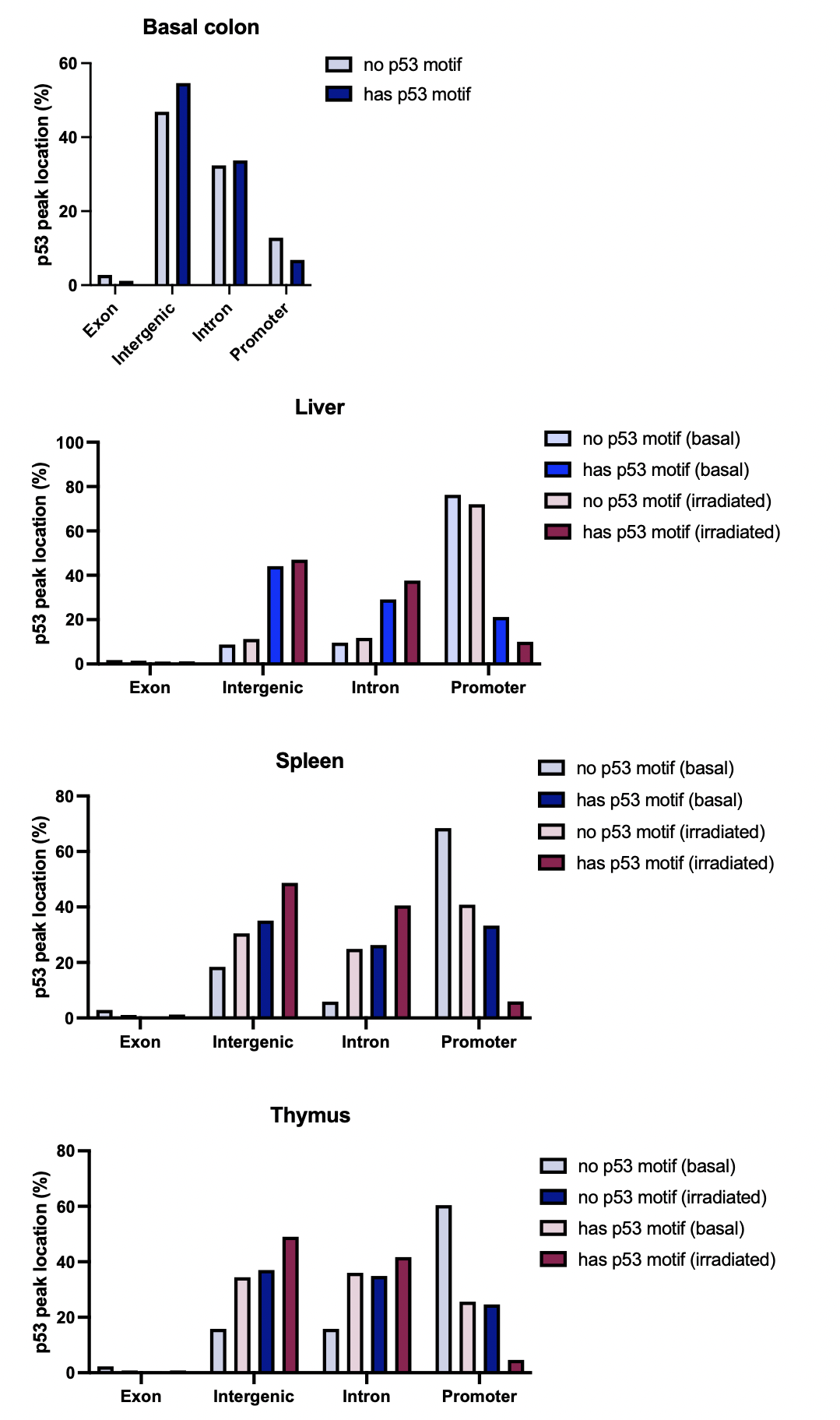

**Supplementary Figure 13. Location of p53 peaks depending on whether they contain a p53 consensus sequence.**

Location of p53 peaks in basal and irradiated tissues. p53 peaks are further divided between whether a consensus binding sequence is present.

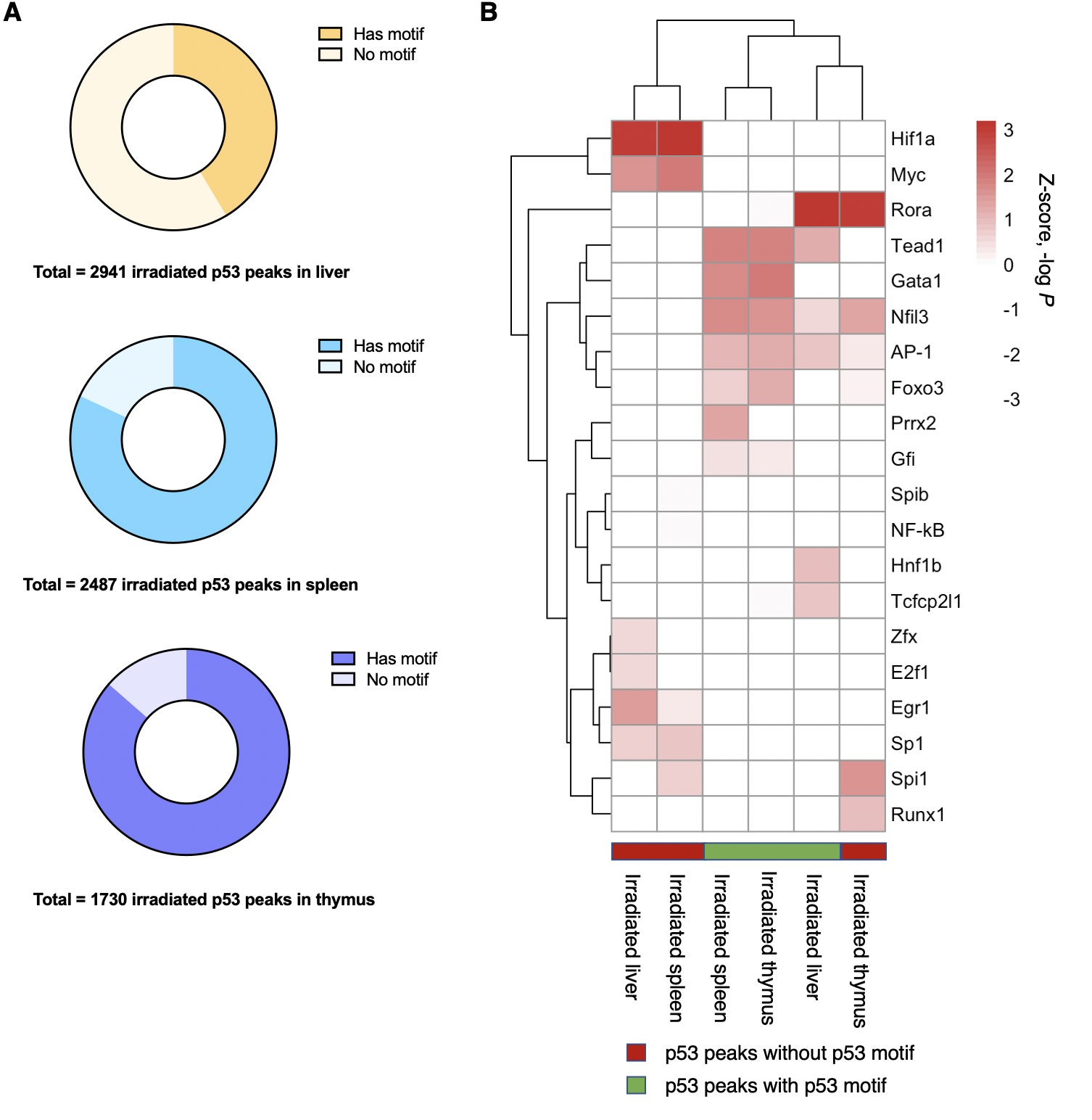

**Supplementary Figure 14. Transcription factor motif analysis of irradiated p53 peaks depending on whether they contain a p53 consensus sequence.**

(A) Pie chart of irradiated p53 peaks depending on whether a p53 consensus binding sequence is present. (B) Heatmap and unsupervised hierarchical clustering of transcription factor motifs (by significance of motif enrichment) among irradiated tissues. Columns are further divided by whether a p53 consensus binding sequence is present. Only expressed transcription factors are included—a non-expressed transcription factor (TPM < 1) is assigned a *P* = 1.

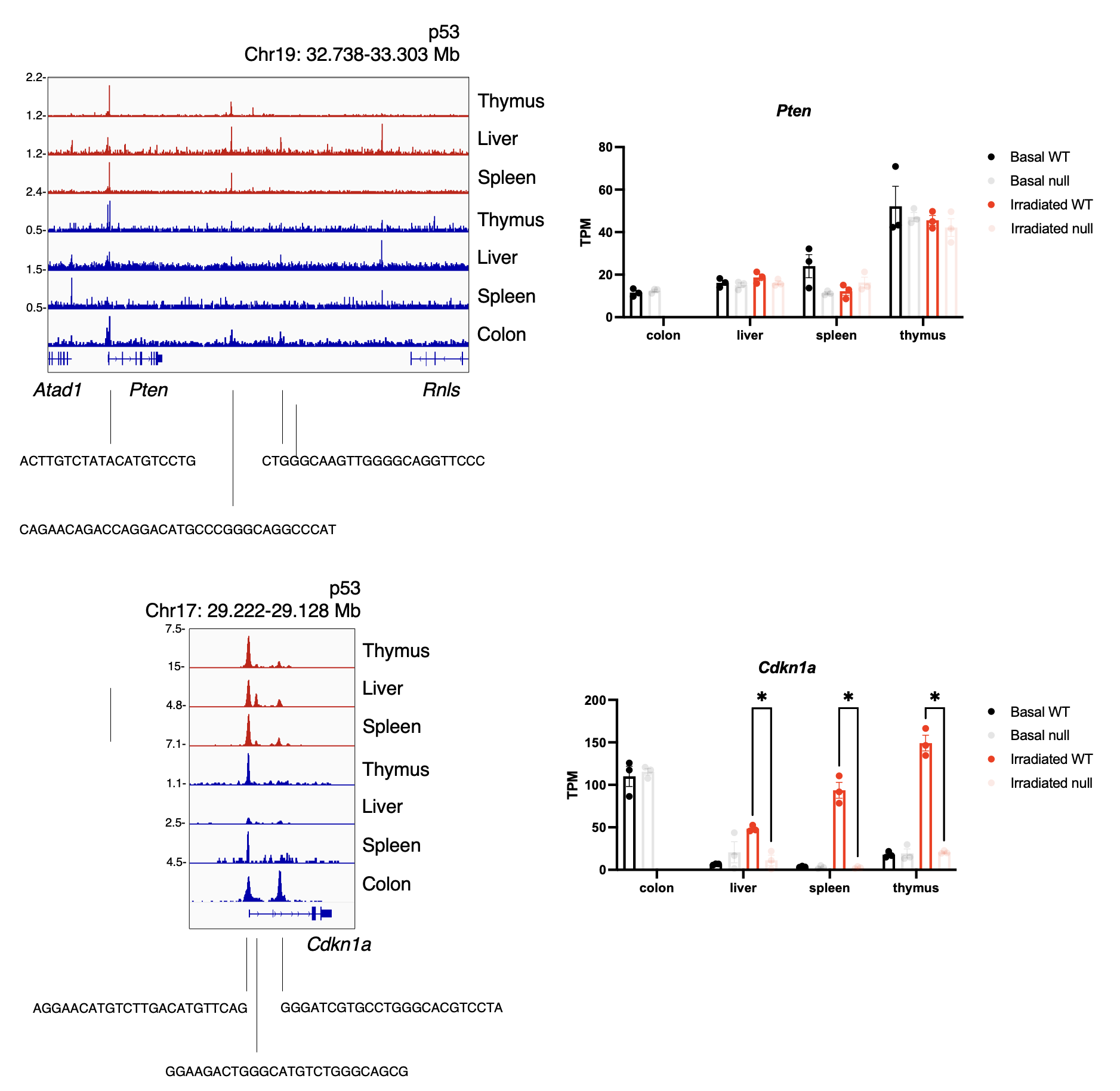

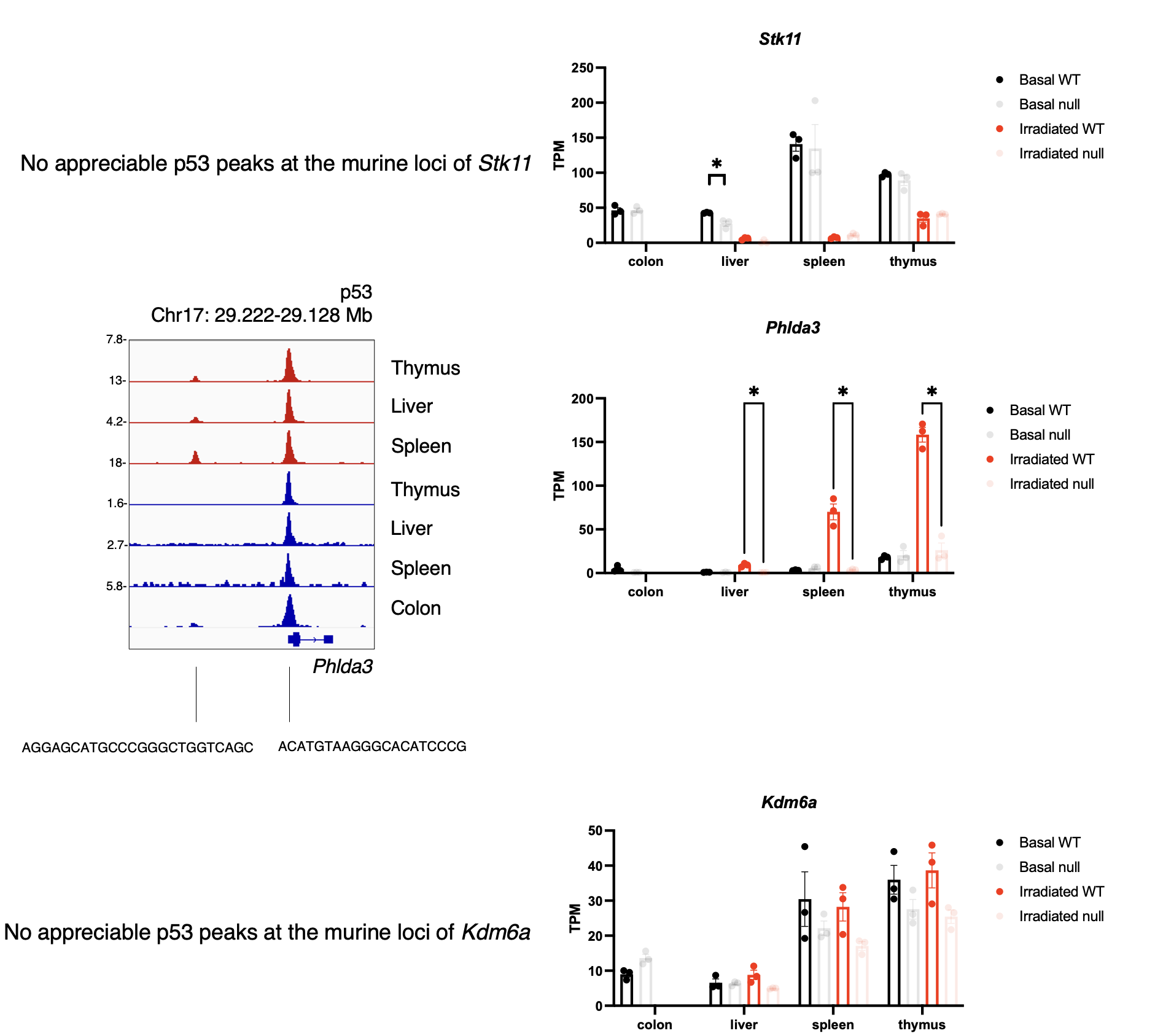

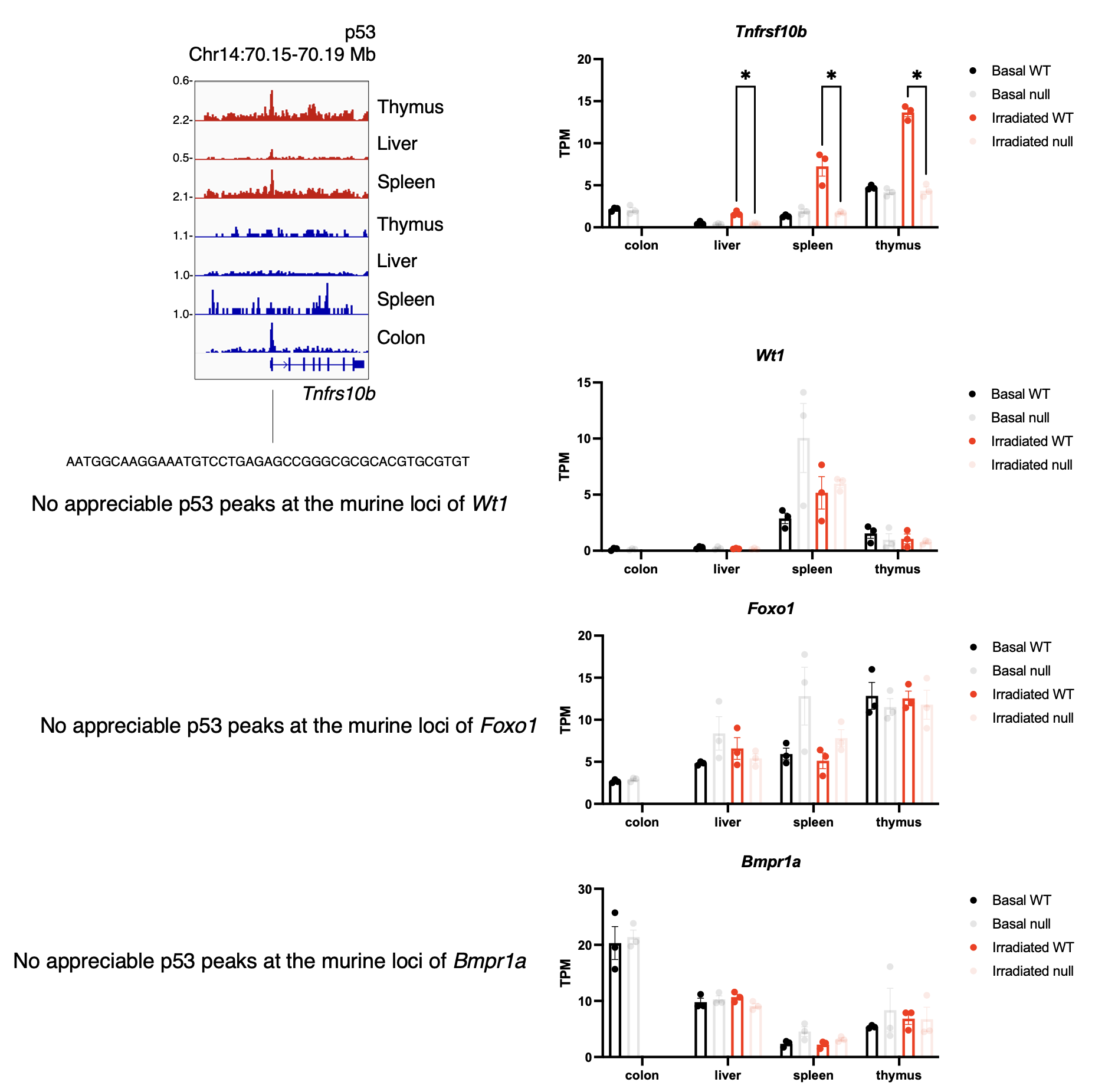

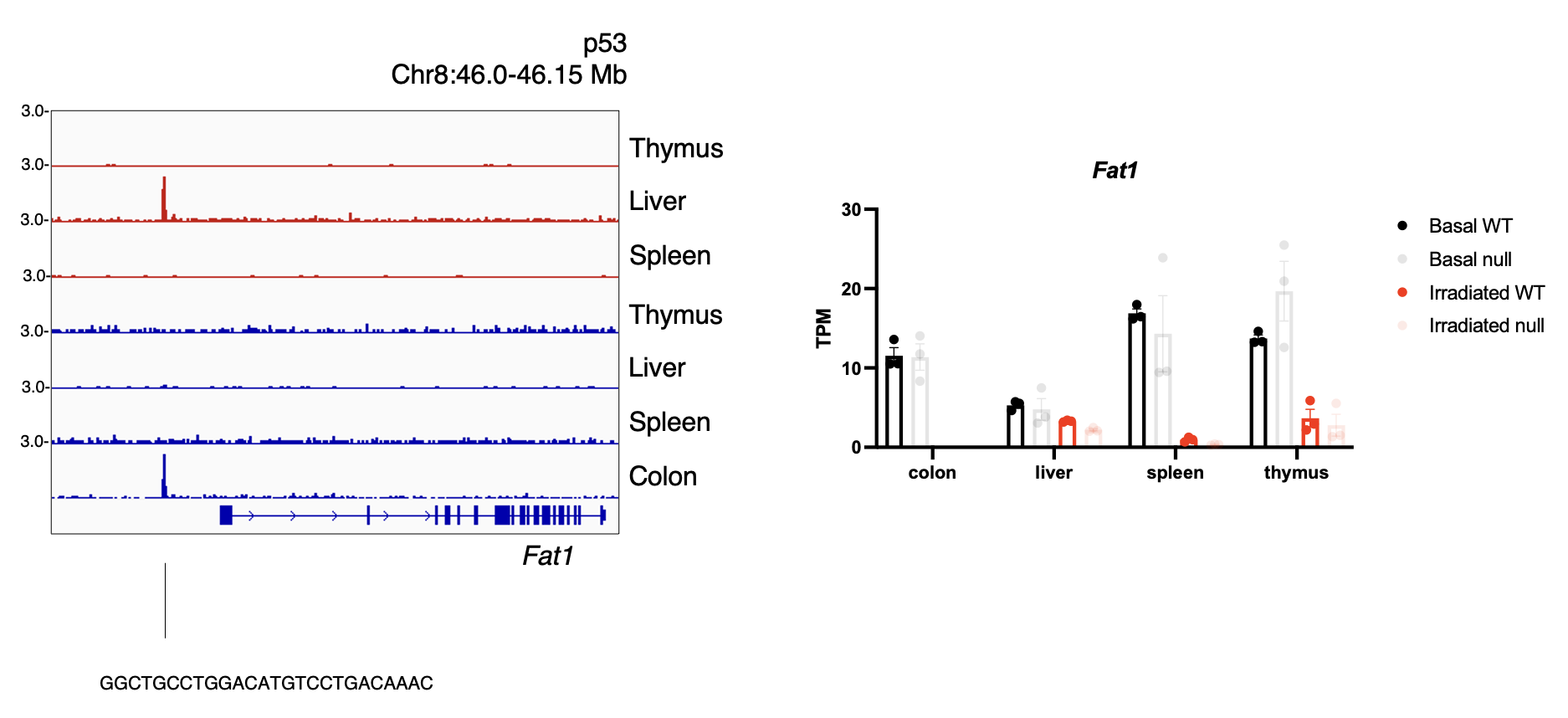
